# Large-Scale Evaluation of Spatial Metabolomics Protocols and Technologies

**DOI:** 10.1101/2024.01.29.577354

**Authors:** Veronika Saharuka, Lucas M. Vieira, Lachlan Stuart, Måns Ekelöf, Martijn R. Molenaar, Alberto Bailoni, Katja Ovchinnikova, Jens Soltwisch, Tobias Bausbacher, Dennis Jakob, Mary King, Max A. Müller, Janina Oetjen, Crystal Pace, Fernanda E. Pinto, Nicole Strittmatter, Dusan Velickovic, Bernhard Spengler, David C. Muddiman, Manuel Liebeke, Christian Janfelt, Richard Goodwin, Livia S. Eberlin, Christopher R. Anderton, Carsten Hopf, Klaus Dreisewerd, Theodore Alexandrov

## Abstract

Spatial metabolomics using imaging mass spectrometry (MS) enables untargeted and label-free metabolite mapping in biological samples. Despite the range of available imaging MS protocols and technologies, our understanding of metabolite detection under specific conditions is limited due to sparse empirical data and predictive theories. Consequently, challenges persist in designing new experiments, and accurately annotating and interpreting data. In this study, we systematically measured the detectability of 172 biologically-relevant metabolites across common imaging MS protocols using custom reference samples. We evaluated 24 MALDI-imaging MS protocols for untargeted metabolomics, and demonstrated the applicability of our findings to complex biological samples through comparison with animal tissue data. We showcased the potential for extending our results to further analytes by predicting metabolite detectability based on molecular properties. Additionally, our interlaboratory comparison of 10 imaging MS technologies, including MALDI, DESI, and IR-MALDESI, showed extensive metabolite coverage and comparable results, underscoring the broad applicability of our findings within the imaging MS community. We share our results and data through a new interactive web application integrated with METASPACE. This resource offers an extensive catalogue of detectable metabolite ions, facilitating protocol selection, supporting data annotation, and benefiting future untargeted spatial metabolomics studies.

## Introduction

Spatial metabolomics is an emerging field focused on detecting metabolites in tissues and cells in their native spatial context, offering valuable insights in biology, medicine, and pharmacology^1^. To comprehensively map metabolic phenotypes in a sample, a wide range of metabolites must be measured in an untargeted manner. This comprehensive metabolite profiling is vital for enabling complex applications, such as discovering novel metabolic interactions^2^, understanding metabolic reprogramming in disease^3^, or analysing metabolic states at the single-cell level^4^. Among the technologies used for spatial metabolomics, matrix-assisted laser desorption/ionisation mass spectrometry (MALDI-MS) imaging is recognised as a key tool.^5^ However, the untargeted analysis of metabolites using MALDI-MS presents challenges related to two factors: the ability to detect diverse metabolites, and confidently annotating the resulting data. Firstly, analyte detectability varies depending on the specific protocol used, including the ionisation mode, organic solvent, and the MALDI matrix. Although numerous matrices have been described, there is no universal protocol optimal for all metabolites due to their high chemical diversity. The lack of comprehensive empirical data on metabolite detectability in MALDI-MS, coupled with the absence of quantitative ion formation theories, hinders our ability to predict the most suitable protocol for specific metabolites. Secondly, without detailed knowledge of the range of metabolites detectable with a given protocol, there is a risk of ion misannotation when relying solely on full scan (MS1) data predominantly used in imaging MS. Similar challenges exist across other imaging MS technologies like desorption electrospray ionisation (DESI) and infrared laser matrix-assisted laser desorption electrospray ionisation (IR-MALDESI). The limited understanding of the disparity in the results between different protocols and technologies makes it difficult to establish result reproducibility and compare outcomes across studies. Addressing these challenges is crucial for enhancing experiment design, reducing uncertainty in data interpretation, and advancing untargeted spatial metabolomics research effectively.

Efforts have been made in the past to tackle similar challenges, including cataloguing analytes detected by various MALDI matrices, including lipids^6^ and small molecules^7^. However, comprehensive information on metabolite detectability remains limited, with many novel matrices^8^ yet to be evaluated for spatial metabolomics. Additionally, studies attempting to explain ion yields in terms of matrix-analyte chemistry have not yet found consistent relationships for predicting analyte detectability.^9–12^ Empirical studies have measured the detectability of lipid classes using different MALDI-MS protocols in tissue sections and homogenates spiked with chemical standards.^13,14^ However, these studies have limited scalability to more diverse analyte classes and may not generalise to other sample types, as ion yields can vary depending on the tissue used.^15^

In this study, we systematically measured the detectability of metabolites representative of biological samples by common imaging MS technologies, focusing primarily on MALDI-MS. A procedure was established to produce a reference sample comprising 172 chemical standards. A computational pipeline was developed to process imaging MS data from the reference samples, allowing us to compare 24 MALDI-MS protocols and determine their suitability for untargeted spatial metabolomics. Chemical standards were used to avoid ion suppression effects typically present in complex biological samples and determine metabolite detectability under ideal conditions. We demonstrated the relevance of our results to biological sample imaging by comparison with metabolite detectabilities in animal tissues for selected protocols. Additionally, an interlaboratory survey involving 10 imaging MS technologies provided insights into metabolite detection within the wider imaging MS community. The results showed that an extensive metabolite coverage is achievable and comparable across different technologies, highlighting that untargeted spatial metabolomics is accessible on a wide range of instruments and that the results presented in this work are informative beyond MALDI-MS. To facilitate the design of future untargeted metabolite experiments by imaging MS, the results and data were made openly accessible through an interactive web resource integrated into the METASPACE online platform. This community resource provides the information on detectability and intensities of individual metabolite ion species, helping to select appropriate protocols and understand their limitations, thereby facilitating data annotation in untargeted metabolite imaging. Furthermore, our methodology and the reference samples could be used to benchmark other imaging MS protocols and instrumentation, as well as assess the detectability of different sets of analytes.

## Results and Discussion

### Workflow to measure analyte detectability and intensity by imaging MS

Our study aimed to evaluate the effectiveness of imaging MS protocols and technologies for untargeted spatial metabolomics experiments. To achieve this, we developed an experimental and computational procedure to measure the detectability and intensities of up to 180 molecules simultaneously in a controlled manner (**Fig. 1b**).

**Figure 1.**
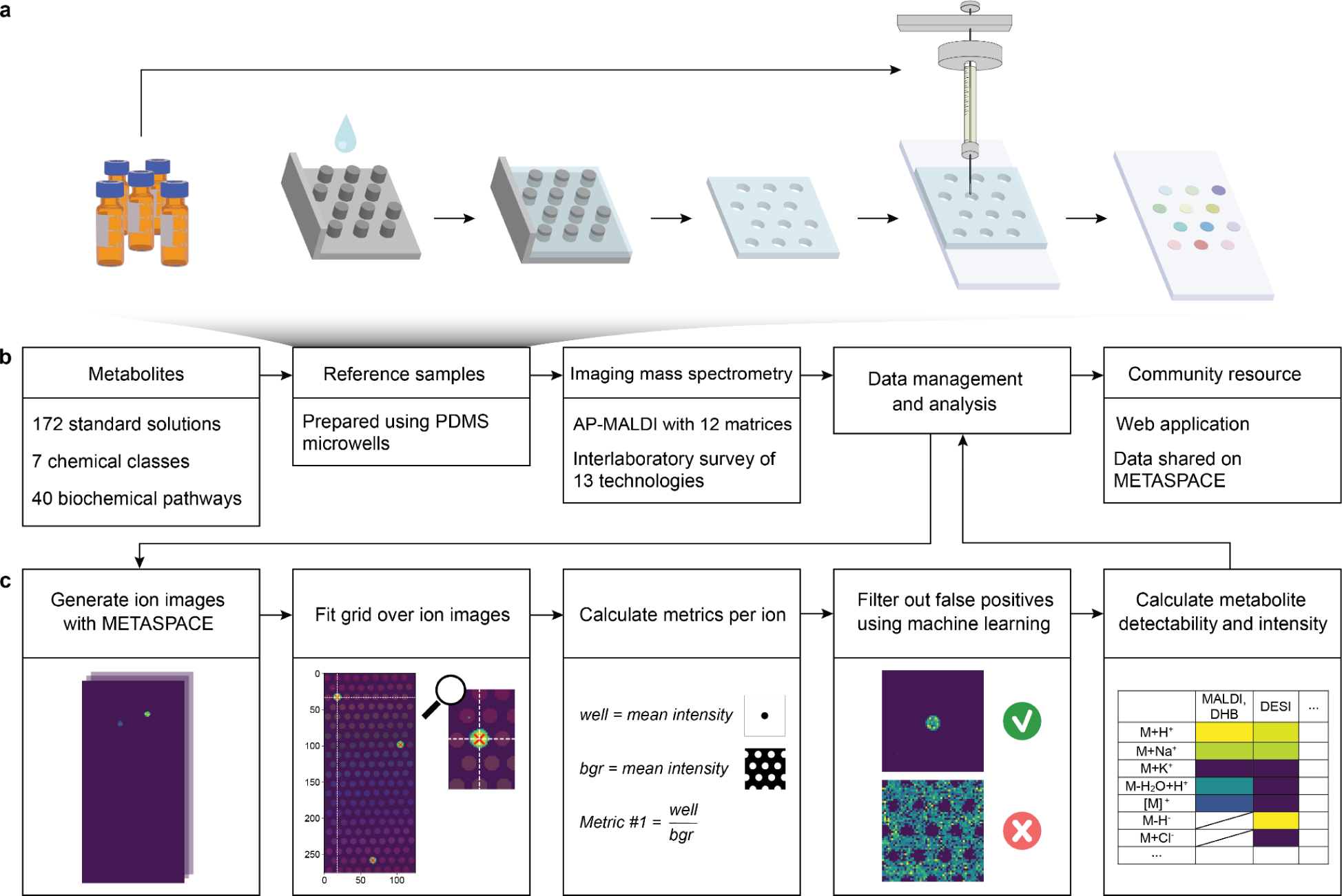
Experimental and computational workflow to measure detectability and intensities of 172 biologically-relevant metabolites using reference samples. ***a****, Schematic representation of the experimental protocol used to prepare the reference samples. An automatic compound dispenser was used to deposit solutions of individual chemical standards onto glass slides, supported by a removable polymer template*. ***b****, Experimental and computational workflow for the reference sample analysis*. ***c****, Schematic representation of the computational pipeline used for the analysis of imaging MS data. m/z channels were queried against a custom database, converted to ion images, and filtered to remove false-positive annotations. Then, the detectabilities and intensities of selected ions were calculated*.

First, we created a reference sample compatible with common soft ionisation imaging MS instruments (**Fig. 1a**). Individual solutions of chemical standards were deposited on a glass slide in a regular pattern, using an automatic thin-layer chromatography (TLC) sample dispenser. To prevent cross-contamination, we used a custom-made removable polymer template that formed leak-tight microwells when placed on the glass slide. This yielded a spatially and chemically defined array of 180 analyte spots, each of 1 mm in diameter, covering a total area of approximately 800 mm^2^. In total, we prepared 150 reference sample replicates.

To analyse the imaging MS data from reference samples, we developed a computational pipeline that quantified the detectability and intensity of the anticipated analyte ions in each spot of the array (**Fig. 1c**). For every metabolite, we examined up to 70 theoretical ions, consisting of all permutations of an adduct (+H^+^, +Na^+^, +K^+^, [M]^+^ in positive ionisation mode, and -H^+^, +Cl^-^, [M]^-^ in negative ionisation mode), and up to one commonly observed neutral loss (water, carbon dioxide, ammonia, dihydrogen, phosphate) or neutral gain (dihydrogen or respective MALDI matrix). All ion images were generated using the *custom database* functionality of METASPACE and resulted in approximately 4000 images per reference sample. Since some theoretical ions were not chemically plausible or isobaric with matrix peaks, some generated ion images were suspected to be false-positive annotations and were excluded from analysis in the following way. First, we used a grid-fitting algorithm to identify the coordinates of the metabolite spots in each ion image. For every metabolite ion in the corresponding spot, several metrics, such as the intensity ratio between the spot area and the surrounding background, were computed. Subsequently, these metrics were used to train a machine-learning classification model to filter out false positive annotations.

As a result, our approach generates a table with identities and intensities of the ions detected in each spot of the array. Using our pipeline, we were able to perform the analysis of tens of thousands of ion images from multiple data sets, providing the throughput necessary for a comprehensive comparison of metabolite detectability and intensities of various MS imaging protocols and technologies.

### Reference metabolites are representative of a typical metabolome

To enable the evaluation of untargeted spatial metabolomics methods, a reference sample was created using 172 chemical standards that represent the diversity of primary metabolites found in mammals, bacteria and yeast. Comprehensive coverage of various biological functions was ensured by selecting metabolites from a wide range of biochemical pathways, as illustrated in the KEGG metabolic pathways map (**Fig. 2a**, **SI Fig. 4**). We included multiple intermediates per pathway, covering the metabolism of amino acids, carbohydrates, lipids, nucleotides, vitamins and cofactors (**SI Fig. 5, SI Table 9**). Additionally, we selected the metabolites from a wide range of chemical classes using a two-level classification (“class” and “subclass”) derived from ClassyFire^24^ (**SI Fig. 6, SI Table 10**). The structural diversity of the reference metabolites was compared to that of the metabolome used in the genome-scale metabolic mouse model BiGG iMM1865^21^. For this, we converted the molecular structure of each metabolite to a binary extended-connectivity fingerprint^25^. Then, we mapped all metabolites from BiGG iMM1865 together with our 172 metabolites into a chemical space by applying Principal Component Analysis to their fingerprints (**Fig. 2b**). We found that the reference metabolites span the entire chemical space occupied by the BiGG iMM1865 metabolome. Metabolites assume a wide range of concentrations in living cells. To ensure that we included the most abundant metabolites in our reference sample, we compared our selection to the data set of absolute abundances of over 100 metabolites in *Escherichia coli* (data not available for cells of other organisms).^26^ Our reference sample incorporates a majority of the most abundant metabolites, including 15 of the top 20. (**SI Fig. 7**). In summary, our carefully selected set of 172 reference metabolites is representative of the biochemical, chemical, and structural diversity found in a typical metabolome.

**Figure 2.**
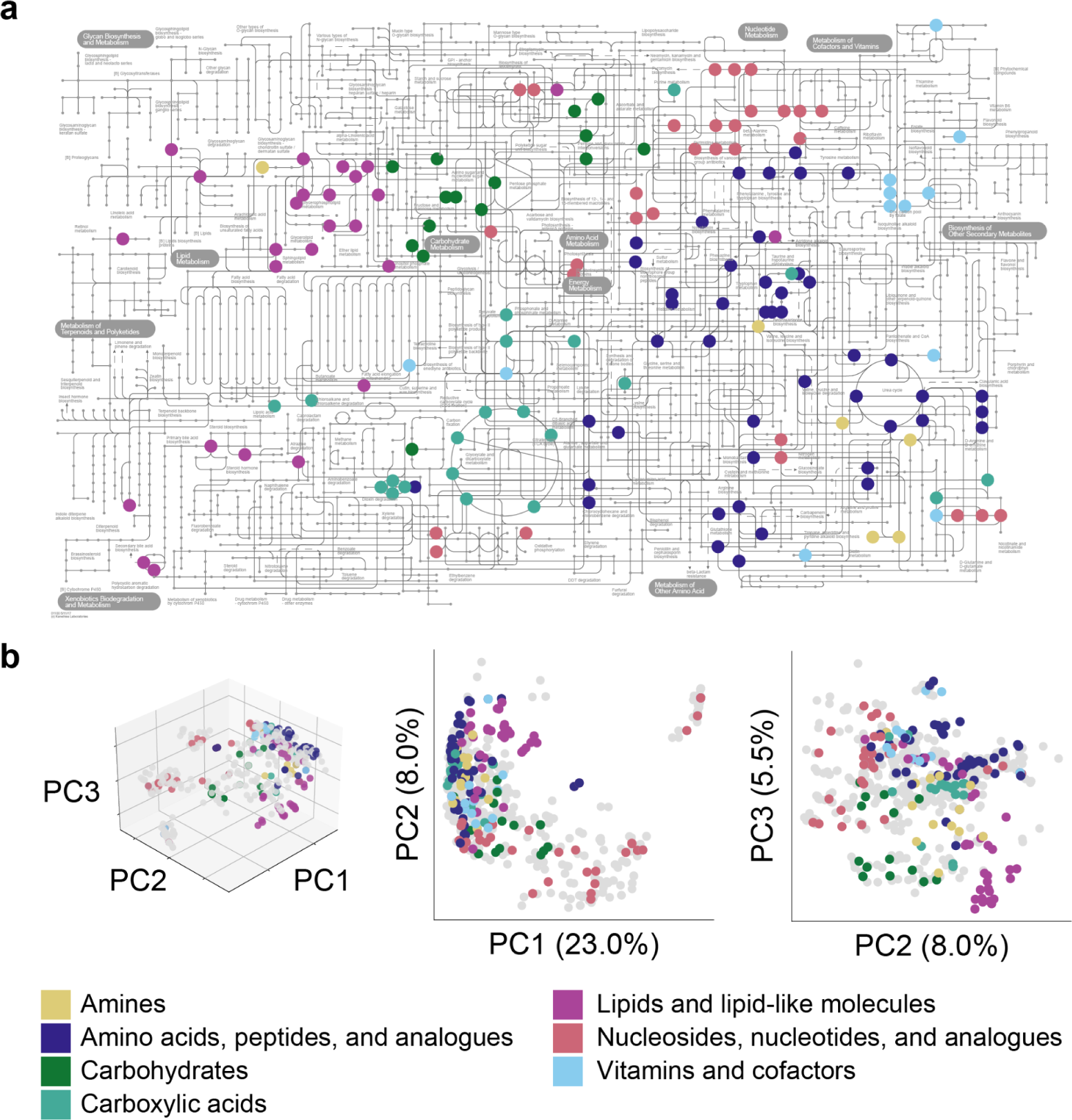
Function, structure and natural abundance of reference metabolites. ***a****, KEGG map of common metabolic pathways shows the distribution of reference metabolites across different biochemical pathways. The nodes represent metabolites and the lines represent reactions. Here and in panel b reference metabolites are coloured according to the chemical class*. ***b****, Structural similarities between 172 reference metabolites (coloured) and metabolites from the BiGG iMM1865 genome-scale mouse model (in grey). The chemical space is visualised using Principal Component Analysis of metabolite extended-connectivity fingerprints*.

### Metabolite detectability and intensities for 24 different MALDI-MS protocols

Using our reference samples, 24 MALDI-MS protocols for untargeted spatial metabolomics were compared. We selected 12 MALDI matrices known for detecting low molecular weight compounds, including some commonly used in over 7000 METASPACE public datasets (**SI Table 4**, https://metaspace2020.eu/datasets/summary, **SI Fig. 13**). The matrices were sprayed onto the reference samples with a consistent matrix-to-analyte ratio, and the samples were imaged using atmospheric pressure (AP-) MALDI-MS in both positive and negative ionisation modes. We used matrices commonly deployed in positive ion mode DHB, CHCA, DHAP, ClCCA, and CMBT, matrices commonly deployed in negative ion mode 9AA, NEDC, and MAPS, as well as dual-mode matrices DAN, NOR, pNA, and PNDI.

Our computational pipeline determined the detectability of each metabolite ion with each protocol (**see SI files**). A metabolite was considered to be detected if any of its intact ions were detected (excluding neutral losses/gains). The majority of metabolites were detected by at least one MALDI matrix (91% in positive mode, 82% in negative mode). Among the 11 undetected metabolites, some could be detected after a neutral loss (cholesterol, cholesteryl acetate), some were obscured by isobaric background peaks (pyruvic, lactic, butyric and acetoacetic acids, dihydroxyacetone), or were only detectable from concentrated solutions (arachidonic acid, phosphoserine, carbamoyl phosphate). Indole was undetectable, likely due to its volatility. In each polarity, half of the metabolites were detected by at least half of the protocols (**Fig. 3a, SI Fig. 8**). In positive polarity, most metabolites in the considered chemical classes were detected by at least one protocol, while in negative polarity, analytes that are also known to poorly ionise as deprotonated species in liquid-chromatography MS (amines, glycerolipids, and positively charged metabolites) were not detected (**Fig. 3b**, **SI Fig. 9**). The number of metabolites detected per protocol ranged from 16 to 154 (**Fig. 3c**). DHB, CHCA, and ClCCA performed best in positive mode, while 9AA resulted in the highest number of detections in negative mode. The dual-mode matrices (DAN, NOR, and pNA) achieved good coverage in both polarities, making them attractive for applications in which the sample amounts are limited but analysis in both ionisation modes is desired. The NEDC, MAPS, and PNDI protocols detected fewer than 50 metabolites each, indicating their limited suitability for untargeted spatial metabolomics at the matrix-to-analyte ratio used. However, given their reported advantages^27–29^, further research is needed to determine the optimal protocols and applications.

**Figure 3.**
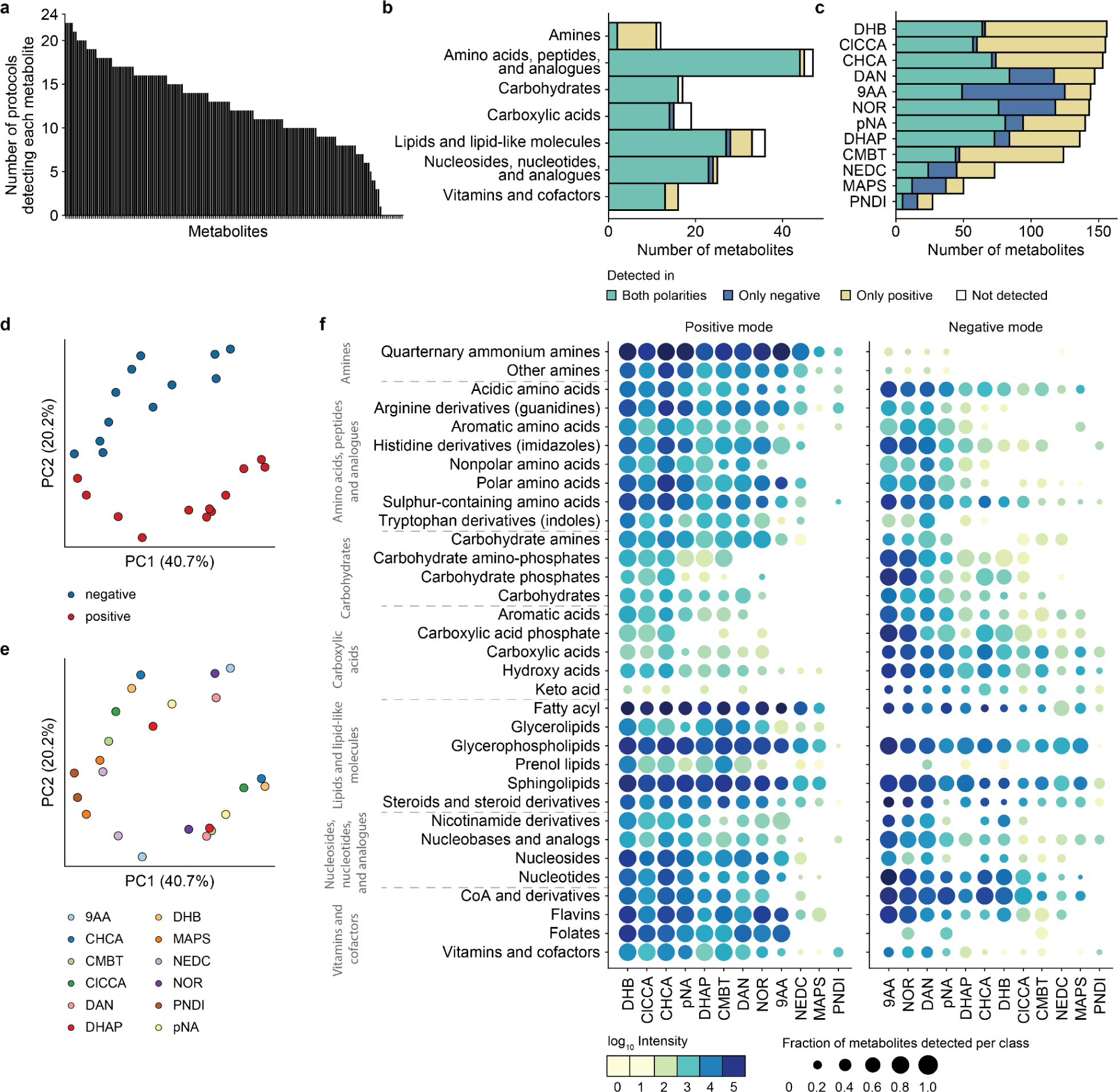
Metabolite detectability by AP-MALDI imaging mass spectrometry. ***a****, Number of MALDI-MS protocols detecting each reference metabolite. One bar represents one metabolite*. ***b****, Number of reference metabolites detected in each chemical class by at least one protocol, coloured by polarity*. ***c****, Numbers of reference metabolites detected per MALDI matrix, coloured by polarity*. ***d-e****, PCA comparing MALDI-MS protocols, coloured by polarity (d) and matrix (e), based on the detectabilities and intensities of reference metabolites*. ***f****, Detection of metabolites in chemical classes and subclasses by different MALDI-MS protocols. Dot size shows the fraction of metabolites detected in a given class among those considered; colour shows the average intensity among the detected ions*.

We compared the protocols based on the detectabilities and intensities of reference metabolites (**Fig. 3d-e**). As expected, polarity significantly drove orthogonal differences between protocols, as seen by the separation of data sets acquired with the same matrix along PC2. Within each polarity, the protocols varied based on the matrix used (along PC1). Some groups of matrices notably exhibited similar performance, for instance, DHB, CHCA, and ClCCA in positive mode.

In a detailed comparison of chemical classes (**Fig. 3f**), most metabolites were detectable in both polarities but many showed a preferred ionisation mode. Amines were predominantly detected in positive mode but not in negative mode, regardless of the matrix. Intact carbohydrate ions were better detected in negative ionisation mode, especially with the 9AA and NOR matrices. Carboxylic acids had higher intensities in negative mode as carboxylate ions, however were also observed in positive mode as sodiated ions. Amino acids produced intense ions in both polarities, as expected from their ability to form zwitterions, with the differences driven by the chemical properties of the side chain. For example, arginine and creatine with guanidine functional groups were better detected in positive mode, while the acidic amino acids were better detected in negative mode. Among lipids, ions from a wider variety of subclasses were detected in positive mode, with glycerolipids and prenol lipids being almost exclusively detected in this polarity. The detectability of lipid metabolites in glycerolipid and sphingolipid subclasses is often influenced by the lipid headgroup. Members of these classes were found to be detected with high coverage and ion intensities by most MALDI matrices. In particular, acidic or carbohydrate groups led to higher ion intensities in negative polarity, for example, for cholic acid, phosphoinositol or glycosyl cholesterol. Conversely, lipids containing quaternary amines, such as phosphatidylcholine and acylcarnitine, were preferentially detected in positive mode. Nucleotides and derivatives showed similar intensities in both polarities, except for those with phosphate groups (e.g. UDP-glucose, coenzyme A, FAD), which had higher intensities in negative mode.

Some chemical classes showed comparable detection across most tested matrices in each polarity, such as quaternary ammonium amines, glycerophospholipids, and sphingolipids in positive mode. However, optimal matrix-polarity combinations were required for other chemical classes to achieve high ion intensities. For example, carbohydrates in negative polarity were best detected with 9AA and NOR matrices. Among these, carbohydrate phosphates including fructose-, glucose-, and glycerol phosphates showed poor detection with other matrices. A follow-up experiment confirmed that fructose-1,6-bisphosphate and glucose-6-phosphate could not be detected in a mixture with DAN, but yielded strong deprotonated peaks with 9AA. Our results suggest that the "top 3" matrices in each polarity (DHB, ClCCA, and CHCA in positive mode; 9AA, NOR, and DAN in negative mode) provide better metabolite detectability and ion yield compared to other protocols, making them best-suited matrices for untargeted spatial metabolomics. It should be noted that other factors such as matrix vacuum stability, crystal size, optical absorption at the laser wavelength used, and chemical safety should also be considered in matrix selection.

Finally, we grouped our results by biochemical pathway, which can be instructive for biological applications requiring targeted analysis of specific metabolic aspects (**SI Fig. 10**). Note that our study did not cover all intermediates in every pathway. We found that positive mode protocols effectively detected metabolites of amino acid metabolism (e.g. urea cycle, polyamine biosynthesis and creatine metabolism), one-carbon cycle (including folate and methionine cycle, and vitamins riboflavin and pyridoxine), and intermediates of carnitine biosynthesis and fatty acid oxidation. Negative mode protocols provided superior coverage of carbohydrate metabolism (e.g. glycolysis, pentose phosphate pathway, and the citric acid cycle), the nucleotide sugars of the hexosamine pathway, and intermediates of nucleotide biosynthesis. Negative mode protocols were also better suited for detecting RedOx intermediates (e.g. NAD, NADH, FMN, and FAD).

Overall, our results provide insight into the detectability of metabolites by 24 MALDI-MS imaging protocols, emphasise the importance of matrix and polarity choice for obtaining orthogonal or comparable results, and serve as a guide for both untargeted spatial metabolomics and targeted analysis of metabolic pathways or chemical classes.

### Reference metabolite detectability is relevant for spatial metabolomics of biological tissues

The analysis of pure chemical standards offers precise control over sample composition, prevents signal suppression by other analytes, and simplifies result interpretation. However, it primarily reflects metabolite detectability in idealised conditions, which may differ from untargeted spatial metabolomics experiments with biological samples. To assess the relevance of our results for biological applications, we compared metabolite detectability between reference samples and biological tissue data sets acquired on the same instrument. We selected 38 mouse tissue data sets from the public knowledge base METASPACE, analysed using two commonly used protocols: DHB in positive mode and DAN in negative mode. This included healthy tissues (brain, lungs, liver, intestine), lung tumour, and inflamed liver tissue. For each tissue data set, we generated ion images corresponding to metabolites present in the reference samples. We manually filtered out false-positive annotations, retaining only the ions detected within the tissue sections. Then, we compared the fractions of detected metabolites in the tissue and reference sample data sets for each chemical class (**Fig. 4a-b, SI Fig. 11**). Based on the assumption that the reference sample represents metabolite detectability under ideal conditions, we hypothesised that all ions detectable in biological tissues would also be detected in the reference samples, and that ions undetectable in the reference samples would not be found in biological tissues. Our results supported this hypothesis, with 95% of DHB-detected and 78% of DAN-detected metabolites in biological tissues also found in corresponding reference samples. Most chemical classes had higher detectability in reference samples than in biological tissues, which is expected given that some metabolites may not be present in the tissues at concentrations exceeding the limit of detection. Notably, the subclasses "Flavins" and "CoA and derivatives" were exclusively present in the reference sample in both protocols. In a small subset of cases (13% with DHB, 24% with DAN), metabolites were found in biological tissues but not in reference samples, e.g. in the subclasses “Keto acids” and “Glycerolipids” in negative mode DAN protocol. The discrepancy in these cases may be attributed to false positive metabolite annotation in biological tissues, as the use of MS1-based metabolite annotation in complex samples can result in misidentification due to naturally occurring isomers, isobars or fragments. Additional methods (such as LC-MS/MS, ion mobility separation, or comparing annotation scores of isobars) may be necessary to confirm metabolite identities in these cases. Our analysis demonstrates an agreement between metabolite detectability in reference samples and animal tissue sections, highlighting the informative and relevant nature of our findings for untargeted spatial metabolomics of tissues using MALDI-MS.

**Figure 4.**
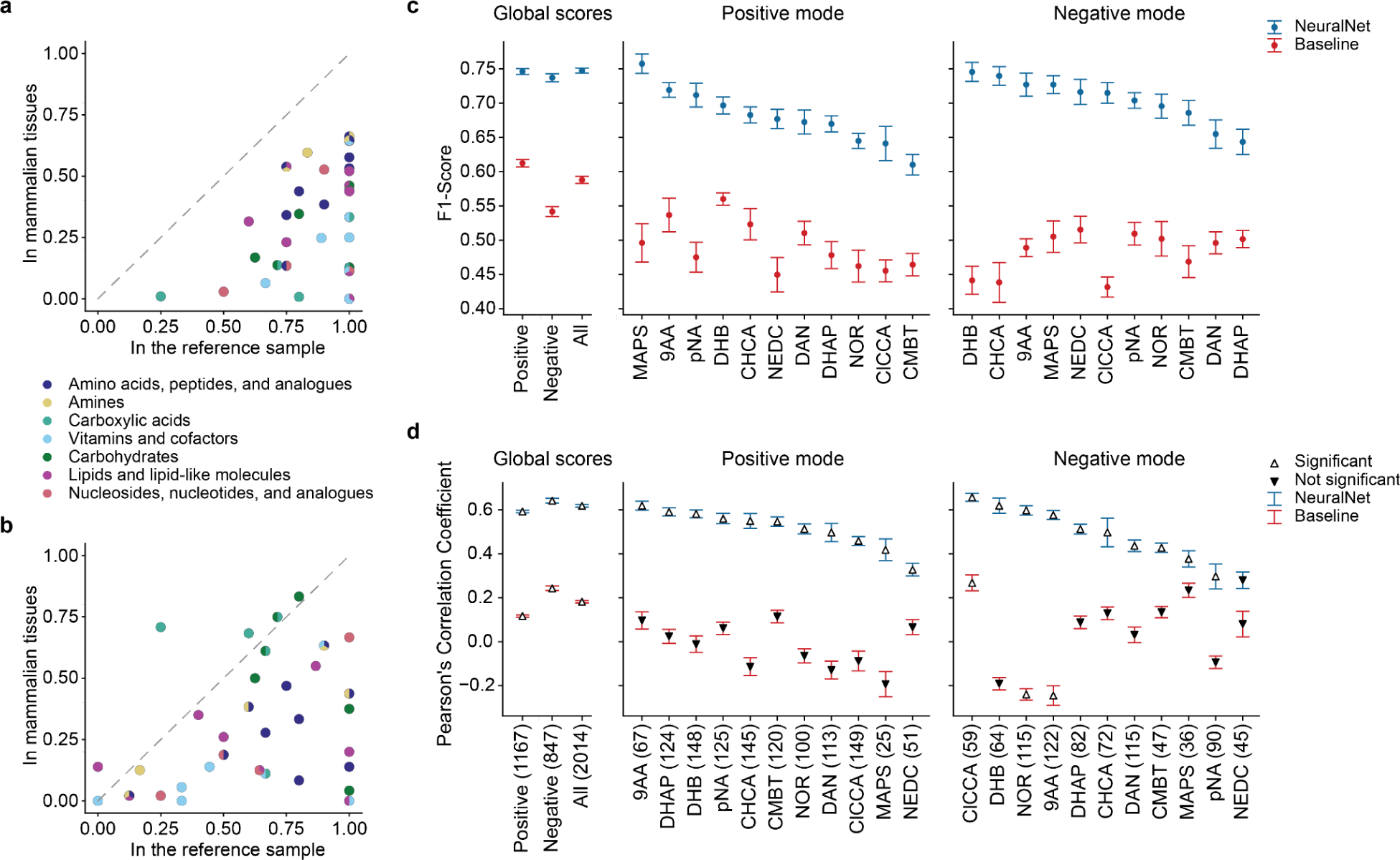
Applicability of reference sample results obtained using AP-MALDI to other samples and metabolites. ***a****-**b**, Fraction of metabolites detected per subclass in the reference sample, and in biological tissues (average across multiple tissue data sets), using DHB positive mode protocol (a) and DAN negative mode protocol (b). Each dot represents a chemical subclass, coloured by chemical class*. ***c****, Accuracy of classifiers predicting metabolite detectability for a specific MALDI matrix and polarity. Left to right: global F1-scores for predictions from all classifiers; scores for each individual classifier*. ***c-d****, Mean values and confidence intervals (+-standard deviation) after repeating the training process 10 times are shown. For the baseline model, the input features were randomly shuffled across all data points*. ***d****, Accuracy of regression models predicting intensities of metabolites. Left to right: global Pearson’s correlation scores for predictions from all models; scores for each individual model. The number in brackets shows the number of detected metabolites considered in each model. Triangle symbols indicate the significance (p-value<0.05) of the calculated correlation*.

### Machine learning predicts metabolite detectability and intensities

To explore the generalisability of our results to a wider range of metabolites, we used machine learning to predict metabolite detectability and intensities in a two-step procedure. First, we trained classifiers predicting metabolite detectability for every protocol individually. For model training, we used the data obtained from the reference samples, along with metabolite physicochemical properties as input features. The properties were obtained from HMDB and included pKa of the most acidic and most basic sites, polar surface area, polarisability, donor and acceptor count and physiological charge. The classifiers were evaluated using F1-scores, and three summary scores were computed by considering predictions from all models: for the positive polarity, negative polarity and both polarities together (**Fig. 4c**). We achieved high accuracy in predicting metabolite detectabilities, with a global F1-score close to 0.75, outperforming a baseline predictor with an F1-score below 0.6. The baseline predictor was trained on the same data, except that the physicochemical property values were randomly shuffled among all metabolites. Next, we trained regression models to predict intensities of all metabolites classified as detected (**Fig. 4d**). Our approach accurately predicted intensities, achieving a global Pearson correlation above 0.6 compared to below 0.3 for the baseline model. Although regression models were trained with limited data points for individual matrix-polarity combinations (numbers in brackets on the X-axis of **Fig. 4d**), they demonstrated robust predictive capability for the majority of protocols in estimating metabolite intensities. It should be noted that the direct comparison between protocols was only possible because the data was collected on the same instrument with a standardised set of parameters and cannot be extrapolated to other MALDI-MS instruments. To determine the impact of individual physicochemical properties on the regression, we employed SHAP (SHapley Additive exPlanations) analysis (**SI Fig. 12**). Higher polarisability values were associated with higher predicted ion intensities in the majority of models. In positive mode, ions with stronger basic sites were predicted to have higher intensities, likely because stronger bases can form more stable, and thus more abundant cations. In negative mode, ions with lower physiological charge were predicted to have higher intensities, a plausible explanation for which is that physiological charge reflects the stability of metabolite anions.

Our proof-of-concept investigation showed that physicochemical properties can be predictive of metabolite detectability and intensities using data from pure chemical standards. Future studies could expand on these findings. We anticipate that using data sets including more analytes and additional molecular properties would result in better predictive models, enabling predictions for metabolites beyond our reference sample and furthering understanding of analyte ionisation mechanisms.

### Interlaboratory survey achieved comparable metabolite detectability across imaging MS technologies

Imaging MS can be performed using a variety of instruments capable of spatial sampling, regardless of ionisation and detection principles. However, the overlap in detected ions across different technologies and the intercomparability of results in the context of untargeted spatial metabolomics remain unclear. To address this, we conducted an interlaboratory survey involving 10 expert laboratories and compared the metabolite detection capabilities of 13 instruments. We selected instruments representing the diversity and state-of-the-art in imaging MS, focusing on soft ionisation technologies capable of detecting intact metabolite ions across a wide mass range. (**SI Table 7**). Further requirements were the ability to generate complete images of the reference sample compatible with METASPACE annotation (centroided spectra with a mass accuracy better than 10 ppm for all peaks and resolving power of at least 50,000 at *m/z* 200). Our selection included major technologies represented in public METASPACE data sets (**SI Fig. 13**): MALDI, DESI, and IR-MALDESI sources, with Orbitrap, FTICR, or QTOF analysers. We aimed to provide an overview of potential variations in spatial metabolomics results when analysing the same sample with different technologies, provided that each technology is operated by an experienced user with the most suitable protocol. Thus, all participants were asked to choose protocols best suited for wide-range spatial metabolomics on their system, and acquire images of the reference samples in positive and negative ionisation modes. The resulting data was submitted to METASPACE and processed through our computational pipeline, as previously explained.

Due to inherent differences in technologies, such as variations in the amount of material sampled and ionised, and differences in mass analysers and acquisition settings, direct comparison of raw ion abundances was not feasible. Instead, technologies were evaluated based on their ability to detect individual metabolites. Our analysis showed no systematic differences in metabolite coverage between technologies based on ion source, mass analyser, pressure mode, matrix choice or participant, except for DESI data sets in positive polarity, which stood apart from other data (**Fig. 5a-b, SI Fig. 14**) likely due to differences in detection of metabolites in classes such as lipids, carbohydrates, vitamins and cofactors (**Fig. 5c**). However, a cluster of technologies with similar metabolite detectability was identified in positive mode (shown with dashed line), encompassing a diverse set of instruments. Further analysis revealed that this cluster exhibited the most extensive metabolite coverage, including data sets acquired with AP-MALDI-Orbitrap, IR-MALDESI-Orbitrap, MALDI-FTICR, MALDI2-Orbitrap, and MALDI-qTOF protocols (see 7J, 8A, 9A, 12C 4F and 13G in **Fig. 5c**). In negative mode, the individual data sets acquired with DESI-Orbitrap, IR-MALDESI-Orbitrap, and AP-MALDI-Orbitrap achieved best overall metabolite coverage (see 1C, 9E and 7A in **Fig. 5c**). It should be noted that protocol choice alone can significantly impact the results, therefore protocol optimisation for a sample of interest is strongly recommended (e.g. matrix-to-analyte ratio, polarity of the matrix solvent, wetness of the spray, or sublimation conditions may be especially important for complex samples). Overall, most metabolite classes were well-detected by all technologies, and high coverage was achievable by all instruments. Differences in chemical class coverage were noted depending on polarity, consistent with previous findings from AP-MALDI protocol comparison (**Fig. 3**). For example, basic amines in negative ionisation mode and carboxylic acids in positive mode presented challenges for detection across most technologies. Within each polarity, we did not observe direct orthogonality of metabolite detection profiles among technologies. Notably, differences between results from instruments with the highest metabolite coverage were smaller than differences between individual chemical classes. This suggests that the detectability of sufficiently abundant metabolites is primarily determined by their chemistry, and differences in instrumentation alone are unlikely to be the sole cause of contradictory results in similar untargeted spatial metabolomics experiments. Selecting the right technology, however, is important for particular research questions. For example, MALDI2 was shown to have increased sensitivity for specific glycerophospholipids in complex biological samples with low-abundant analytes or notable ion suppression.^30^ Additionally, DESI’s sampling approach allows imaging of bacterial colonies on agar without sample destruction, facilitating subsequent subculturing.

**Figure 5.**
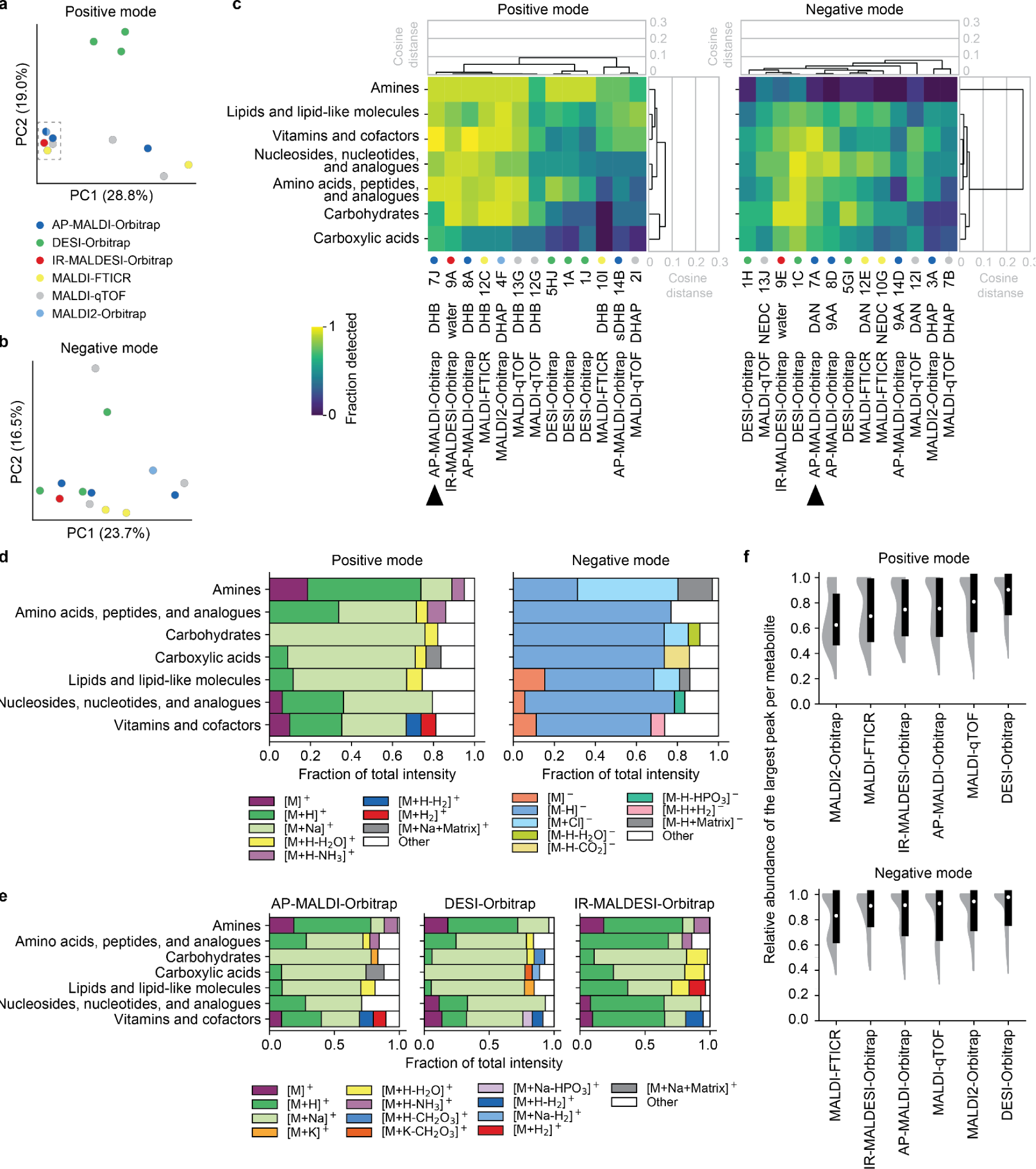
Interlaboratory survey of metabolite detectabilities and signal dilution across 13 imaging MS instruments. ***a-b****, Comparison of imaging MS technologies based on the detectability of reference metabolites in positive and negative polarities*. ***c****, Detectabilities of reference metabolites across chemical classes and technologies. Heatmap represents the fraction of metabolites detected in each chemical class. Data sets generated using the same technology as in* Fig. 3 *are marked with black arrows. Dendrograms show cosine distances between individual classes and technologies. Dots show the same technologies as in panel a*. ***d****, Proportional contribution of different ion types to total detected metabolite signal, averaged per chemical class across all technologies. Contributions below 5% are labelled as "Other"*. ***e****, Contribution of ion types to total detected metabolite signal (as in panel a) for three selected technologies*. ***f****, Relative abundance of the most intense ion peak per metabolite compared to the total signal detected per that metabolite in each considered imaging MS technology. Violin plot and box plot (25th, 50th, and 75th percentiles) depict values for detected reference metabolites*.

Our survey demonstrates that common imaging MS technologies offer extensive and comparable coverage of metabolites in reference samples. This highlights the accessibility of untargeted spatial metabolomics and positions our metabolite detectability results as a valuable resource for the broader imaging MS community.

### Investigation into the formation of different ion types

To accurately interpret untargeted spatial metabolomics data, establishing confidence in ion annotation is crucial. One of the challenges in imaging MS is the division of signal from a single analyte between multiple mass channels, e.g., arising through different ionising adducts or fragmentation reactions. The increased spectral complexity from such “signal dilution” can lead to false positive identifications due to *m/z* overlaps with isomeric or isobaric ions, or missed annotations due to reduced sensitivity for the target analyte ion. Here, we used the data from the interlaboratory survey to estimate the extent of signal dilution, and determine the prevalent ion types across different imaging MS technologies. For this, we considered all permutations of 6 common adducts and 10 neutral losses/gains and compared intensities of detected ions for every reference metabolite.

Our analysis revealed intact metabolite ions produced through protonation, deprotonation, or metal cation adduction were the most abundant ions across all technologies (**Fig. 5d**). In positive mode, sodium adduction [M+Na]+ was the primary ionisation mechanism for classes containing oxygen as the only heteroatom, such as carboxylic acids and carbohydrates, while direct protonation [M+H]+ was more prevalent for nitrogen-containing metabolites like non-quaternary amines and amino acids. Molecular ions without adducts, [M]+ and [M]-, were also detected, some as a result of radical formation, and others due to databases only listing charged formulas (e.g. thiamine). Furthermore, characteristic ions formed by the loss of neutral molecules were observed, particularly in metabolites with good leaving groups. In positive polarity, the most prevalent neutral losses were the loss of water from amino acids, carbohydrates, carboxylic acids, and lipids, and the loss of ammonia from amines and amino acids. In negative polarity, the loss of carbon dioxide from carboxylic acids, and the loss of phosphoric acids from phosphate-containing metabolites were observed. Certain metabolites exhibited a neutral gain of a matrix unit, especially with uncommon matrix-polarity combinations, such as 9AA in positive mode or DHAP in negative mode. However, the intensity of these ions was generally low. Technology-specific differences were also observed (**SI Fig. 15**). For example, AP-MALDI-Orbitrap and DESI-Orbitrap showed similar ion formation patterns, whereas IR-MALDESI-Orbitrap exhibited a higher abundance of protonated cations compared to the total metabolite signal across all chemical classes (**Fig. 5e**). This finding aligns with prior research demonstrating the preferential production of protonated cations by IR-MALDESI.^31^

Overall, signal dilution was more prominent in positive mode across all technologies, primarily due to the split between sodiated and protonated ions. On average, the most abundant ion accounted for 62-90% of total metabolite intensity in positive mode and 83-97% in negative mode. This highlights the potential loss of significant spectral information if only the primary peak is considered (**Fig. 5f**). For a few metabolites, the dilution effect was more severe. For example, over 50% of the adenosine triphosphate signal was split due to fragmentation, cholesterol was predominantly detected as a dehydrated carbocation, and retinoic acid formed multiple abundant ions such as radical cation in positive mode (**SI Fig. 16**). It is important to note that our figures may underestimate the extent of signal dilution, as we only considered common adduct and neutral loss combinations. Additionally, signal dilution observed in biological samples may differ depending on the availability of ionising adducts.^15,32^

In summary, our data helped estimate the extent of signal dilution and identified prevalent ion types across different imaging MS technologies. This provides a basis for improving measurement sensitivity and annotation confidence in untargeted imaging MS analysis. Further research can build upon our findings by exploring additional adducts, neutral losses, and fragments in the data.

### Web application for interactive visualisation of metabolite detectabilities

We developed an interactive web application (http://metaspace2020.eu/detectability) to enhance the accessibility of study results and data (**Fig. 6a**). The application visualises metabolite ion detectabilities and intensities as dot plots or heatmaps. Users can aggregate results by various parameters such as metabolite, chemical class, biochemical pathway, polarity, MALDI matrix, and technology, exploring fractions and average intensities of detected metabolites within specific groups of interest. Additional filters can be applied to visualise subsets of results, for example by selecting specific metabolites (**Fig. 6b**). The application includes three “Data Source” tabs: "EMBL" for data sets used in the AP-MALDI-MS protocol comparison (**Fig. 3**), "INTERLAB" for data sets from the interlaboratory comparison (**Fig. 5**), and "ALL" displaying all data reprocessed for comparability.

**Figure 6.**
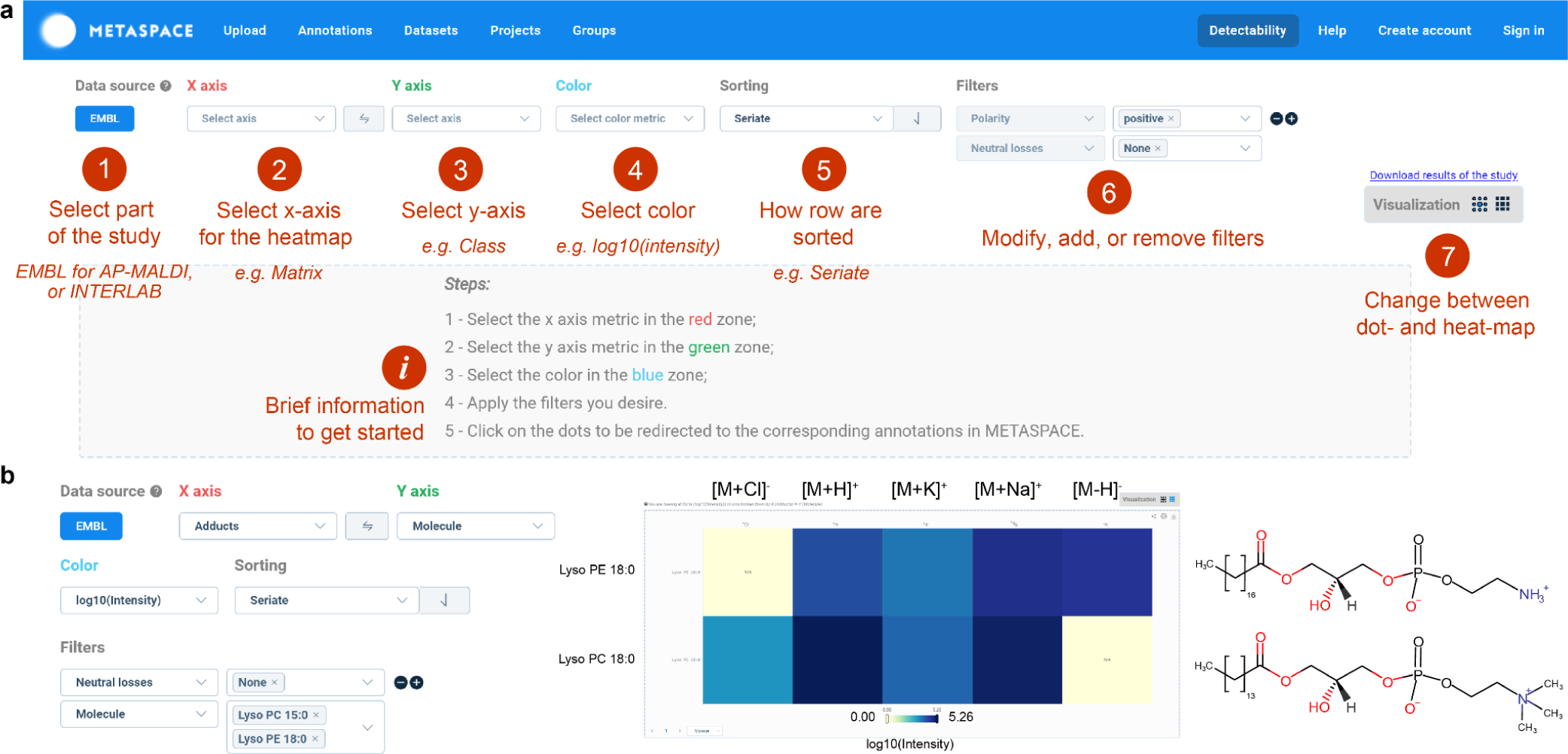
Interactive web application facilitates access to study results and data. ***a,*** *Web application interface displays customisable options and filters for visualising the results of AP-MALDI-MS protocol comparison and interlaboratory survey*. ***b****, Example: Intensities of ions of two isomeric lipids, LysoPE(18:0) and LysoPC(15:0), detected using AP-MALDI technology. Both isomers are detected in positive mode. However, they can be distinguished by examining the deprotonated ion in negative polarity, as only LysoPC(18:0) forms this ion species. Permanent URL: https://metaspace2020.eu/detectability?filter=nL,name&filterValue=None%7CLyso%20PC%2015%3A0%23Lyso%20PE%2018%3A0&xAxis=a&yAxis=name&agg=log10_intensity&metric=2&page=1&pageSize=5&src=EMBL&vis=2*

Our web resource is particularly valuable for scientists new to spatial metabolomics, aiding in the selection of appropriate polarity and MALDI matrix for untargeted spatial metabolomics or targeted detection of specific metabolites, chemical classes, or biochemical pathways. The application also identifies the most abundant ion species for a given metabolite, highlighting the potential loss of measurement sensitivity due to signal dilution. Additionally, it can point out the differences between various imaging MS technologies. **SI Table 11** provides links to example plots generated with our application. The web resource not only presents metabolite detectabilities and intensities but also connects the results to the original data sets. Users can access relevant data sets on METASPACE by clicking on dots in the dot plots or cells in the heatmaps, verifying ion image and mass spectrometry signal quality in the “Diagnostics” section of METASPACE, exploring potential isomers or isobars, and accessing experimental metadata and analysis parameters. The downloadable data in the centroided imzML format, can be further utilised for generating new hypotheses and addressing unexplored questions.

## Outlook

The main accomplishment of our study is an openly accessible and comprehensive account of metabolite detectability in a diverse range of imaging MS protocols and technologies. Using defined reference samples, we provided an unbiased empirical basis for evaluating common assumptions and situating our findings within the broader context of untargeted metabolite analysis. Beyond the shared results, our data holds potential for further analysis. Exploring unaddressed aspects of metabolite ionisation, such as in-source fragmentation and its comparison with MS/MS databases, could enhance compound identification based on MS1 data, as was demonstrated recently in a limited scope.^33^ Investigating the molecular properties of metabolites and MALDI matrices and observed detectabilities across atmospheric pressure and vacuum instruments, could offer new insights into the ion formation mechanisms in MALDI-MS and contribute to predictive theories for metabolite ionisation. Additionally, the reference samples could serve as a valuable benchmarking tool for future studies, aiding in the evaluation of new MALDI matrices and the assessment of imaging MS instrument performance following hardware or software updates. Future investigations could utilise our experimental and computational procedure to expand findings to other compounds of interest, such as plant metabolites or drugs. With further validation, the preparation of reference samples with analytes deposited on alternative substrates, like sections of homogenised tissue^34^, could facilitate the study of suppression effects observed in biological tissues. By promoting the use of standardised reference samples, we believe their availability in various formats, including commercial, would greatly enhance reproducibility and facilitate result inter-comparability in the research community. We are confident that the outcomes of this work and the opportunities they provide will significantly benefit the imaging MS community by making untargeted spatial metabolomics more effective, interpretable, and impactful.

## Methods

### Materials

Analytical standards were purchased from MetaSci, Toronto, ON, Canada; Sigma-Aldrich Chemie GmbH, Taufkirchen, Germany; Avanti Polar Lipids, Inc., Birmingham, AL, U.S.A.; GLSynthesis Inc., Worcester MA, U.S.A.(**SI Table 1**). LC-MS grade acetonitrile, methanol and water; ACS grade chloroform; sodium hydroxide (aqueous, 2M); hydrochloric acid (aqueous, 37%), nitric acid (conc.) were purchased from Fisher Scientific, Schwerte, Germany. SYLGARD 184 silicone resin and curing agent were purchased from Biesterfeld, Hamburg, Germany. Nitrogen, argon and oxygen gas of 99.999% or higher purity were from EMBL internal supply. PNDI was purchased from Ossila BV, Leiden, The Netherlands. 4-maleicanhydridoproton sponge (MAPS) was kindly provided by Prof. Dr. Karsten Niehaus, Bielefeld University, Bielefeld, Germany. Other MALDI matrix substances were purchased from Sigma-Aldrich Chemie GmbH, Taufkirchen, Germany.

### Preparation of metabolite solutions

A two-step procedure was used to prepare metabolite solutions. First, a stock solution was prepared in a solvent system suitable for a given metabolite. Next, the stock solution was diluted to the final concentration in a volatile solvent to produce a quick-drying nanospray and ensure uniform deposition. All metabolite standards were prepared to 200 µM final concentration, except for four lipid standards that were available in limited quantities. For these, concentrations of 50 or 10 µM were used as specified in the preparations table (**SI Table 2**). After preparation, all solutions were stored in HPLC vials at -80 ℃ until use.

### Production of reference samples

A custom mould with 180 regularly spaced columns was precision-cut from a solid polyoxymethylene block (**SI Fig. 1**). The 3D design is provided in **Supplementary Data 1**. Sylgard 184, a two-component PDMS silicone, was mixed according to the manufacturer’s instructions, and 4 g was poured into each mould. The moulds were placed to de-gas under vacuum for 30 min before being set to cure at 65 ℃ for 2 h. Any visible burrs or defects were manually trimmed, and freshly cured polymer templates were placed on regular glass slides that had been pre-cleaned using an acid-cleaning protocol^16^ or on indium tin oxide (ITO)-coated slides (Bruker Daltonics GmbH, Bremen, Germany). Indexing strips included in the mould design enabled the reproducible alignment of the polymer templates to standard-sized glass slides (25 x 75 mm). The templates created slides with leak-tight microwells arranged in 20 rows and 9 columns, each well measuring 1 mm in diameter. Metabolite solutions were dispensed into the microwells using an ATS4 automatic TLC sampler (CAMAG, Muttenz, Switzerland). A compatible 3D-printed sample holder for 10 glass slides was designed to fit on the stage in place of the normal silica plate. This allowed registration of well positions to the stage coordinates, requiring only a brief manual 3-point recalibration for each slide to ensure correct spray positioning across the slide surface. Calibration was verified by depositing a solution of rhodamine B and caffeine into three dedicated microwells. The dispenser was then programmed to deposit 2 µl of a single compound solution into each microwell. The programs were prepared as coordinate lists, then imported and run in “free mode” using the instrument software Wincats v. 1.4.4.6337 (**SI Fig. 2)**. The complete slide layout used can be found in the SI material (**SI Fig. 3, SI Table 3**). After compound deposition and drying, polymer templates were removed, and finished slides were vacuum-packed under an argon atmosphere, to minimise the risk of sample oxidation, and stored at -80 ℃ until use.

### AP-MALDI mass spectrometry imaging

On the day of analysis, the reference sample slides were brought to room temperature for at least 1 h before breaking the vacuum seal. Matrix substances (**SI Table 4**) were sprayed onto the reference samples using a TM Sprayer (HTX Technologies, Chapel Hill, NC, U.S.A.) following the protocols in **SI Table 5**. The reference samples were imaged using an AP-SMALDI5 AF ion source (TransMIT, Giessen, Germany) coupled to a QExactive Plus mass spectrometer (Thermo Fisher Scientific, Bremen, Germany). Instrument parameters are listed in **SI Table 6**. Each slide was imaged separately in both ion polarities, as well as in low and high mass ranges (following instrument guidelines to ensure that data is acquired in the mass range where Orbitrap mass analyser can maintain high resolution and accuracy), resulting in a total of four sequential acquisitions. To prevent repeated sampling of the same locations, the imaging raster was manually offset by 50 μm between each acquisition.

### Mass spectrometry imaging by interlaboratory survey participants

The experimental protocols used by the interlaboratory survey participants can be found in **SI Supplementary Methods**.

### Data preprocessing

AP-MALDI data were converted from the vendor raw format to mzML using MSConvert^17^ with the vendor-supplied method for spectrum centroiding. The mzML format was then converted to imzML using imzMLConverter^18^. External collaborator data was converted separately by each participant and uploaded to METASPACE^19^ as centroided imzML files. In cases where the same sample slide had been imaged separately in lower and higher mass ranges for a given polarity, the two datasets were merged pixel-by-pixel through a Python script utilising the pyimzML library to form a wide mass range spectrum per pixel. If merging was not possible, lower- and higher-mass-range data sets were processed separately, and only their result tables were merged. All images were submitted to METASPACE with the following processing parameters: positive mode adducts [M]^+^, [M+H]^+^, [M+Na]^+^, [M+K]^+^; negative mode adducts [M]^-^, [M-H]^-^, [M+Cl]^-^; neutral losses -H_2_O, -2H, -CO_2_, -CH_2_O_3_, -CH_2_O_2_, -HPO_3_, -H_3_PO_4_, -NH_3_; neutral gains +2H, +matrix (C_13_H_10_N_2_, C_10_H_7_NO_3_, C_10_H_6_ClNO_2_, C_10_H_10_N_2_, C_10_H_8_N_2_, C_8_H_8_O_3_, C_7_H_6_O_4_, C_18_H_18_N_2_O_3_, C_12_H_14_N_2_, C_11_H_8_N_2_, C_6_H_6_N_2_O_2_, C_7_H_4_ClNS_2_); mass tolerance of 3 ppm (for AP-MALDI protocol comparison) or 10 ppm (for interlaboratory technology comparison). Data annotation was performed against a custom database containing only the reference metabolites, and only one adduct and up to one either neutral loss or neutral gain per ion was considered.

### Data analysis

#### Ion image classification and determining metabolite ion detection

For each data set, we determined which spectra were sampled in each metabolite spot by fitting reference points to image features using an in-house interactive grid-fitting tool developed in Python. A significant number of ion images exhibited spatial patterns that did not correspond to the expected spots, indicating the presence of spatially distributed ions unrelated to the reference metabolites (e.g. matrix ions or other chemical noise). To differentiate between ion images representing “real signal” and those containing “background noise”, we developed a machine learning-based classifier using CatBoost^20^. We manually annotated a total of 284 ion images to train the classifier to identify ion images with the localised intensity of a putative metabolite ion in its expected well location. The model was trained using a set of pre-computed metrics for each pair of metabolite well and ion image, where at least one peak in the ion image appeared inside the well. The model features were: the average TIC-normalised intensity of the pixels inside the spot, the percentage of pixels with a non-zero intensity (further referred to as occupancy) in the spot, the ratio between the occupancy in the spot and occupancy in the "far background" (background area at least 4 spot-radii away from the spot), the ratio between the average intensities inside the spot and in the far background, and the ratio between the average intensities inside the spot and in other spots. The classifier provided a probability score, which we used to classify ions as detected or undetected. Ions with a probability score of 80% or higher were considered "real signal" and treated as "detected" for further analysis. The remaining ions were classified as "not detected." Among the undetected ions, we designated certain ions as “matrix-obscured” to indicate difficulty in determining whether the analyte was detected or not due to the high signal both within and outside the metabolite spot. Specifically, any undetected ions with intensity and occupancy ratios exceeding 0.95, as well as background occupancy rates greater than 40%, were categorised as matrix-obscured.

#### Assessment of chemical diversity of reference metabolites

To construct the chemical space, we used the chemical structures of all metabolites in the mouse genome-scale metabolic model (iMM1865^21^) from the BiGG open-source repository^22^. First, we obtained the Simplified Molecular Input Line Entry System (SMILES) notation for the considered metabolites from the Human Metabolome Database (HMDB, v.4) or by converting chemical structures to SMILES using Marvin software. The SMILES were then used to calculate "Morgan/Circular fingerprint" vectors using the RDkit Python library. The resulting vectors were visualised as 2D and 3D PCA score plots, with the metabolites used in the reference samples highlighted.

#### Visualisation

All data visualisations (dot plots, bar plots, line plots, scatter plots, and heatmaps) were performed in Python. A map of common biochemical pathways with highlighted reference metabolites was created using IPath v.3^23^ and KEGG identifiers obtained from HMDB (lipids without KEGG identifiers were excluded from the visualisation).

To compare metabolite detectabilities between AP-MALDI protocols and other imaging MS technologies, only detected ions without neutral losses or neutral gains were considered. If at least one ion was detected for a metabolite, it was considered detected. For each dataset, the fraction of metabolites detected per chemical subclass or biochemical pathway was calculated, along with the mean intensity of detected ions.

To compare metabolite detectability between reference samples and biological tissues, we chose 38 tissue data sets acquired on the same instrument as described for AP-MALDI mass spectrometry imaging using either DHB matrix in positive polarity or DAN matrix in negative polarity, and available on METASPACE. Data annotation was performed in the same manner as for reference sample data, except we considered only main ion types: [M+H]^+^, [M+Na]^+^, and [M+K]^+^ in positive polarity and [M-H]^-^ and [M+Cl]^-^ in negative polarity. We obtained 15550 ion images of reference metabolites, and manually annotated each ion as detected or not detected. Every metabolite for which at least one ion was detected, was considered detected. For each data set, we calculated the fraction of metabolites detected per chemical subclass. Then, we averaged the detectability values per chemical subclass across tissue data sets. In the final comparison, only ions with the same adducts were considered in the reference samples.

To calculate the average signal composition in a dataset, all considered ions, including those with neutral losses and neutral gains, were taken into account. First, we calculated the relative abundances of detected ions for each metabolite. Next, the average abundances of detected ions per chemical class in each data set were determined. The obtained values were further averaged across imaging MS technologies for the summary plots.

#### Predicting metabolite detectability and intensities

Machine learning was used to predict metabolite intensities for AP-MALDI-MS. First, we trained a CatBoost classifier to predict the detectability of a given molecule for a combination of the MALDI matrix and polarity. As described in the Methods, we defined a metabolite to be detected if at least one of its ions achieved a probability score of 80% or higher. Matrix-obscured ions were excluded from the analysis. Seven predicted molecular properties from HMDB were used as features for each metabolite: "pka_strongest_acidic", "pka_strongest_basic", "polar_surface_area", "polarizability", "acceptor_count", "donor_count", "physiological_charge". These feature values were normalised using a power transform to achieve Gaussian distribution. Next, we trained a regression model to predict the sum intensities of only detected metabolites using a neural network with one layer (100 features), ReLu activation function, and Adam optimizer. The model was trained using scikit-learn (v1.1.1, class sklearn.neural_network.MLP) with default parameters, employing the Binary Cross-Entropy loss function for classification and the Mean Squared Error loss function for regression. The model’s performance was validated using 10-fold cross-validation, ensuring diverse folds by stratifying the data points based on all features. We compared our model’s performance to a baseline predictor using the same data with randomly shuffled features across all metabolites and reported the average scores plus standard deviations over 10 repetitions.

After training the models, we computed and plotted the SHAP (SHapley Additive exPlanations) values to determine how much each molecular property contributed to the model output.

### Web application

We developed a web application to provide interactive access to the results generated in this study. The application was built as a Single Page Application using the Vue.JS framework and the eCharts chart library. It allows users to create visualisations in the form of heatmaps or dot plots by specifying various parameters. The web application offers the following options for the X and Y axes: adduct, chemical class, chemical subclass, dataset ID, MALDI matrix, metabolic pathway, molecule, neutral losses, polarity and sample name. Users can choose to visualise properties such as raw or mean intensity, log10-transformed intensity, and the fraction of detected metabolites per category as the size and/or colour. The results can be further filtered by adding one or multiple filters (e.g. to view a specific metabolite in a polarity of interest). The application provides three data tabs: EMBL, INTERLAB, and ALL. The EMBL tab contains data sets used for AP-MALDI protocol comparison (see **Fig. 3**), processed with 3 ppm mass tolerance, and provides information on both detectabilities and intensities of the metabolite ions. The INTERLAB contains data sets used for interlaboratory comparison (see **Fig. 5**), processed with 10 ppm mass tolerance. The ALL tab includes all datasets processed with a 10 ppm mass tolerance. Both the tab INTERLAB and ALL provide only ion detectability information as the intensities are not directly comparable across different technologies.

### Data availability

The imaging MS data collected in this study are publicly available on METASPACE (https://metaspace2020.eu/project/saharuka-2024) for browsing and download. The individual data set metadata and links are provided (**SI Table 8, Supplementary Data 2**). The results of the study are also provided in a comma-separated values (CSV) file format as SI material (**Supplementary data 3**). The source code for the data analysis can be accessed on GitHub (https://github.com/saharuka/metabolite_detectability and https://github.com/alexandrovteam/predicting-APMALDI-response). The web application developed for this study is open-source and available as a part of the METASPACE platform at https://metaspace2020.eu/detectability, with the codebase available on GitHub (https://github.com/metaspace2020/metaspace).

## Supporting information

Supplement Guide

## Acknowledgements

We thank Prof. Dr. Karsten Niehaus for providing the MAPS matrix, EMBL mechanical workshop for help with design and production of the mould, Dr. Prasad Phapale from the EMBL Metabolomics Core Facility for technical assistance, and Sebastian Bessler (Münster University) for assistance with MALDI measurements.

## CRediT author statement

**Veronika Saharuka**: conceptualization (lead); methodology (lead); investigation (equal); formal analysis (lead); visualization; project administration; writing – original draft (lead). **Lucas Maciel Vieira**: software; formal analysis (supporting); visualization. **Lachlan Stuart**: software; formal analysis (supporting); visualization. **Måns Ekelöf**: methodology (supporting); investigation (equal); formal analysis (supporting); writing – original draft (supporting). **Martijn R. Molenaar**: formal analysis (supporting); writing – review & editing. **Alberto Bailoni**: formal analysis (supporting); visualization. **Katja Ovchinnikova**: formal analysis (supporting). **Theodore Alexandrov**: supervision; conceptualization (supporting); writing – original draft (supporting); funding acquisition. All remaining authors (**Jens Soltwisch, Tobias Bausbacher, Dusan Velickovic, Dennis Jakob, Fernanda E. Pinto, Nicole Strittmatter, Mary King, Crystal Pace, Max A. Müller, Janina Oetjen, Bernhard Spengler, David C. Muddiman, Livia S. Eberlin, Richard Goodwin, Christian Janfelt, Manuel Liebeke, Christopher R. Anderton, Carsten Hopf, Klaus Dreisewerd**) performed the analysis of reference samples, contributed their laboratory protocols and the data to the interlaboratory survey, and edited the manuscript.

