## Supplement Guide for "Large-Scale Evaluation of Spatial Metabolomics Protocols and Technologies"

**Supplementary Information** contains supplementary methods provided by all interlaboratory study participants, 11 supplementary tables and 16 supplementary figures.
**Supplementary Data 1**.STEP - 3D design used for the production of PDMS mould
**Supplementary Data 2**.csv - Links and metadata for data sets uploaded to METASPACE
**Supplementary Data 3**.csv - Results for MALDI matrix comparison
**Supplementary Data 4**.csv - Chemical standards information
