## Supplementary Information for "Large-Scale Evaluation of Spatial Metabolomics Protocols and Technologies"

[1] Structural and Computational Biology Unit, European Molecular Biology Laboratory, Heidelberg, Germany, [2] Collaboration for joint PhD degree between EMBL and Heidelberg University, Faculty of Biosciences, Heidelberg, Germany; [3] Institute of Hygiene, University of Münster, Münster, Germany, [4] Center for Mass Spectrometry and Optical Spectroscopy (CeMOS), Mannheim University of Applied Sciences, Mannheim, Germany, [5] Mannheim Center for Translational Neuroscience (MCTN), Medical Faculty Mannheim, Heidelberg University, Mannheim, Germany, [6] Max Planck Institute for Marine Microbiology, Bremen, Germany,
[7] Baylor College of Medicine, Houston, TX, U.S.A., [8] Justus Liebig University Giessen, Institute of Inorganic and Analytical Chemistry, Giessen, Germany, [9] Bruker Daltonics GmbH & Co. KG, Bremen, Germany, [10] Department of Chemistry, North Carolina State University, NC, U.S.A., [11] Department of Pharmacy, University of Copenhagen, Copenhagen, Denmark, [12] Imaging and Data analytics, Clinical Pharmacology and Safety Sciences, R&D, AstraZeneca, Cambridge, UK, [13] Environmental Molecular Sciences Laboratory, Pacific Northwest National Laboratory, WA, U.S.A., [14] Institute for Metabolomics, Kiel University, Germany, [15] Medical Faculty, Heidelberg University, Heidelberg, Germany, [16] Metabolomics Core Facility, EMBL, Heidelberg, Germany, [17] Molecular Medicine Partnership Unit, EMBL and Heidelberg University, Heidelberg, Germany, [18] BioStudio, BioInnovation Institute, Copenhagen, Denmark

### Supplementary Methods

#### Methods for 10I/10G

MALDI imaging analyses were performed on a Bruker Daltonics 15T solariX FTICR MS, equipped with a ParaCell. This instrument has an Apollo II ESI and MALDI source with a SmartBeam II frequency-tripled (355 nm) Nd: YAG laser (Bremen, Germany).

##### Positive ion mode analyses

DHB was prepared at a concentration of 40 mg/mL DHB (70% MeOH) and was sprayed using HTX TM-5 sprayer (HTX Technologies, Chapel Hill, NC) at a flow rate of 50 µL/min. The nozzle temperature was set to 70 °C, with 12 cycles at 3 mm track spacing with a crisscross pattern. A 2 s drying period was added between cycles, linear flow was set to 1,200 mm/min with 10 PSI of nitrogen gas and 40 mm nozzle height. This resulted in matrix coverage of ~667 µg/cm$^2$ for DHB.

FTICR was optimised separately for the analysis of low and high *m/z* ions. For low *m/z* ions, data were acquired with broadband excitation from *m/z* 92.1 to 400, resulting in a detected transient of 0.214 s — the observed mass resolution was ~100k at *m/z* 400. Analyses were performed using a 100 µm step size, “small” laser focus at 2,000 Hz with 200 laser shots. For high *m/z* ions data were acquired with broadband excitation from *m/z* 153.6 to 1500.0, resulting in a detected transient of 0.357 s — the observed mass resolution was ~170k at *m/z* 400. Analyses were performed using a 100 µm step size, “small” laser focus at 2,000 Hz with 250 laser shots.

##### Negative ion mode analyses

NEDC was prepared at a concentration of 7 mg/mL in 70% MeOH and was sprayed at a flow rate of 120 µL/min using the M5-Sprayer. The nozzle temperature was set to 70 °C, with 8 cycles at 3 mm track spacing with a crisscross pattern. No drying period was added between cycles, and linear flow was set to 1200 mm/min with 10 PSI of nitrogen gas and 40 mm nozzle height. This resulted in matrix coverage of ~187 µg/cm$^2$ for NEDC.

FTICR was optimised separately for the analysis of low and high *m/z* ions. For low *m/z* ions data were acquired with broadband excitation from *m/z* 92.1 to 500, resulting in a detected transient of 0.214 s — the observed mass resolution was ~100k at *m/z* 400. Analyses were performed using a 100 µm step size, “small” laser focus at 2,000 Hz with 200 laser shots. For high *m/z* ions data were acquired with broadband excitation from *m/z* 207.3 to 1500.0, resulting in a detected transient of 0.482 s — the observed mass resolution was ~230k at *m/z* 400. Analyses were performed using a 100 µm step size, “small” laser focus at 2,000 Hz with 250 laser shots.

#### Methods for 8A/8D

##### Sample preparation

| Matrix | 9AA | DHB |
|---|---|---|
| Matrix concentration | 5 mg/mL | 30 mg/mL |
| Solvent composition | 70/30 Ethanol/water | 50/50 Acetone/water; |

| | | |
|---|---|---|
| | | 0.1 vol% TFA added to the solvent mix |
| Method | SMALDIPrep area spraying | SMALDIPrep area spraying |
| Spraying rate | 20 µl/min | 20 µl/min |
| Application speed | 2 mm/s | 2 mm/s |
| Application mode | xy-raster | xy-raster |

##### Instrumentation

| **Parameter** | **Low mass** | **High mass** |
|---|---|---|
| Attenuator angle* | 20° | 20° |
| Pixel size | 100 µm | 100 µm |
| **Mass range** | ***m/z* 75 - 300** | ***m/z* 300 - 1200** |
| Voltage | ±3 kV | ±3 kV |
| S-Lens RF level | 100 | 100 |
| Injection time | 500 ms | 500 ms |
| Mass resolving power | 240,000 | 240,000 |
| Lock mass positive-ion mode | 273.0394 | 716.1246 |
| Lock mass negative-ion mode | None | None |

*Individual AP-SMALDI5 instruments require different attenuator settings to reach the same results, as the precise laser energy output onto the sample depends not only on the attenuator but also on the exact beam path through the optics, which can vary slightly between instruments.

##### Chemical and instrument vendor information

| **Instrumentation / Chemical** | **Vendor** |
|---|---|
| Ion source | AP-SMALDI5 AF, TransMIT GmbH, Giessen, Germany |
| Mass spectrometer | Q Exactive HF, Thermo Fisher Scientific, Bremen, Germany |
| Sprayer | SMALDIPrep, TransMIT GmbH, Giessen, Germany |
| Water | VWR Chemicals, Darmstadt, Germany |
| Acetone | Merck, Darmstadt, Germany |
| Ethanol | Merck, Darmstadt, Germany |
| TFA | Merck, Darmstadt, Germany |
| DHB | Merck, Darmstadt, Germany |
| 9AA | TCI Deutschland GmbH, Eschborn, Germany |

#### Methods for 9A/9E

Provided glass slides were placed on a translational stage and cooled to -8℃ inside a humidity-controlled enclosure. Once frozen, a thin ice layer was formed by controlled exposure to ambient humidity. A mid-IR laser was utilised for desorption of spotted standards at a wavelength of 2.94 µm and an energy of 0.9 mJ per shot resulting in a spot size of approximately 100 µm. Undersampling allowed two analytical runs per glass slide offset on the y-axis (200 µm). Fresh electrospray solvent was prepared daily for the post-ionisation of spotted standards. For positive ionisation, electrospray consisted of 60% acetonitrile in water containing 0.2% formic acid and was stabilised between 3200-3500 V with a flow rate of 2 µL/min. For the negative ionisation mode, electrospray solvent consisted of 60% acetonitrile in water containing 1 mM acetic acid and was stabilised between 3000 and 3200 V at a flow rate of 1.5 µL/min. The IR-MALDESI source was coupled to an Exploris 240 operating with automatic gain control (AGC) turned off at a fixed injection time of 15 ms for optimal transmission with a resolving power of 240,000. Two mass ranges, 75-350 *m/z* (RF ratio set to 5) and 200-1550 *m/z* (RF ratio set to 8) were analysed per glass slide in both ionisation modes. The internal calibrant was utilised for lock mass correlation to achieve high mass accuracy.

#### Methods for 7J/7A

Please see the Methods section of the main text.

#### Methods for 5HJ/5GI

Analysis was performed on a Thermo QExactive orbitrap mass spectrometer equipped with a custom-built DESI-MSI ion source, as described in detail elsewhere.[1] A mixture of methanol and water (90:10) was used as spray solvent, supplied at a flow rate of 5 µL/min, using a nebuliser gas pressure of 6 bar. Imaging was performed at 150 µm pixel size. To ensure optimal detection of the entire *m/z* range of the analytes, the detection was split into two mass ranges of *m/z* 75-500 and *m/z* 100-1500 respectively, for both positive and negative ion modes.

#### Methods for 12C/12E

##### Chemicals

2,5-Dihydroxybenzoic acid (DHB; 99% CAT#: A11459) was purchased from Alfa Aesar. 1,5-Diaminonaphthalene (DAN; 97% CAT#: D21200) and Trifluoroacetic acid (CAT#: 108262) were purchased from Sigma-Aldrich. LC-MS grade Acetonitrile (99.9% CAT#: 34967) was purchased from Honeywell Research Chemicals.

##### Sample preparation

Sample slides were stored at -80 °C until use. Matrix solutions were sonicated for 10 minutes before spraying. Immediately before use, the vacuum-sealed slide mailer was left to equilibrate at room temperature for 30 minutes. After breaking the vacuum seal, a bright field image was recorded using a Leica Aperio CS2 slide scanner, and the matrix was applied using an HTX M5 matrix sprayer with the following spray parameters.

| | **Spray Method 1 (positive ion mode)** | **Spray Method 2 (negative ion mode)** |
|---|---|---|
| **Matrix** | 2,5-Dihydroxybenzoic acid (DHB) | 1,5-Diaminonaphtalene (DAN) |
| **Concentration [mg/mL]** | 50 | 10 |
| **Solvent composition** | 70% ACN 29.9% H2O 0.1% TFA | 70% ACN 30 % H2O |
| **Nozzle temperature [°C]** | 75 | 75 |
| **Number of passes** | 8 | 17 |
| **Flow rate [mL/min]** | 0.1 | 0.05 |
| **Velocity [mm/min]** | 1200 | 1200 |
| **Track spacing [mm]** | 3 | 2 |
| **Pressure [psi]** | 10 | 10 |
| **Nozzle height [mm]** | 40 | 40 |

#### Data acquisition

##### FT-ICR (Bruker solariX XR 7T)

Data acquisition was performed on a 7T Fourier-transform ion cyclotron resonance (FT-ICR) mass spectrometer (MS) (solariX XR 7T, Bruker Daltonik) in two steps for each polarity. First, a method optimised for the detection of molecules in the *m/z* range 65-700 was used at a lateral step size of 100 µm. After that, a method optimised for the detection of molecules in the *m/z* range of 500-1700 was used on the same slide with an XY offset of 50 µm and a lateral step size of 100 µm. Peak filtering was set to SNR >3 and an absolute intensity threshold of $10^5$ a.u.

| FT-ICR methods | Method 1 | Method 2 | Method 3 | Method 4 |
|---|---|---|---|---|
| **Polarity** | Positive | Negative | Positive | Negative |
| **Low *m/z*** | 64.5 | 64.5 | 129 | 118 |
| **High *m/z*** | 700 | 700 | 1700 | 1700 |
| **Data size** | 2M | 2M | 1M | 1M |
| **Laser shots** | 300 | 150 | 300 | 150 |
| **Laser Frequency [1/s]** | 2000 | 1000 | 2000 | 1000 |
| **Resolving power at *m/z* 400** | 85,000 | 85,000 | 85,000 | 77,000 |
| **Funnel RF Amplitude [Vpp]** | 150 | 120 | 250 | 250 |
| **Q1 mass** | 150 | 80 | 600 | 600 |
| **Collision RF Amplitude [Vpp]** | 1600 | 1600 | 1600 | 1600 |
| **Time of Flight [ms]** | 0.4 | 0.8 | 0.9 | 1.2 |
| **Data Reduction factor** | 97% | 97% | 97% | 97% |

Centroided data was imported into SCiLS Lab 2023a Pro, exported as imzML file and uploaded to METASPACE using mass tolerance of 3 ppm. The resolving power was set to the estimated resolving power at *m/z* 400 provided by ftmsControl.

### Methods for 12G/12I

#### Chemicals

2,5-Dihydroxybenzoic acid (DHB; 99% CAT#: A11459) was purchased from Alfa Aesar. 1,5-Diaminonaphthalene (DAN; 97% CAT#: D21200) and Trifluoroacetic acid (CAT#: 108262) were purchased from Sigma-Aldrich. LC-MS grade Acetonitrile (99.9% CAT#: 34967) was purchased from Honeywell Research Chemicals.

#### Sample preparation

Sample slides were stored at -80 °C until use. Matrix solutions were sonicated for 10 minutes before spraying. Immediately before use, the vacuum-sealed slide mailer was left to equilibrate at room temperature for 30 minutes. After breaking the vacuum seal, a bright field image was recorded using a Leica Aperio CS2 slide scanner, and the matrix was applied using an HTX M5 matrix sprayer with the following spray parameters.

| | Spray Method 1 (positive ion mode) | Spray Method 2 (negative ion mode) |
|---|---|---|
| **Matrix** | 2,5-Dihydroxybenzoic acid (DHB) | 1,5-Diaminonaphtalene (DAN) |
| **Concentration [mg/mL]** | 50 | 10 |
| **Solvent composition** | 70% ACN 29.9% H2O 0.1% TFA | 70% ACN 30 % H2O |
| **Nozzle temperature [°C]** | 75 | 75 |
| **Number of passes** | 8 | 17 |
| **Flow rate [mL/min]** | 0.1 | 0.05 |
| **Velocity [mm/min]** | 1200 | 1200 |
| **Track spacing [mm]** | 3 | 2 |
| **Pressure [psi]** | 10 | 10 |
| **Nozzle height [mm]** | 40 | 40 |

#### Data acquisition

##### qTOF (Bruker TimsTOF flex)

Data acquisition was performed on a MALDI-TIMS-QTOF mass spectrometer (TimsTOF flex, Bruker Daltonik) with the ion mobility option deactivated. For each polarity, data acquisition was performed in two steps. First, a method optimised for the detection of molecules in the *m/z* range of 60-700 was used at a lateral step size of 100 µm. After that, a method optimised for the detection of molecules in the *m/z* range of 400-1700 was used on the same slide with an XY offset of 50 µm and a lateral step size of 100 µm.

| timsTOF flex Methods | Method 1 | Method 2 | Method 3 | Method 4 |
|---|---|---|---|---|
| **Polarity** | Positive | Negative | Positive | Negative |
| **Scan Begin *m/z*** | 60 | 60 | 400 | 400 |
| **Scan End *m/z*** | 700 | 700 | 1700 | 1700 |
| **Laser Geometry** | M5 small | M5 small | M5 small | M5 small |
| **Laser shots** | 200 | 200 | 200 | 200 |
| **Laser Frequency [1/s]** | 10,000 | 1000 | 1000 | 1000 |
| **Resolving power at *m/z* 400** | 35,000 | 35,000 | 35,000 | 35,000 |
| **Funnel 1 RF [Vpp]** | 200 | 200 | 300 | 300 |
| **Funnel 1 RF [Vpp]** | 250 | 250 | 300 | 300 |
| **Multipole RF [Vpp]** | 200 | 200 | 300 | 300 |
| **Collision Energy [eV]** | 15 | 3 | 10 | 10 |
| **Collision RF [Vpp]** | 1000 | 200 | 2500 | 2500 |
| **Low Mass *m/z*** | 200 | 100 | 500 | 300 |
| **Transfer Time [µs]** | 50 | 50 | 90 | 100 |
| **Pre Pulse Storage [µs]** | 12 | 5 | 10 | 7 |

Centroided data was imported into SCiLS Lab 2023a Pro, exported as imzML file and uploaded to METASPACE using mass tolerance of 3 ppm. The resolving power was set to 30,000.

#### Methods for 1JI/1GH

All DESI-MSI analyses were performed on a Thermo Q-Exactive HF Orbitrap mass spectrometer fitted with a laboratory-built DESI-MS source at a resolving power of 60,000 and spatial resolution of 100 µm. The nebulizing $N_2$ gas pressure was set to 180 psi. For negative ion mode analysis, 1:3 DMF/ACN (v/v) was used as the spray solvent at a flow rate of 1.5 µL/min. For positive ion mode analysis, 100% ACN was used as the spray solvent at a flow rate of 3 µL/min. MS imaging data were collected in two mass ranges for both ion modes: the “low mass” range at *m/z* 75-400 and the “high mass” range at *m/z* 400-1500.

#### Methods for 14B/14D

All MALDI imaging analyses were performed on a Q Exactive HF (Thermo Scientific, Bremen, Germany), equipped with an AP-SMALDI10 source (TransMIT, Gießen, Germany). This source has a nitrogen laser with a pulse rate of 60 Hz. The mass resolution was set to 240,000 @200 *m/z* and the automatic gain control was set to 1,000,000. Ions from 30 laser pulses were accumulated for each mass spectrum. The attenuator was set to 20º. The mass spectrometer was calibrated with a mass accuracy (rms) of <1 ppm using red phosphorus. Data was acquired using a pixel size of 100 µm.

| **Parameter** | **Low mass** | **High mass** |
|---|---|---|
| Attenuator angle* | 20° | 20° |
| Pixel size | 100 µm | 100 µm |
| **Mass range** | ***m/z* 75 - 500** | ***m/z* 300 - 1500** |
| Voltage | ±3 kV | ±3 kV |
| S-Lens RF level | 100 | 100 |
| Injection time | 500 ms | 500 ms |
| Mass resolving power | 240,000 | 240,000 |

Slides were stored at -80ºC prior processing. All slides were covered with matrix using a SunCollect Matrix Sprayer (SunChrom, Friedrichsdorf, Germany), applying a constant nitrogen flow of 2 bar.

##### sDHB spraying for positive mode analyses

Super-DHB matrix (sDHB, 9:1 (w/w) mixture of 2,5-dihydroxybenzoic acid and 2-hydroxy-5-methoxybenzoic acid, Merck, Darmstadt, Germany) was prepared in a concentration of 30 mg/mL in methanol:water:TFA (50:50:0.1). Spraying was performed in a meandering movement with 2 mm line distance and a nozzle distance of 25 mm. The X- and Y-axis velocity were ‘low 6’ and ‘medium 3’ respectively. A total of 10 layers were applied to the slide where the first layer was deposited with 10 µL/min flow rate and layers 2-10 with 15 µL/min flow rate.

##### 9AA spraying for negative ion mode analyses

9-aminoacridine (9AA, Honeywell Fluka, Morristown, USA) was prepared in a concentration of 10 mg/mL in methanol:water:TFA (70:30:0.1). Spraying was performed in a meandering movement with 2 mm line distance and a nozzle distance of 25 mm. The X- and Y-axis velocities were both set to ‘medium 1’. A total of 8 layers were applied to the slide where the first layer was deposited with 10 µL/min, the second with 20 µL/min, the third with 30 µL/min and layers 4-8 with 40 µL/min flow rate.

### Methods for 13G/13J

#### Chemicals

2,5-Dihydroxybenzoic acid (DHB; CAT#: 8201346) was purchased from Bruker. N-(1-Naphthyl)ethylendiamin-dihydrochlorid (NEDC; CAT#: 222488) and Trifluoroacetic acid (CAT#: 108262) were purchased from Sigma-Aldrich. ULC-MS grade Acetonitrile (99.9% CAT#: 012041) and Methanol (99.98%; CAT# 136841) were purchased from Biosolve Chimie SARL.

#### Sample preparation

Sample slides were stored at -80 °C until use. Matrix solutions were sonicated for 10 minutes before spraying. Immediately before use, the vacuum-sealed slide mailer was left to equilibrate at room temperature for 30 minutes. Before matrix spraying, a low-resolution grayscale scan was generated on an Epson Perfection V850 Pro scanner with 3200 dpi image resolution. The matrix was applied using a HTX TM sprayer with the following spray parameter:

| | **Spray Method 1 (positive ion mode, 13G)** | **Spray Method 2 (negative ion mode, 13J)** |
|---|---|---|
| **Matrix** | 2,5-Dihydroxybenzoic acid (DHB) | N-(1-Naphthyl)ethylendiamin-dihydrochlorid (NEDC) |
| **Concentration [mg/mL]** | 15 | 7 |
| **Solvent composition** | 90% ACN, 9.9% $H_2O$, 0.1% TFA | 70% methanol, 30 % $H_2O$ |
| **Nozzle temperature [°C]** | 60 | 75 |
| **Number of passes** | 14 | 10 |
| **Flow rate [mL/min]** | 0.125 | 0.12 |
| **Velocity [mm/min]** | 1200 | 1200 |
| **Track spacing [mm]** | 3 | 3 |
| **Pressure [psi]** | 10 | 10 |
| **Nozzle height [mm]** | 40 | 40 |

#### Data acquisition

##### qTOF (Bruker timsTOF fleX)

Data acquisition was performed on a MALDI-TIMS-QTOF mass spectrometer (timsTOF fleX, Bruker Daltonics GmbH & Co. KG) with the ion mobility option deactivated. For each polarity, data acquisition was performed in two steps. First, a method optimised for the detection of molecules in the *m/z* range of 60-700 was used at a lateral step size of 100 µm. After that, a method optimised for the detection of molecules in the *m/z* range of 400-1700 was used on the same slide with an XY offset of 50 µm and a lateral step size of 100 µm.

| **timsTOF flex Methods** | **Method 1 (13J)** | **Method 2 (13J)** | **Method 3 (13G)** | **Method 4 (13G)** |
|---|---|---|---|---|
| **Polarity** | Negative | Negative | Positive | Positive |
| **Scan Begin *m/z*** | 50 | 250 | 50 | 300 |
| **Scan End *m/z*** | 900 | 1500 | 600 | 1500 |

| | | | | |
|---|---|---|---|---|
| **Laser Geometry** | Singe with 45x45µm beam scan pattern | Singe with 45x45µm beam scan pattern | Singe with 45x45µm beam scan pattern | Singe with 45x45µm beam scan pattern |
| **Laser shots** | 400 | 600 | 400 | 400 |
| **Laser Frequency [1/s]** | 10,000 | 10,000 | 10,000 | 10,000 |
| **Resolving power at *m/z* 400** | 35,000 | 35,000 | 35,000 | 35,000 |
| **Funnel 1 RF [Vpp]** | 250 | 350 | 300 | 350 |
| **Funnel 2 RF [Vpp]** | 250 | 350 | 300 | 350 |
| **Multipole RF [Vpp]** | 300 | 350 | 300 | 350 |
| **Collision Energy [eV]** | 10 | 10 | 10 | 10 |
| **Collision RF [Vpp]** | 800 | 2800 | 580 | 3500 |
| **Low Mass *m/z*** | 50 | 250 | 50 | 300 |
| **Transfer Time [µs]** | 50 | 110 | 65 | 75 |
| **Pre Pulse Storage [µs]** | 5 | 10 | 5 | 10 |

Centroided data was imported into SCiLS Lab 2023a MVS, exported as imzML file and uploaded to METASPACE using mass tolerance of 3 ppm. The resolving power was set to 30,000.

#### Methods for 1A/1C

Data was recorded using a home-built DESI spray assembly mounted to an automated 2D imaging source (Prosolia Inc, IN, USA) and a Thermo Scientific Q-Exactive Plus mass spectrometer. Four slides were measured, slide A and D in positive ion mode, Slide B and C in negative ion mode. Each slide was measured optimised for high masses and low masses, respectively. Common settings were DESI geometrical settings (sprayer to surface distance of 2 mm, sprayer to MS inlet distance of 7 mm, inlet so surface distance of <0.1 mm), capillary temperature of 320 °C, injection tome of 150 ms, AGC target of 5E6 (AGC turned off). Spray parameters were ±4,5 kV spray voltage and 1.5 µL/min methanol/water (95:5 v/v as spray solvent) and a nebulising gas pressure of 6.5 bar Nitrogen N4.8. Imaging lateral resolution was 100 µm (pixels size in the x-direction). Low mass data was recorded from *m*/*z* 70 to 400, using an S-Lens RF Level of 50. High mass data was recorded from *m*/*z* 350-1300 using an S-Lens RF setting of 100. To achieve high and low mass measurements from the same slides, settings per row were alternated while the image was recorded at a row height (pixel size in y-direction) of 50 µm.

#### Methods for 2I/7B

##### Sample preparation

2,5-dihydroxyacetophenone (DHAP) was used as the MALDI matrix. Spray-coating was performed using an ultrasonic sprayer (SimCoat with ACCUMIST nozzle, Sono-Tek, Milton, NY). The matrix was dissolved to 15 mg/mL in ACN:MeOH:H2O (8:1:1, v/v/v). The syringe pump of the sprayer was operated at a flow rate of 50 µL/min, with the ultrasonic nebulizer working at 50% in “power mode” and the nitrogen backing gas at ~70 hPa. The distance of the spray nozzle outlet from the sample surface was set to 60 mm. Robotic spraying was performed by continuously moving the nozzle across the sample with a scan speed of 30 mm/s and a meandering pattern with a track spacing of 1.8 mm. Each cycle included two passes across the whole sample area with a change of pattern tracks by 90° at half-time. Twenty cycles were sprayed with a resting period of 10 s after each cycle to allow for solvent evaporation.

##### Sample analysis

A modified Synapt G2 Si (Waters) was used for MALDI-MS imaging analysis in continuous raster mode. Details of the setup are described elsewhere[2]. In short, the ion source was modified to allow the use of increased pressure at 0.7 mbar. A $N_2$-Laser was used for desorption at a repetition rate of 30 Hz, a laser spot size of ca. 25 µm in diameter, and a laser pulse energy of ca. 9.6 µJ. Waters Research Enabled Software (WREnS) enabled a typewriter-like continuous raster mode. A scan speed of 0.2 mm/s and a scan time of 0.5 s was used. Together with a distance between lines of 100 µm this produced a pixel size of 100 X 100 µm². Data were acquired in the “high-resolution mode” of the instrument. Image processing was performed after lock mass correction and conversion of the data into centroid format using MassLynx. This was followed by the conversion of the .raw data into .imzML format.

#### Methods for 4F/3A

##### Sample preparation

2,5-dihydroxyacetophenone (DHAP) was used as the MALDI matrix. Spray-coating was performed using an ultrasonic sprayer (SimCoat with ACCUMIST nozzle, Sono-Tek, Milton, NY). The matrix was dissolved to 15 mg/mL in ACN:MeOH:H2O (8:1:1, v/v/v). The syringe pump of the sprayer was operated at a flow rate of 50 µL/min, with the ultrasonic nebuliser working at 50% in “power mode” and the nitrogen backing gas at ~70 hPa. The distance of the spray nozzle outlet from the sample surface was set to 60 mm. Robotic spraying was performed by continuously moving the nozzle across the sample with a scan speed of 30 mm/s and a meandering pattern with a track spacing of 1.8 mm. Each cycle included two passes across the whole sample area with a change of pattern tracks by 90° at half-time. Twenty cycles were sprayed with a resting period of 10 s after each cycle to allow for solvent evaporation.

##### Sample analysis

A QExactive plus (Thermo Fisher) equipped with a dual ion funnel source (spectroglyph) adapted for MALDI-2 was used for the analysis as described elsewhere[3]. The orbitrap was set to a resolving power of 140000 at 200 m/z with a fixed acquisition time of 500 ms. The mass range was set to *m/z* 75 to *m/z* 1125. Pixel size was set to 100 X 100 µm² and laser spot size was approximately 50 µm in diameter. Laser power was adjusted to produce optimal MALDI-2 spectra. Negative and positive ion mode samples were measured on the same sample with an offset of 50 µm between measurement runs. Pressure in the ion source was set to 10 mbar and a delay of 10 µs between ablation and postionization laser pulses was used for MALDI-2. Data was converted into imzML using image insight (spectroglyph).

### Supplementary Tables

#### Supplementary Table 1

Chemical standards with vendor information. Vendors represented:

1. MetaSci, Toronto, ON, Canada
2. Sigma-Aldrich Chemie, Taufkirchen, Germany
3. Avanti Polar Lipids, Birmingham, AL, U.S.A.
4. GLSynthesis, Worcester, MA, U.S.A.

| Name | Formula | Mass | Vendor |
|---|---|---|---|
| 2-Oxoglutaric acid | C5H6O5 | 146.0215 | 1 |
| 4,5-Dihydroorotic acid | C5H6N2O4 | 158.0328 | 1 |
| 5-Hydroxyindole-3-acetic acid | C10H9NO3 | 191.0582 | 1 |
| Acetylcholine chloride | C7H16NO2 | 146.1181 | 1 |
| Adenine hydrochloride | C5H5N5 | 135.0545 | 1 |
| Adenosine 5'-diphosphate potassium salt | C10H15N5O10P2 | 427.0294 | 1 |
| Adenosine 5'-monophosphate | C10H14N5O7P | 347.0631 | 1 |
| Adenosine 5'-triphosphate disodium salt | C10H16N5O13P3 | 506.9958 | 1 |
| Agmatine sulfate | C5H14N4 | 130.1218 | 1 |
| all-trans-Retinoic acid | C20H28O2 | 300.2089 | 1 |
| alpha-Tocopherol | C29H50O2 | 430.3811 | 1 |
| Ascorbic acid | C6H8O6 | 176.0321 | 1 |
| Biotin | C10H16N2O3S | 244.0882 | 1 |
| Butyric acid sodium salt | C4H8O2 | 88.0524 | 1 |
| Carbamoyl phosphate dilithium salt hydrate | CH4NO5P | 140.9827 | 1 |
| Cholecalciferol | C27H44O | 384.3392 | 1 |
| Cholesterol | C27H46O | 386.3549 | 1 |
| Cholesteryl heptadecanoate | C44H78O2 | 638.6002 | 1 |
| Cholic acid | C24H40O5 | 408.2876 | 1 |
| Choline chloride | C5H14NO | 104.1075 | 1 |
| cis-Aconitic acid | C6H6O6 | 174.0164 | 1 |
| Citric acid | C6H8O7 | 192.0270 | 1 |
| Creatine anhydrous | C4H9N3O2 | 131.0695 | 1 |
| Creatine phosphate disodium salt hydrate | C4H10N3O5P | 211.0358 | 1 |
| Creatinine | C4H7N3O | 113.0589 | 1 |
| Cyclic AMP | C10H12N5O6P | 329.0525 | 1 |
| Cytidine | C9H13N3O5 | 243.0855 | 1 |
| Cytidine 5'-monophosphate | C9H14N3O8P | 323.0519 | 1 |
| Cytosine | C4H5N3O | 111.0433 | 1 |
| D-3-Phosphoglyceric acid disodium salt | C3H7O7P | 185.9929 | 1 |
| D-Fructose 1,6-bisphosphate trisodium salt octahydrate | C6H14O12P2 | 339.9961 | 1 |
| D-Glucosamine 6-phosphate | C6H14NO8P | 259.0457 | 1 |

| | | | |
|---|---|---|---|
| D-Glucosamine hydrochloride | C6H13NO5 | 179.0794 | 1 |
| D-Glucose | C6H12O6 | 180.0634 | 1 |
| D-Glucose 6-phosphate sodium salt | C6H13O9P | 260.0297 | 1 |
| Dihydroxyacetone | C3H6O3 | 90.0317 | 1 |
| Dopamine hydrochloride | C8H11NO2 | 153.0790 | 1 |
| D-Ribose 5-phosphate disodium salt hydrate | C5H11O8P | 230.0192 | 1 |
| Epinephrine | C9H13NO3 | 183.0895 | 1 |
| Estradiol | C18H24O2 | 272.1776 | 1 |
| Flavin adenine dinucleotide disodium salt hydrate | C27H33N9O15P2 | 785.1571 | 1 |
| Flavin mononucleotide sodium salt | C17H21N4O9P | 456.1046 | 1 |
| Folic acid | C19H19N7O6 | 441.1397 | 1 |
| Fumaric acid | C4H4O4 | 116.0110 | 1 |
| gamma-Aminobutyric acid | C4H9NO2 | 103.0633 | 1 |
| Gluconic acid sodium salt | C6H12O7 | 196.0583 | 1 |
| Glycerophosphocholine | C8H20NO6P | 257.1028 | 1 |
| Glycine | C2H5NO2 | 75.0320 | 1 |
| Guanine | C5H5N5O | 151.0494 | 1 |
| Histamine dihydrochloride | C5H9N3 | 111.0796 | 1 |
| Homocysteine | C4H9NO2S | 135.0354 | 1 |
| Hypoxanthine | C5H4N4O | 136.0385 | 1 |
| Indole | C8H7N | 117.0578 | 1 |
| Kynurenic acid | C10H7NO3 | 189.0426 | 1 |
| Kynurenine | C10H12N2O3 | 208.0848 | 1 |
| L-4-Hydroxyproline | C5H9NO3 | 131.0582 | 1 |
| L-Acetylcarnitine hydrochloride | C9H17NO4 | 203.1158 | 1 |
| L-Arginine | C6H14N4O2 | 174.1117 | 1 |
| L-Asparagine | C4H8N2O3 | 132.0535 | 1 |
| L-Aspartic acid | C4H7NO4 | 133.0375 | 1 |
| L-Carnitine | C7H15NO3 | 161.1052 | 1 |
| L-Carnosine | C9H14N4O3 | 226.1066 | 1 |
| L-Citrulline | C6H13N3O3 | 175.0957 | 1 |
| L-Cysteine | C3H7NO2S | 121.0198 | 1 |
| L-Cystine | C6H12N2O4S2 | 240.0239 | 1 |
| L-Dihydroxyphenylalanine | C9H11NO4 | 197.0688 | 1 |
| L-Glutamic acid | C5H9NO4 | 147.0532 | 1 |
| L-Glutamine | C5H10N2O3 | 146.0691 | 1 |
| L-glutathione oxidised | C20H32N6O12S2 | 612.1520 | 1 |
| L-Glutathione reduced | C10H17N3O6S | 307.0838 | 1 |
| L-Histidine | C6H9N3O2 | 155.0695 | 1 |
| L-Lactic acid sodium salt | C3H6O3 | 90.0317 | 1 |
| L-Malic acid | C4H6O5 | 134.0215 | 1 |
| L-Methionine | C5H11NO2S | 149.0511 | 1 |
| L-Phenylalanine | C9H11NO2 | 165.0790 | 1 |
| L-Proline | C5H9NO2 | 115.0633 | 1 |

| | | | |
|---|---|---|---|
| L-Serine | C3H7NO3 | 105.0426 | 1 |
| L-Tryptophan | C11H12N2O2 | 204.0899 | 1 |
| L-Tyrosine | C9H11NO3 | 181.0739 | 1 |
| L-Valine | C5H11NO2 | 117.0790 | 1 |
| Malonic acid disodium salt | C3H4O4 | 104.0110 | 1 |
| Melatonin | C13H16N2O2 | 232.1212 | 1 |
| myo-Inositol | C6H12O6 | 180.0634 | 1 |
| N-Acetyl-D-glucosamine | C8H15NO6 | 221.0899 | 1 |
| N-Acetyl-L-aspartic acid | C6H9NO5 | 175.0481 | 1 |
| Niacinamide | C6H6N2O | 122.0480 | 1 |
| Nicotinamide adenine dinucleotide | C21H28N7O14P2 | 664.1170 | 1 |
| Nicotinamide adenine dinucleotide (reduced) disodium salt hydrate | C21H29N7O14P2 | 665.1248 | 1 |
| Nicotinic acid | C6H5NO2 | 123.0320 | 1 |
| Ornithine hydrochloride | C5H12N2O2 | 132.0899 | 1 |
| Orotic acid anhydrous | C5H4N2O4 | 156.0171 | 1 |
| Oxalacetic acid | C4H4O5 | 132.0059 | 1 |
| Palmitic acid | C16H32O2 | 256.2402 | 1 |
| Pantothenic acid sodium salt | C9H17NO5 | 219.1107 | 1 |
| Phosphoenolpyruvic acid potassium salt | C3H5O6P | 167.9824 | 1 |
| Phosphoserine | C3H8NO6P | 185.0089 | 1 |
| Phylloquinone | C31H46O2 | 450.3498 | 1 |
| Propionylcarnitine hydrochloride | C10H19NO4 | 217.1314 | 1 |
| Putrescine dihydrochloride | C4H12N2 | 88.1000 | 1 |
| Pyridoxine hydrochloride | C8H11NO3 | 169.0739 | 1 |
| Pyruvic acid | C3H4O3 | 88.0160 | 1 |
| Quinolinic acid | C7H5NO4 | 167.0219 | 1 |
| Riboflavin | C17H20N4O6 | 376.1383 | 1 |
| S-(5'-Adenosyl)-L-methionine p-toluenesulfonate salt | C15H23N6O5S | 399.1451 | 1 |
| Serotonin hydrochloride | C10H12N2O | 176.0950 | 1 |
| Spermidine | C7H19N3 | 145.1579 | 1 |
| Spermine tetrahydrochloride | C10H26N4 | 202.2157 | 1 |
| Succinic acid | C4H6O4 | 118.0266 | 1 |
| Taurine | C2H7NO3S | 125.0147 | 1 |
| Taurocholic acid sodium salt | C26H45NO7S | 515.2917 | 1 |
| Thiamine hydrochloride | C12H17N4OS | 265.1123 | 1 |
| Thymine | C5H6N2O2 | 126.0429 | 1 |
| trans-Urocanic acid | C6H6N2O2 | 138.0429 | 1 |
| Uracil | C4H4N2O2 | 112.0273 | 1 |
| Ureidosuccinic acid | C5H8N2O5 | 176.0433 | 1 |
| Uric acid | C5H4N4O3 | 168.0283 | 1 |
| Uridine | C9H12N2O6 | 244.0695 | 1 |
| Uridine 5'-diphosphate disodium salt hydrate | C9H14N2O12P2 | 404.0022 | 1 |
| Uridine diphosphate alpha-D-glucose disodium salt | C15H24N2O17P2 | 566.0550 | 1 |

| | | | |
|---|---|---|---|
| Xanthine | C5H4N4O2 | 152.0334 | 1 |
| (R)-Mevalonic acid lithium salt | C6H12O4 | 148.0736 | 2 |
| (R)-Pantetheine | C11H22N2O4S | 278.1300 | 2 |
| 3-Hydroxy-3-methylglutaric acid | C6H10O5 | 162.0528 | 2 |
| 3-Hydroxyanthranilic acid | C7H7NO3 | 153.0426 | 2 |
| 4-nitrobenzyl pyridinium chloride | C12H11N2O2 | 215.0821 | 2 |
| 5-Methyltetrahydrofolic acid disodium salt | C20H25N7O6 | 459.1866 | 2 |
| 6-Phosphogluconic acid trisodium salt | C6H13O10P | 276.0246 | 2 |
| Acetoacetic acid lithium salt | C4H6O3 | 102.0317 | 2 |
| Acetyl-CoA trisodium salt | C23H38N7O17P3S | 809.1258 | 2 |
| Arachidonic acid | C20H32O2 | 304.2402 | 2 |
| Argininosuccinic acid lithium salt | C10H18N4O6 | 290.1226 | 2 |
| Cholesteryl acetate | C29H48O2 | 428.3654 | 2 |
| Dihydroxyacetone phosphate dilithium salt | C3H7O6P | 169.9980 | 2 |
| D-Ribulose 5-phosphate disodium salt | C5H11O8P | 230.0192 | 2 |
| L-Ribulose | C5H10O5 | 150.0528 | 2 |
| myo-Inositol 1-phosphate dipotassium salt | C6H13O9P | 260.0297 | 2 |
| N6,N6,N6-Trimethyl-L-lysine hydrochloride | C9H20N2O2 | 188.1525 | 2 |
| N-Acetylaspartylglutamic acid | C11H16N2O8 | 304.0907 | 2 |
| N-Acetyl-D-galactosamine 1-phosphate | C8H16NO9P | 301.0563 | 2 |
| Nicotinamide riboside chloride | C11H15N2O5 | 255.0981 | 2 |
| Nicotinamide ribotide | C11H15N2O8P | 334.0566 | 2 |
| S-(5'-Adenosyl)-L-homocysteine | C14H20N6O5S | 384.1216 | 2 |
| sn-Glycerol 3-phosphate lithium salt | C3H9O6P | 172.0137 | 2 |
| Uridine 5'-diphosphoglucuronic acid trisodium salt | C15H22N2O18P2 | 580.0343 | 2 |
| Uridine 5'-diphospho-N-acetylglucosamine disodium salt | C17H27N3O17P2 | 607.0816 | 2 |
| 1-(10Z-heptadecenoyl)-2-hydroxy-sn-glycero-3-[phospho-L-serine] (sodium salt) | C23H44NO9P | 509.2754 | 3 |
| 1-(10Z-heptadecenoyl)-2-hydroxy-sn-glycero-3-phospho-(1'-myo-inositol) (ammonium salt) | C26H49O12P | 584.2962 | 3 |
| 1,2-dioleoyl-sn-glycero-3-phospho-(1'-myo-inositol-5'-phosphate) (diammonium salt) | C45H84O16P2 | 942.5235 | 3 |
| 1',3'-bis[1,2-dioleoyl-sn-glycero-3-phospho]-glycerol (disodium salt) | C81H150O17P2 | 1457.0350 | 3 |
| 1,3-dipentadecanoyl-2-oleoyl-glycerol | C51H96O6 | 804.7207 | 3 |
| 1-O-hexadecyl-2-oleoyl-sn-glycero-3-phosphocholine | C42H84NO7P | 745.5985 | 3 |
| 1-oleoyl-2-hydroxy-sn-glycero-3-phosphate (sodium salt) | C21H41O7P | 436.2590 | 3 |
| 1-oleoyl-rac-glycerol | C21H40O4 | 356.2927 | 3 |
| 1-palmitoyl-2-hydroxy-sn-glycero-3-phospho-(1'-rac-glycerol) (sodium salt) | C22H45O9P | 484.2801 | 3 |
| 1-palmitoyl-2-oleoyl-sn-glycero-3-phosphate (sodium salt) | C37H71O8P | 674.4887 | 3 |
| 1-palmitoyl-2-oleoyl-sn-glycero-3-phospho-(1'-rac-glycerol) (sodium salt) | C40H77O10P | 748.5254 | 3 |

| | | | |
|---|---|---|---|
| 1-palmitoyl-2-oleoyl-sn-glycero-3-phosphoinositol (ammonium salt) | C43H81O13P | 836.5415 | 3 |
| 1-palmitoyl-2-oleoyl-sn-glycero-3-phospho-L-serine (sodium salt) | C40H76NO10P | 761.5207 | 3 |
| 1-pentadecanoyl-2-hydroxy-sn-glycero-3-phosphoch oline | C23H48NO7P | 481.3168 | 3 |
| 1-pentadecanoyl-2-oleoyl-sn-glycero-3-phosphocholi ne | C41H80NO8P | 745.5622 | 3 |
| 1-stearoyl-2-arachidonoyl-sn-glycero-3-phosphoetha nolamine | C43H78NO8P | 767.5465 | 3 |
| 1-stearoyl-2-docosahexaenoyl-sn-glycerol | C43H72O5 | 668.5380 | 3 |
| 1-stearoyl-2-hydroxy-sn-glycero-3-phosphoethanola mine | C23H48NO7P | 481.3168 | 3 |
| 3-O-sulfo-D-galactosyl-beta1-1'-N-[2"(S)-hydroxystea royl]-D-erythro-sphingosine (ammonium salt), (synthetic) | C42H81NO12S | 823.5480 | 3 |
| beta-D-glucosyl cholesterol | C33H56O6 | 548.4077 | 3 |
| C18:0 GM3 Ganglioside (synthetic, ammonium salt) | C59H108N2O21 | 1180.7450 | 3 |
| Coenzyme A | C21H36N7O16P3S | 767.1152 | 3 |
| D-glucosyl-beta-1,1'-N-stearoyl-D-erythro-sphingosin e | C42H81NO8 | 727.5962 | 3 |
| N-behenoyl-D-erythro-sphingosine | C40H79NO3 | 621.6060 | 3 |
| N-palmitoyl-D-erythro-sphingosylphosphorylcholine | C39H79N2O6P | 702.5676 | 3 |
| Palmitoyl L-carnitine | C23H45NO4 | 399.3349 | 3 |
| Palmitoyl-CoA triammonium salt | C37H66N7O17P3S | 1005.3450 | 3 |
| Prostaglandin E1 | C20H34O5 | 354.2406 | 3 |
| 4-chlorobenzyl pyridinium chloride | C12H11ClN | 204.0580 | 4 |
| 4-cyanobenzyl pyridinium chloride | C20H17N2 | 285.1392 | 4 |
| 4-methoxybenzyl pyridinium chloride | C13H14NO | 200.1075 | 4 |
| 4-methylbenzyl pyridinium chloride | C13H14N | 184.1126 | 4 |

Compound metadata including both vendor name and short name can be found in **Supplementary Data 4**

#### Supplementary Table 2

Preparation protocols used for metabolite standards.

Diluent solvent systems:

- M:W - 9:1 Methanol/Water (v/v)
- M:C - 1:2 Methanol/Chloroform (v/v).

Titrant shorthand:

- NaOH – Sodium Hydroxide (aqueous)
- HCl – Hydrochloric Acid (aqueous)

| Name | Stock (mM) | Stock Solvent | Dilution (μM) | Diluent | Titrant |
|---|---|---|---|---|---|
| D-Glucose | 98.3 | Water | 200 | M:W | |
| D-Glucose 6-phosphate sodium salt | 39.0 | Water | 200 | M:W | |
| D-Fructose 1,6-bisphosphate trisodium salt octahydrate | 79.8 | Water | 200 | M:W | |
| Dihydroxyacetone phosphate dilithium salt | 5.5 | Water | 200 | M:W | |
| D-3-Phosphoglyceric acid disodium salt | 21.4 | Water | 200 | M:W | |
| Phosphoenolpyruvic acid potassium salt | 63.7 | Water | 200 | M:W | |
| Pyruvic acid | 211.4 | Water | 200 | M:W | |
| Acetyl-CoA trisodium salt | 1.1 | Water | 200 | M:W | |
| L-Lactic acid sodium salt | 93.6 | Water | 200 | M:W | |
| 6-Phosphogluconic acid trisodium salt | 2.9 | Water | 200 | M:W | |
| D-Ribulose 5-phosphate disodium salt | 3.6 | Water | 200 | M:W | |
| D-Ribose 5-phosphate disodium salt hydrate | 40.8 | Water | 200 | M:W | |
| Dihydroxyacetone | 169.8 | Water | 200 | M:W | |
| Gluconic acid sodium salt | 87.6 | Water | 200 | M:W | |
| L-Ribulose | 199.8 | Water | 200 | M:W | |
| Citric acid | 79.1 | Water | 200 | M:W | |
| cis-Aconitic acid | 76.5 | Water | 200 | M:W | |
| 2-Oxoglutaric acid | 158.8 | Water | 200 | M:W | |
| Succinic acid | 124.7 | Water | 200 | M:W | |
| Fumaric acid | 137.0 | Methanol | 200 | M:C | |
| L-Malic acid | 79.1 | Water | 200 | M:W | |
| Oxalacetic acid | 77.2 | Water | 200 | M:W | |
| Uridine diphosphate alpha-D-glucose disodium salt | 8.2 | Water | 200 | M:W | |
| Uridine 5'-diphosphoglucuronic acid trisodium salt | 1.5 | Water | 200 | M:W | |
| D-Glucosamine 6-phosphate | 20.4 | Water | 200 | M:W | |

| | | | | | |
|---|---|---|---|---|---|
| N-Acetyl-D-galactosamine 1-phosphate | 3.3 | Water | 200 | M:W | |
| Uridine 5'-diphospho-N-acetylglucosamine disodium salt | 5.4 | Water | 200 | M:W | |
| N-Acetyl-D-glucosamine | 51.0 | Water | 200 | M:W | |
| myo-Inositol | 174.8 | Water | 200 | M:W | |
| Ascorbic acid | 98.2 | Water | 200 | M:W | |
| myo-inositol 1-phosphate dipotassium salt | 28.5 | Water | 200 | M:W | |
| Palmitic acid | 67.5 | Methanol | 200 | M:C | |
| (R)-Mevalonic acid lithium salt | 6.5 | Water | 200 | M:W | |
| Cholesterol | 26.9 | 1:5 Chloroform/Methanol | 200 | M:C | |
| Malonic acid disodium salt | 33.3 | Water | 200 | M:W | |
| 3-Hydroxy-3-methylglutaric acid | 6.2 | Water | 200 | M:W | |
| Palmitoyl-CoA triammonium salt | 4.7 | 2:1:1 MeOH/H2O/CHCl3 | 200 | M:C | |
| L-Carnitine | 93.6 | Water | 200 | M:W | |
| Cholic acid | 46.7 | Methanol | 200 | M:C | |
| Glycine | 251.8 | Water | 200 | M:W | |
| Taurocholic acid sodium salt | 26.8 | 40% Methanol | 200 | M:W | |
| Estradiol | 40.8 | Methanol | 200 | M:C | |
| L-Asparagine | 160.3 | 20% Methanol | 200 | M:W | HCl |
| L-Aspartic acid | 119.6 | 20% Methanol | 200 | M:W | HCl |
| N-Acetyl-L-aspartic acid | 100.7 | Water | 200 | M:W | |
| L-Glutamine | 109.5 | Water | 200 | M:W | |
| L-Glutamic acid | 201.8 | 20% Methanol | 200 | M:W | HCl |
| N-Acetylaspartylglutamic acid | 3.3 | Water | 200 | M:W | |
| gamma-Aminobutyric acid | 164.9 | Water | 200 | M:W | |
| L-Phenylalanine | 110.4 | 20% Methanol | 200 | M:W | HCl |
| L-Tyrosine | 112.0 | 20% Methanol | 200 | M:W | HCl |
| L-Dihydroxyphenylalanine | 99.9 | 20% Methanol | 200 | M:W | HCl |
| Dopamine hydrochloride | 69.6 | Water | 200 | M:W | |
| Epinephrine | 47.3 | 90% Methanol | 200 | M:W | NaOH |
| Acetoacetic acid lithium salt | 9.3 | Water | 200 | M:W | |
| L-Histidine | 92.2 | Water | 200 | M:W | |
| L-Carnosine | 74.3 | Water | 200 | M:W | |
| trans-Urocanic acid | 97.4 | 40% Methanol | 200 | M:W | NaOH |
| Histamine dihydrochloride | 97.8 | Water | 200 | M:W | |
| L-Arginine | 109.1 | Water | 200 | M:W | |
| L-Citrulline | 134.1 | Water | 200 | M:W | |
| Argininosuccinic acid lithium salt | 3.4 | Water | 200 | M:W | |
| Ornithine hydrochloride | 102.0 | Water | 200 | M:W | |
| Carbamoyl phosphate dilithium salt hydrate | 63.4 | Water | 200 | M:W | |
| L-Proline | 271.0 | Water | 200 | M:W | |
| L-4-Hydroxyproline | 176.2 | Water | 200 | M:W | |

| | | | | | |
|---|---|---|---|---|---|
| Putrescine dihydrochloride | 87.5 | Water | 200 | M:W | |
| Spermidine | 477.8 | Water | 200 | M:W | |
| Spermine tetrahydrochloride | 33.0 | Water | 200 | M:W | |
| Creatine anhydrous | 117.4 | Water | 200 | M:W | |
| Creatinine | 282.0 | 40% Methanol | 200 | M:W | |
| Agmatine sulfate | 55.6 | Water | 200 | M:W | |
| Folic acid | 45.8 | 40% Methanol | 200 | M:W | NaOH |
| 5-Methyltetrahydrofolic acid disodium salt | 2.0 | 40% Methanol | 200 | M:W | |
| L-Methionine | 179.6 | 20% Methanol | 200 | M:W | HCl |
| L-Cystine | 142.3 | 20% Methanol | 200 | M:W | HCl |
| L-glutathione oxidised | 24.3 | Water | 200 | M:W | |
| L-Glutathione reduced | 36.7 | Water | 200 | M:W | |
| Taurine | 112.3 | Water | 200 | M:W | |
| S-(5'-Adenosyl)-L-methionine p-toluenesulfonate salt | 27.0 | Water | 200 | M:W | |
| S-(5'-Adenosyl)-L-homocysteine | 2.6 | Water | 200 | M:W | |
| Homocysteine | 102.8 | Water | 200 | M:W | |
| L-Cysteine | 203.0 | Water | 200 | M:W | HCl |
| N6,N6,N6-Trimethyl-L-lysine hydrochloride | 9.4 | Water | 200 | M:W | |
| L-Valine | 398.6 | Water | 200 | M:W | |
| Pantothenic acid sodium salt | 55.6 | Water | 200 | M:W | |
| (R)-Pantetheine | 3.6 | Methanol | 200 | M:W | |
| Coenzyme A | 3.0 | Water | 200 | M:W | |
| L-Serine | 211.2 | Water | 200 | M:W | |
| L-Tryptophan | 96.0 | Water | 200 | M:W | HCl |
| Kynurenine | 24.0 | Water | 200 | M:W | HCl |
| Kynurenic acid | 43.0 | 1:1 Methanol/Water | 200 | M:W | NaOH |
| 3-Hydroxyanthranilic acid | 6.5 | Methanol | 200 | M:W | |
| Quinolinic acid | 118.5 | 20% Methanol | 200 | M:W | NaOH |
| Nicotinamide adenine dinucleotide | 15.5 | Water | 200 | M:W | |
| Nicotinamide adenine dinucleotide (reduced) disodium salt hydrate | 15.1 | Water | 200 | M:W | |
| Niacinamide | 147.4 | Water | 200 | M:W | |
| Nicotinic acid | 139.2 | 95% Methanol | 200 | M:W | NaOH |
| Serotonin hydrochloride | 111.4 | Water | 200 | M:W | |
| 5-Hydroxyindole-3-acetic acid | 68.0 | Ethanol | 200 | M:W | |
| Melatonin | 111.5 | Methanol | 200 | M:C | |
| Nicotinamide riboside chloride | 3.4 | Water | 200 | M:W | |
| Nicotinamide ribotide | 3.0 | Water | 200 | M:W | |
| Hypoxanthine | 62.4 | Water | 200 | M:W | NaOH |
| Adenosine 5'-monophosphate | 60.3 | Water | 200 | M:W | NaOH |
| Adenine hydrochloride | 57.9 | Water | 200 | M:W | |
| Xanthine | 84.8 | 10% Methanol | 200 | M:W | NaOH |

| | | | | | |
|---|---|---|---|---|---|
| Guanine | 91.5 | Water | 200 | M:W | NaOH |
| Adenosine 5'-diphosphate potassium salt | 30.9 | Water | 200 | M:W | |
| Adenosine 5'-triphosphate disodium salt | 55.3 | Water | 200 | M:W | |
| Cyclic AMP | 24.1 | Water | 200 | M:W | |
| Ureidosuccinic acid | 72.3 | Water | 200 | M:W | |
| 4,5-Dihydroorotic acid | 105.0 | Water | 200 | M:W | |
| Orotic acid anhydrous | 67.0 | 40% Methanol | 200 | M:W | NaOH |
| Uridine | 100.3 | Water | 200 | M:W | |
| Uracil | 161.5 | Water | 200 | M:W | NaOH |
| Thymine | 119.7 | Water | 200 | M:W | NaOH |
| Cytidine 5'-monophosphate | 60.7 | Water | 200 | M:W | |
| Cytidine | 42.3 | Water | 200 | M:W | |
| Cytosine | 106.3 | Water | 200 | M:W | HCl |
| Uridine 5'-diphosphate disodium salt hydrate | 72.5 | Water | 200 | M:W | |
| Choline chloride | 208.4 | Water | 200 | M:W | |
| Acetylcholine chloride | 239.5 | Water | 200 | M:W | |
| Thiamine hydrochloride | 47.1 | Water | 200 | M:W | |
| Riboflavin | 27.4 | Water | 200 | M:W | NaOH |
| Pyridoxine hydrochloride | 56.4 | Water | 200 | M:W | |
| Biotin | 59.3 | 40% Methanol | 200 | M:W | NaOH |
| Cholecalciferol | 76.7 | Methanol | 200 | M:C | |
| alpha-Tocopherol | 165.4 | Methanol | 200 | M:C | |
| Phylloquinone | 163.7 | Chloroform | 200 | M:C | |
| Flavin mononucleotide sodium salt | 22.4 | Water | 200 | M:W | |
| Flavin adenine dinucleotide disodium salt hydrate | 13.8 | Water | 200 | M:W | |
| Creatine phosphate disodium salt hydrate | 57.2 | Water | 200 | M:W | |
| Glycerophosphocholine | 47.4 | Water | 200 | M:W | |
| Propionylcarnitine hydrochloride | 75.7 | 40% Methanol | 200 | M:W | |
| L-Acetylcarnitine hydrochloride | 51.1 | Water | 200 | M:W | |
| Phosphoserine | 55.4 | Water | 200 | M:W | |
| Uric acid | 81.3 | Water | 200 | M:W | NaOH |
| Cholesteryl acetate | 5.3 | Chloroform | 200 | M:C | |
| Indole | 126.3 | Methanol | 200 | M:C | |
| D-Glucosamine hydrochloride | 80.7 | Water | 200 | M:W | |
| Butyric acid sodium salt | 193.5 | Water | 200 | M:W | |
| sn-Glycerol 3-phosphate lithium salt | 5.8 | Water | 200 | M:W | |
| all-trans-Retinoic acid | 45.9 | Chloroform | 200 | M:C | |
| Arachidonic acid | 3.3 | Chloroform | 200 | M:C | |
| 1-pentadecanoyl-2-oleoyl-sn-glycero-3-phosphocholine | 1.3 | Chloroform | 200 | M:C | |

| | | | | |
|---|---|---|---|---|
| 1,3-dipentadecanoyl-2-oleoyl-glycerol | 1.2 | Chloroform | 200 | M:C |
| 1-oleoyl-rac-glycerol | 2.8 | Chloroform | 200 | M:C |
| beta-D-glucosyl cholesterol | 3.0 | 2:1 Chloroform/Methanol | 200 | M:C |
| 1',3'-bis[1,2-dioleoyl-sn-glycero-3-phospho]-glycerol (disodium salt) | 16.6 | Chloroform | 200 | M:C |
| 1-stearoyl-2-docosahexaenoyl-sn-glycerol | 3.0 | Chloroform | 200 | M:C |
| 1-palmitoyl-2-oleoyl-sn-glycero-3-phospho-L-serine (sodium salt) | 12.8 | Chloroform | 200 | M:C |
| 1-palmitoyl-2-oleoyl-sn-glycero-3-phospho-(1'-rac-glycerol) (sodium salt) | 13.0 | Chloroform | 200 | M:C |
| 1-palmitoyl-2-oleoyl-sn-glycero-3-phosphate (sodium salt) | 35.9 | Chloroform | 200 | M:C |
| 1-(10Z-heptadecenoyl)-2-hydroxy-sn-glycero-3-phospho-(1'-myo-inositol) (ammonium salt) | 0.2 | 1:1 Chloroform/Methanol | 50 | M:C |
| 1-palmitoyl-2-oleoyl-sn-glycero-3-phosphoinositol (ammonium salt) | 0.1 | Chloroform | 50 | M:C |
| 1,2-dioleoyl-sn-glycero-3-phospho-(1'-myo-inositol-5'-phosphate) (diammonium salt) | 0.2 | Chloroform | 50 | M:C |
| 1-stearoyl-2-arachidonoyl-sn-glycero-3-phosphoethanolamine | 13.0 | Chloroform | 200 | M:C |
| 1-pentadecanoyl-2-hydroxy-sn-glycero-3-phosphocholine | 8.5 | 1:1 Chloroform/Methanol | 200 | M:C |
| 1-Stearoyl-2-Hydroxy-sn-Glycero-3-Phosphoethanolamine | 1.1 | 1:1 Chloroform/Methanol | 200 | M:C |
| 1-oleoyl-2-hydroxy-sn-glycero-3-phosphate (sodium salt) | 10.9 | Chloroform | 200 | M:C |
| 1-palmitoyl-2-hydroxy-sn-glycero-3-phospho-(1'-rac-glycerol) (sodium salt) | 8.7 | Methanol | 200 | M:C |
| 1-(10Z-heptadecenoyl)-2-hydroxy-sn-glycero-3-[phospho-L-serine] (sodium salt) | 4.7 | 1:1 Chloroform/Methanol | 200 | M:C |
| C18:0 GM3 Ganglioside (synthetic, ammonium salt) | 0.1 | Methanol | 10 | M:C |
| N-behenoyl-D-erythro-sphingosine | 2.7 | 2:1 Chloroform/Methanol | 200 | M:C |
| D-glucosyl-beta-1,1'-N-stearoyl-D-erythro-sphingosine | 6.9 | 1:1 Chloroform/Methanol | 200 | M:C |
| N-palmitoyl-D-erythro-sphingosylphosphorylcholine | 7.1 | Chloroform | 200 | M:C |
| 3-O-sulfo-D-galactosyl-beta1-1'-N-[2"(S)-hydroxystearoyl]-D-erythro-sphingosine (ammonium salt), (synthetic) | 1.2 | Chloroform | 200 | M:C |
| palmitoyl L-carnitine | 6.3 | Methanol | 200 | M:C |
| 1-O-hexadecyl-2-oleoyl-sn-glycero-3-phosphocholine | 3.4 | Chloroform | 200 | M:C |
| Prostaglandin E1 | 1.4 | 1:1 Chloroform/methanol | 200 | M:C |
| Cholesteryl heptadecanoate | 11.9 | 1:1 Methanol/Chloroform | 200 | M:C |
| 4-nitrobenzyl pyridinium chloride | 1.4 | Methanol | 200 | M:C |
| 4-methoxybenzyl pyridinium chloride | 0.9 | Methanol | 200 | M:C |

| | | | | |
|---|---|---|---|---|
| 4-methylbenzyl pyridinium chloride | 4.3 | Methanol | 200 | M:C |
| 4-cyanobenzyl pyridinium chloride | 0.6 | Methanol | 200 | M:C |
| 4-chlorobenzyl pyridinium chloride | 4.1 | Methanol | 200 | M:C |

#### Supplementary Table 3

List of metabolites with index number (key to layout)

| Index | Name |
|---|---|
| 0 | Glucose |
| 1 | Glucose 6-phosphate |
| 3 | Fructose 1,6-bisphosphate |
| 4 | Dihydroxyacetone phosphate |
| 7 | 3-Phosphoglyceric acid |
| 9 | Phosphoenolpyruvic acid |
| 10 | Pyruvic acid |
| 11 | Acetyl-CoA |
| 12 | Lactic acid |
| 14 | 6-Phosphogluconic acid |
| 15 | Ribulose 5-phosphate |
| 16 | Ribose 5-phosphate |
| 17 | Dihydroxyacetone |
| 20 | Gluconic acid |
| 21 | Ribulose |
| 22 | Citric acid |
| 24 | cis-Aconitic acid |
| 26 | 2-Oxoglutaric acid |
| 28 | Succinic acid |
| 29 | Fumaric acid |
| 30 | Malic acid |
| 31 | Oxalacetic acid |
| 33 | Uridine diphosphate glucose |
| 34 | Uridine diphosphate glucuronic acid |
| 35 | Glucosamine 6-phosphate |
| 36 | N-Acetylgalactosamine 1-phosphate |
| 38 | Uridine diphosphate-N-acetylglucosamine |
| 39 | N-Acetylglucosamine |
| 40 | myo-Inositol |
| 41 | Ascorbic acid |
| 42 | myo-Inositol 1-phosphate |
| 46 | Palmitic acid |
| 50 | Mevalonic acid |
| 51 | Cholesterol |

| | |
|---|---|
| 53 | Malonic acid |
| 54 | 3-Hydroxymethylglutaric acid |
| 55 | Palmitoyl-CoA |
| 56 | Carnitine |
| 58 | Cholic acid |
| 59 | Glycine |
| 60 | Taurocholic acid |
| 61 | Estradiol |
| 67 | Asparagine |
| 68 | Aspartic acid |
| 69 | N-Acetylaspartic acid |
| 70 | Glutamine |
| 71 | Glutamic acid |
| 72 | N-Acetylaspartylglutamic acid |
| 73 | gamma-Aminobutyric acid |
| 74 | Phenylalanine |
| 75 | Tyrosine |
| 76 | L-Dopa |
| 77 | Dopamine |
| 79 | Epinephrine |
| 83 | Acetoacetic acid |
| 84 | Histidine |
| 85 | Carnosine |
| 86 | Urocanic acid |
| 89 | Histamine |
| 90 | Arginine |
| 91 | Citrulline |
| 92 | Argininosuccinic acid |
| 93 | Ornithine |
| 95 | Carbamoyl phosphate |
| 97 | Proline |
| 98 | 4-Hydroxyproline |
| 99 | Putrescine |
| 100 | Spermidine |
| 101 | Spermine |
| 103 | Creatine |
| 105 | Creatinine |
| 106 | Agmatine |
| 107 | Folic acid |
| 108 | Tetrahydrofolic acid |
| 110 | Methionine |
| 111 | Cystine |
| 113 | Oxidized glutathione |
| 114 | Glutathione |

| | |
|---|---|
| 117 | Taurine |
| 119 | S-Adenosylmethionine |
| 120 | S-Adenosylhomocysteine |
| 121 | Homocysteine |
| 123 | Cysteine |
| 125 | N6,N6,N6-Trimethyllysine |
| 129 | Valine |
| 132 | Pantothenic acid |
| 136 | Pantetheine |
| 138 | Coenzyme A |
| 141 | Serine |
| 148 | Tryptophan |
| 150 | Kynurenine |
| 151 | Kynurenic acid |
| 154 | 3-Hydroxyanthranilic acid |
| 155 | Quinolinic acid |
| 158 | Nicotinamide adenine dinucleotide (NAD) |
| 159 | Nicotinamide adenine dinucleotide, reduced (NADH) |
| 160 | Niacinamide |
| 161 | Nicotinic acid |
| 165 | Serotonin |
| 166 | 5-Hydroxyindoleacetic acid |
| 168 | Melatonin |
| 169 | Nicotinamide riboside |
| 170 | Nicotinamide ribotide |
| 184 | Hypoxanthine |
| 185 | Adenosine monophosphate |
| 187 | Adenine |
| 190 | Xanthine |
| 193 | Guanine |
| 194 | Adenosine diphosphate |
| 195 | Adenosine triphosphate |
| 198 | Cyclic adenosine monophosphate |
| 204 | Ureidosuccinic acid |
| 205 | 4,5-Dihydroorotic acid |
| 206 | Orotic acid |
| 209 | Uridine |
| 210 | Uracil |
| 213 | Thymine |
| 214 | Cytidine monophosphate |
| 215 | Cytidine |
| 216 | Cytosine |
| 217 | Uridine diphosphate |
| 226 | Choline |

| | |
|---|---|
| 227 | Acetylcholine |
| 231 | Thiamine |
| 232 | Riboflavin |
| 233 | Pyridoxine |
| 235 | Biotin |
| 236 | Cholecalciferol |
| 237 | alpha-Tocopherol |
| 238 | Phylloquinone |
| 239 | Flavin mononucleotide |
| 241 | Flavin adenine dinucleotide |
| 246 | Phosphocreatine |
| 250 | Glycerophosphocholine |
| 254 | Propionylcarnitine |
| 257 | Acetylcarnitine |
| 264 | Phosphoserine |
| 274 | Uric acid |
| 276 | Cholesteryl acetate |
| 310 | Indole |
| 318 | Glucosamine |
| 322 | Butyric acid |
| 323 | Glycerol 3-phosphate |
| 326 | Retinoic acid |
| 501 | Arachidonic acid |
| 502 | PC 15:0-18:1 |
| 503 | TG 15:0-18:1-15:0 |
| 504 | MG 18:1 |
| 505 | Glucosyl cholesterol |
| 506 | Cardiolipin 18:1 |
| 508 | DG 18:0-22:6 |
| 509 | PS (POPS) 16:0-18:1 |
| 510 | PG 16:0-18:1 |
| 511 | PA 16:0-18:1 |
| 512 | Lyso PI 17:1 |
| 513 | PI 16:0-18:1 |
| 514 | PI(5)P 18:1 |
| 515 | PE 18:0-20:4 |
| 516 | Lyso PC 15:0 |
| 517 | Lyso PE 18:0 |
| 518 | Lyso PA 18:1 |
| 519 | Lyso PG 16:0 |
| 520 | Lyso PS 17:1 |
| 521 | GM3 Ganglioside 18:0 |
| 522 | Cer d18:1-22:0 |
| 523 | GlcCer d18:1-18:0 |

| | |
|---|---|
| 524 | SM d18:1-16:0 |
| 525 | Sulfo GalCer 18:0(2S-OH) |
| 526 | CAR 16:0 |
| 527 | PC (POPC) 16:0-18:1 |
| 528 | Prostaglandin E1 |
| 529 | Cholesteryl ester 17:0 |
| 901 | 4-nitrobenzyl pyridinium chloride |
| 902 | 4-methoxybenzyl pyridinium chloride |
| 903 | 4-methylbenzyl pyridinium chloride |
| 904 | 4-cyanobenzyl pyridinium chloride |
| 905 | 4-chlorobenzyl pyridinium chloride |
| 906-908 | Rhodamine B + Caffeine (calibration spots) |
| 900 | Blank |

#### Supplementary Table 4

List of matrix substances used with abbreviated and full names and application examples.

| MALDI matrix shorthand | Chemical name | Typical polarity | Applications |
|---|---|---|---|
| DHB | 2,5-dihydroxybenzoic acid | Positive | Amino acids, lipids, peptides[4] |
| DHAP | 2,5-dihydroxybenzophenone | Positive | Phospholipids[5] |
| CHCA | α-cyano-4-hydroxycinnamic acid | Positive | Proteins, LMWC[4] |
| ClCCA | 4-chloro-α-cyanocinnamic acid | Positive | Peptides[6] |
| CMBT | 5-chloro-2-mercaptobenzothiazole | Positive | Peptides, oligosaccharides, glycolipids[7], lipids[8] |
| 9AA | 9-aminoacridine | Negative | Peptides, oligonucleotides, acidic LMWC (carboxylic acids, sulphonates, amines)[9], lipids[10] |
| NEDC | N-(1-naphthyl)ethylenediamine dihydrochloride | Negative | Glycerolipids, LMWC[11] |
| MAPS | 4-maleicanhydridoproton sponge | Negative | LMWC[12] |
| DAN | 1,5-diaminonaphthalene | Dual-mode | Lipids[13], LMWC[14] |
| pNA | para-nitroaniline | Dual-mode | Proteins, nucleotides, lipids[15] |
| NOR | norharmane | Dual-mode | Lipids[16] |
| PNDI | naphthalene diimide-co-thiophene (polymer) | Dual-mode | LMWC[17] |

LMWC = low molecular weight compounds

#### Supplementary Table 5

TM Sprayer Protocols. Matrices are listed with abbreviated names. The methods were adjusted to yield the same matrix-to-analyte ratio. In the case of PNDI, the monomer unit was used for estimating molar density.

| Matrix | Solvent System | Conc (mg/mL) | Temp (°C) | Flow rate (mL/min) | Gas Pressure (psi) | Track Spacing (mm) | Velocity (mm/min) | Passes | Density ($mg/mm^2$) |
|---|---|---|---|---|---|---|---|---|---|
| DAN | 7:3 ACN:H2O | 10 | 85 | 0.07 | 10 | 3 | 1350 | 8 | 0.001383 |
| DHB | 7:3 ACN:H2O | 10 | 85 | 0.068 | 10 | 3 | 1350 | 8 | 0.001343 |

| | | | | | | | | | |
|---|---|---|---|---|---|---|---|---|---|
| NOR | 1:1 CHCl3:MeOH | 7 | 30 | 0.122 | 10 | 3 | 1350 | 7 | 0.001476 |
| 9AA | 7:3 MeOH:H2O | 5 | 75 | 0.07 | 10 | 3 | 1350 | 20 | 0.001728 |
| CHCA | 1:1 ACN:H2O | 7 | 90 | 0.071 | 10 | 2 | 1350 | 9 | 0.001657 |
| PNDI | Toluene | 2.5 | 30 | 0.1 | 10 | 3 | 600 | 10 | 0.001389 |
| MAPS | Toluene | 5 | 30 | 0.1 | 10 | 3 | 600 | 10 | 0.002778 |
| NEDC | 7:3 ACN:H2O | 10 | 80 | 0.05 | 10 | 3 | 1350 | 13 | 0.001605 |
| DHAP | 7:3 ACN:H2O | 10 | 85 | 0.067 | 10 | 3 | 1350 | 8 | 0.001323 |
| ClCCA | 1:1 ACN:H2O | 7 | 90 | 0.07 | 10 | 2 | 1350 | 10 | 0.001815 |
| CMBT | 9:1 ACN:H2O | 7 | 70 | 0.1 | 10 | 3 | 1350 | 10 | 0.001728 |
| pNA | 17:3 MeOH:H2O | 11.5 | 65 | 0.068 | 10 | 3 | 1350 | 6 | 0.001153 |

#### Supplementary Table 6

Instrument method used for AP-MALDI-Orbitrap imaging

| Parameter | Low Mass Range | High Mass Range |
|---|---|---|
| Attenuator Angle* | 29° | 29° |
| Pitch (µm) | 150 | 150 |
| Mass Range (*m/z*) | 70-350 | 300-1510 |
| Voltage (kV) | +3 / -3.1 | +3 / -3.1 |
| RF Level | 50 | 100 |
| Injection Time (ms) | 500 | 500 |
| Mass Resolving Power (nominal) | 140000 | 140000 |

*Individual AP-SMALDI5 instruments require different attenuator settings to reach the same results, as the precise laser energy output onto the sample depends not only on the attenuator but also on the exact beam path through the optics, which can vary slightly between instruments.

#### Supplementary Table 7

Experimental parameters for all interlaboratory survey data sets, in positive (+) and negative (-) ionisation modes. Further metadata and METASPACE links to browse and download the data can be accessed through the digital supplement (Sup. Table 8, )

| Slide (+/-) | Source | Analyser | Pressure | Matrix (+/-) | Substrate |
|---|---|---|---|---|---|
| 1A/1C | DESI | Orbitrap | Ambient | None | Glass |
| 13G/13J | MALDI | qTOF | Vacuum | DHB/NEDC | ITO |
| 2I/7B | MALDI | qTOF | Vacuum | DHAP | Glass |
| 4F/3A | MALDI-2 | Orbitrap | Ambient | DHAP | Glass |
| 7J/7A | AP-MALDI | Orbitrap | Ambient | DHB/DAN | Glass |
| 12C/12E | MALDI | FTICR | Vacuum | DHB/DAN | ITO |
| 12G/12I | MALDI | qTOF | Vacuum | DHB/DAN | ITO |
| 8A/8D | AP-MALDI | Orbitrap | Ambient | DHB/9AA | Glass |
| 14B/14D | AP-MALDI | Orbitrap | Ambient | sDHB/9AA | Glass |
| 9A/9E | MALDESI | Orbitrap | Ambient | water | Glass |
| 10I/10G | MALDI | FTICR | Vacuum | DHB/NEDC | ITO |
| 5HJ/5GI | DESI | Orbitrap | Ambient | None | Glass |
| 1J/1H | DESI | Orbitrap | Ambient | None | Glass |

#### Supplementary Table 8

List of datasets with links to METASPACE.
Full project URL: https://metaspace2020.eu/project/Saharuka-2023

| Dataset | METASPACE URL |
|---|---|
| 15C EMBL AP-MALDI-Orbitrap pNA negative 3ppm | https://metaspace2020.eu/annotations?db_id=304&ds=2021-12-07_17h42m27s |
| 15C EMBL AP-MALDI-Orbitrap pNA positive 3ppm | https://metaspace2020.eu/annotations?db_id=304&ds=2021-12-07_17h42m06s |
| 3C EMBL AP-MALDI-Orbitrap CMBT negative 3ppm | https://metaspace2020.eu/annotations?db_id=304&ds=2021-07-10_00h16m39s |
| 3C EMBL AP-MALDI-Orbitrap CMBT positive 3ppm | https://metaspace2020.eu/annotations?db_id=304&ds=2021-07-10_00h13m11s |
| 3D EMBL AP-MALDI-Orbitrap ClCCA negative 3ppm | https://metaspace2020.eu/annotations?db_id=304&ds=2021-06-18_10h45m03s |

| | |
|---|---|
| 3D EMBL AP-MALDI-Orbitrap ClCCA positive 3ppm | https://metaspace2020.eu/annotations?db_id=304&ds=2021-06-21_12h34m54s |
| 3F EMBL AP-MALDI-Orbitrap DHAP negative 3ppm | https://metaspace2020.eu/annotations?db_id=304&ds=2021-06-18_10h52m02s |
| 3F EMBL AP-MALDI-Orbitrap DHAP positive 3ppm | https://metaspace2020.eu/annotations?db_id=304&ds=2021-06-21_12h41m08s |
| 3H EMBL AP-MALDI-Orbitrap NEDC negative 3ppm | https://metaspace2020.eu/annotations?db_id=304&ds=2021-06-18_10h59m48s |
| 3H EMBL AP-MALDI-Orbitrap NEDC positive 3ppm | https://metaspace2020.eu/annotations?db_id=304&ds=2021-06-21_12h50m44s |
| 3J EMBL AP-MALDI-Orbitrap MAPS negative 3ppm | https://metaspace2020.eu/annotations?db_id=304&ds=2021-06-18_10h58m25s |
| 3J EMBL AP-MALDI-Orbitrap MAPS positive 3ppm | https://metaspace2020.eu/annotations?db_id=304&ds=2021-06-21_12h48m21s |
| 6F EMBL AP-MALDI-Orbitrap NOR negative 3ppm | https://metaspace2020.eu/annotations?db_id=304&ds=2021-06-18_11h04m19s |
| 6F EMBL AP-MALDI-Orbitrap NOR positive 3ppm | https://metaspace2020.eu/annotations?db_id=304&ds=2021-06-21_12h54m04s |
| 6G EMBL AP-MALDI-Orbitrap CHCA negative 3ppm | https://metaspace2020.eu/annotations?db_id=304&ds=2021-06-18_10h41m59s |
| 6G EMBL AP-MALDI-Orbitrap CHCA positive 3ppm | https://metaspace2020.eu/annotations?db_id=304&ds=2021-06-21_12h32m53s |
| 6H EMBL AP-MALDI-Orbitrap PNDI negative 3ppm | https://metaspace2020.eu/annotations?db_id=304&ds=2021-06-18_11h09m13s |
| 6H EMBL AP-MALDI-Orbitrap PNDI positive 3ppm | https://metaspace2020.eu/annotations?db_id=304&ds=2021-06-21_12h59m59s |
| 6J EMBL AP-MALDI-Orbitrap 9AA negative 3ppm | https://metaspace2020.eu/annotations?db_id=304&ds=2021-06-18_10h37m54s |
| 6J EMBL AP-MALDI-Orbitrap 9AA positive 3ppm | https://metaspace2020.eu/annotations?db_id=304&ds=2021-06-21_12h29m16s |
| 7A EMBL AP-MALDI-Orbitrap DAN negative 3ppm | https://metaspace2020.eu/annotations?db_id=304&ds=2021-06-18_10h49m47s |
| 7A EMBL AP-MALDI-Orbitrap DAN positive 3ppm | https://metaspace2020.eu/annotations?db_id=304&ds=2021-06-21_12h38m55s |
| 7J EMBL AP-MALDI-Orbitrap DHB negative 3ppm | https://metaspace2020.eu/annotations?db_id=304&ds=2021-06-18_10h54m38s |
| 7J EMBL AP-MALDI-Orbitrap DHB positive 3ppm | https://metaspace2020.eu/annotations?db_id=304&ds=2021-06-21_12h45m04s |
| 10C Dreisewerd group MALDI2-qTOF DHAP negative | https://metaspace2020.eu/annotations?db_id=304&ds=2021-07-29_18h23m02s |
| 10D Dreisewerd group MALDI2-qTOF DHAP positive | https://metaspace2020.eu/annotations?db_id=304&ds=2021-07-30_02h04m13s |
| 10G PNNL MALDI-FTICR NEDC negative low | https://metaspace2020.eu/annotations?db_id=304&ds=2022-03-11_09h27m55s |
| 10G PNNL MALDI-FTICR NEDC negative high | https://metaspace2020.eu/annotations?db_id=304&ds=2022-03-11_09h27m57s |
| 10I PNNL MALDI-FTICR DHB positive low | https://metaspace2020.eu/annotations?db_id=304&ds=2022-03-11_09h27m51s |
| 10I PNNL MALDI-FTICR DHB positive high | https://metaspace2020.eu/annotations?db_id=304&ds=2022-03-11_09h27m53s |

| | |
|---|---|
| 12C HS Mannheim MALDI-FTICR DHB positive | https://metaspace2020.eu/annotations?db_id=304&ds=2021-07-30_05h15m49s |
| 12E HS Mannheim MALDI-FTICR DAN negative | https://metaspace2020.eu/annotations?db_id=304&ds=2021-07-30_05h55m13s |
| 12G HS Mannheim MALDI-qTOF DHB positive | https://metaspace2020.eu/annotations?db_id=304&ds=2021-07-30_06h15m40s |
| 12I HS Mannheim MALDI-qTOF DAN negative | https://metaspace2020.eu/annotations?db_id=304&ds=2021-07-30_07h11m07s |
| 13G Bruker MALDI-qTOF DHB positive | https://metaspace2020.eu/annotations?db_id=304&ds=2021-07-30_03h21m10s |
| 13J Bruker MALDI-qTOF NEDC negative | https://metaspace2020.eu/annotations?db_id=304&ds=2021-07-30_02h22m33s |
| 14B MPI Bremen AP-MALDI-Orbitrap SDHB positive low | https://metaspace2020.eu/annotations?db_id=304&ds=2022-02-18_23h32m39s |
| 14B MPI Bremen AP-MALDI-Orbitrap SDHB positive high | https://metaspace2020.eu/annotations?db_id=304&ds=2022-02-19_09h24m30s |
| 14D MPI Bremen AP-MALDI-Orbitrap 9AA negative low | https://metaspace2020.eu/annotations?db_id=304&ds=2022-02-18_23h32m51s |
| 14D MPI Bremen AP-MALDI-Orbitrap 9AA negative high | https://metaspace2020.eu/annotations?db_id=304&ds=2022-02-18_23h32m54s |
| 15C EMBL AP-MALDI-Orbitrap pNA negative | https://metaspace2020.eu/annotations?db_id=304&ds=2022-06-01_18h21m54s |
| 15C EMBL AP-MALDI-Orbitrap pNA positive | https://metaspace2020.eu/annotations?db_id=304&ds=2022-06-01_18h21m59s |
| 1A AstraZeneca DESI-Orbitrap none positive | https://metaspace2020.eu/annotations?db_id=304&ds=2021-12-23_09h43m49s |
| 1C AstraZeneca DESI-Orbitrap none negative | https://metaspace2020.eu/annotations?db_id=304&ds=2021-12-19_16h27m49s |
| 1H UT Austin DESI-Orbitrap none negative | https://metaspace2020.eu/annotations?db_id=304&ds=2021-12-19_10h34m32s |
| 1J UT Austin DESI-Orbitrap none positive | https://metaspace2020.eu/annotations?db_id=304&ds=2021-12-19_23h56m42s |
| 2F Dreisewerd group MALDI2-qTOF DHAP positive | https://metaspace2020.eu/annotations?db_id=304&ds=2022-03-14_22h35m49s |
| 2I Dreisewerd group MALDI-qTOF DHAP positive | https://metaspace2020.eu/annotations?db_id=304&ds=2022-03-14_22h35m45s |
| 3A Dreisewerd group MALDI2-Orbitrap DHAP negative | https://metaspace2020.eu/annotations?db_id=304&ds=2021-12-02_17h01m03s |
| 3C EMBL AP-MALDI-Orbitrap CMBT negative | https://metaspace2020.eu/annotations?db_id=304&ds=2022-06-01_18h22m01s |
| 3C EMBL AP-MALDI-Orbitrap CMBT positive | https://metaspace2020.eu/annotations?db_id=304&ds=2022-06-01_18h22m03s |
| 3D EMBL AP-MALDI-Orbitrap ClCCA negative | https://metaspace2020.eu/annotations?db_id=304&ds=2021-11-08_17h35m44s |
| 3D EMBL AP-MALDI-Orbitrap ClCCA positive | https://metaspace2020.eu/annotations?db_id=304&ds=2021-11-08_17h35m46s |
| 3F EMBL AP-MALDI-Orbitrap DHAP negative | https://metaspace2020.eu/annotations?db_id=304&ds=2021-11-08_17h35m48s |
| 3F EMBL AP-MALDI-Orbitrap DHAP positive | https://metaspace2020.eu/annotations?db_id=304&ds=2021-11-08_17h35m50s |
| 3H EMBL AP-MALDI-Orbitrap NEDC | https://metaspace2020.eu/annotations?db_id=304&ds=2021-11-08_17h35m34s |

| | |
|---|---|
| negative | |
| 3H EMBL AP-MALDI-Orbitrap NEDC positive | https://metaspace2020.eu/annotations?db_id=304&ds=2021-11-08_17h35m32s |
| 3J EMBL AP-MALDI-Orbitrap MAPS negative | https://metaspace2020.eu/annotations?db_id=304&ds=2021-11-08_17h35m40s |
| 3J EMBL AP-MALDI-Orbitrap MAPS positive | https://metaspace2020.eu/annotations?db_id=304&ds=2021-11-08_17h35m42s |
| 4F Dreisewerd group MALDI2-Orbitrap DHAP positive | https://metaspace2020.eu/annotations?db_id=304&ds=2021-12-02_16h52m45s |
| 5G U Copenhagen DESI-Orbitrap none negative low | https://metaspace2020.eu/annotations?db_id=304&ds=2022-03-11_12h02m38s |
| 5I U Copenhagen DESI-Orbitrap none negative high | https://metaspace2020.eu/annotations?db_id=304&ds=2022-03-11_12h02m40s |
| 5H U Copenhagen DESI-Orbitrap none positive low | https://metaspace2020.eu/annotations?db_id=304&ds=2022-03-11_12h02m34s |
| 5J U Copenhagen DESI-Orbitrap none positive high | https://metaspace2020.eu/annotations?db_id=304&ds=2022-03-11_12h02m36s |
| 6F EMBL AP-MALDI-Orbitrap NOR negative | https://metaspace2020.eu/annotations?db_id=304&ds=2021-11-08_17h35m21s |
| 6F EMBL AP-MALDI-Orbitrap NOR positive | https://metaspace2020.eu/annotations?db_id=304&ds=2021-11-08_17h35m19s |
| 6G EMBL AP-MALDI-Orbitrap CHCA negative | https://metaspace2020.eu/annotations?db_id=304&ds=2021-11-08_17h35m29s |
| 6G EMBL AP-MALDI-Orbitrap CHCA positive | https://metaspace2020.eu/annotations?db_id=304&ds=2021-11-08_17h35m27s |
| 6H EMBL AP-MALDI-Orbitrap PNDI negative | https://metaspace2020.eu/annotations?db_id=304&ds=2021-11-08_17h35m38s |
| 6H EMBL AP-MALDI-Orbitrap PNDI positive | https://metaspace2020.eu/annotations?db_id=304&ds=2021-11-08_17h35m36s |
| 6J EMBL AP-MALDI-Orbitrap 9AA negative | https://metaspace2020.eu/annotations?db_id=304&ds=2021-11-08_17h35m25s |
| 6J EMBL AP-MALDI-Orbitrap 9AA positive | https://metaspace2020.eu/annotations?db_id=304&ds=2021-11-08_17h35m23s |
| 7A EMBL AP-MALDI-Orbitrap DAN negative | https://metaspace2020.eu/annotations?db_id=304&ds=2021-11-08_17h35m16s |
| 7A EMBL AP-MALDI-Orbitrap DAN positive | https://metaspace2020.eu/annotations?db_id=304&ds=2021-11-08_17h35m14s |
| 7B Dreisewerd group MALDI-qTOF DHAP negative | https://metaspace2020.eu/annotations?db_id=304&ds=2022-03-14_22h35m47s |
| 7D Dreisewerd group MALDI2-qTOF DHAP negative | https://metaspace2020.eu/annotations?db_id=304&ds=2022-03-14_22h35m51s |
| 7J EMBL AP-MALDI-Orbitrap DHB negative | https://metaspace2020.eu/annotations?db_id=304&ds=2021-11-08_17h35m13s |
| 7J EMBL AP-MALDI-Orbitrap DHB positive | https://metaspace2020.eu/annotations?db_id=304&ds=2021-11-08_17h27m55s |
| 8A JLU Giessen AP-MALDI-Orbitrap DHB positive | https://metaspace2020.eu/annotations?db_id=304&ds=2021-07-30_04h08m37s |
| 8D JLU Giessen AP-MALDI-Orbitrap 9AA negative | https://metaspace2020.eu/annotations?db_id=304&ds=2021-07-30_04h35m08s |
| 9A NCSU IR-MALDESI-Orbitrap water positive | https://metaspace2020.eu/annotations?db_id=304&ds=2021-07-30_10h05m21s |

| | |
|---|---|
| 9E NCSU IR-MALDESI-Orbitrap water negative | https://metaspace2020.eu/annotations?db_id=304&ds=2021-07-30_09h30m43s |

#### Supplementary Table 9

Classification of reference metabolites by biochemical function

| Name | Group of pathways | Biochemical pathway |
|---|---|---|
| Agmatine | Amino acid metabolism | Arg metabolism/Urea cycle |
| Arginine | Amino acid metabolism | Arg metabolism/Urea cycle |
| Argininosuccinic acid | Amino acid metabolism | Arg metabolism/Urea cycle |
| Carbamoyl phosphate | Amino acid metabolism | Arg metabolism/Urea cycle |
| Citrulline | Amino acid metabolism | Arg metabolism/Urea cycle |
| Ornithine | Amino acid metabolism | Arg metabolism/Urea cycle |
| Asparagine | Amino acid metabolism | Asp and Asn metabolism |
| Aspartic acid | Amino acid metabolism | Asp and Asn metabolism |
| N-Acetylaspartic acid | Amino acid metabolism | Asp and Asn metabolism |
| Carnitine | Amino acid metabolism | Carnitine biosynthesis |
| N6,N6,N6-Trimethyllysine | Amino acid metabolism | Carnitine biosynthesis |
| Creatine | Amino acid metabolism | Creatinine biosynthesis |
| Creatinine | Amino acid metabolism | Creatinine biosynthesis |
| Phosphocreatine | Amino acid metabolism | Creatinine biosynthesis |
| gamma-Aminobutyric acid | Amino acid metabolism | Glu and Gln metabolism |
| Glutamic acid | Amino acid metabolism | Glu and Gln metabolism |
| Glutamine | Amino acid metabolism | Glu and Gln metabolism |
| N-Acetylaspartylglutamic acid | Amino acid metabolism | Glu and Gln metabolism |
| Carnosine | Amino acid metabolism | His metabolism |
| Histamine | Amino acid metabolism | His metabolism |
| Histidine | Amino acid metabolism | His metabolism |
| Urocanic acid | Amino acid metabolism | His metabolism |
| Cysteine | Amino acid metabolism | Metabolism of sulfur-containing amino acids |
| Cystine | Amino acid metabolism | Metabolism of sulfur-containing amino acids |

| | | |
|---|---|---|
| Glutathione | Amino acid metabolism | Metabolism of sulfur-containing amino acids |
| Oxidized glutathione | Amino acid metabolism | Metabolism of sulfur-containing amino acids |
| Taurine | Amino acid metabolism | Metabolism of sulfur-containing amino acids |
| Homocysteine | Amino acid metabolism | Methionine cycle |
| Methionine | Amino acid metabolism | Methionine cycle |
| S-Adenosylhomocysteine | Amino acid metabolism | Methionine cycle |
| S-Adenosylmethionine | Amino acid metabolism | Methionine cycle |
| Glycine | Amino acid metabolism | Other amino acid metabolism |
| Valine | Amino acid metabolism | Other amino acid metabolism |
| Acetoacetic acid | Amino acid metabolism | Phe and Tyr metabolism |
| Dopamine | Amino acid metabolism | Phe and Tyr metabolism |
| Epinephrine | Amino acid metabolism | Phe and Tyr metabolism |
| L-Dopa | Amino acid metabolism | Phe and Tyr metabolism |
| Phenylalanine | Amino acid metabolism | Phe and Tyr metabolism |
| Tyrosine | Amino acid metabolism | Phe and Tyr metabolism |
| Putrescine | Amino acid metabolism | Polyamine biosynthesis |
| Spermidine | Amino acid metabolism | Polyamine biosynthesis |
| Spermine | Amino acid metabolism | Polyamine biosynthesis |
| 4-Hydroxyproline | Amino acid metabolism | Pro metabolism |
| Proline | Amino acid metabolism | Pro metabolism |
| Phosphoserine | Amino acid metabolism | Ser metabolism |
| Serine | Amino acid metabolism | Ser metabolism |
| 3-Hydroxyanthranilic acid | Amino acid metabolism | Trp metabolism |
| 5-Hydroxyindoleacetic acid | Amino acid metabolism | Trp metabolism |
| Indole | Amino acid metabolism | Trp metabolism |
| Kynurenic acid | Amino acid metabolism | Trp metabolism |
| Kynurenine | Amino acid metabolism | Trp metabolism |
| Melatonin | Amino acid metabolism | Trp metabolism |
| Quinolinic acid | Amino acid metabolism | Trp metabolism |

| | | |
|---|---|---|
| Serotonin | Amino acid metabolism | Trp metabolism |
| Tryptophan | Amino acid metabolism | Trp metabolism |
| 2-Oxoglutaric acid | Carbohydrate metabolism | Citric acid cycle |
| Acetyl-CoA | Carbohydrate metabolism | Citric acid cycle |
| cis-Aconitic acid | Carbohydrate metabolism | Citric acid cycle |
| Citric acid | Carbohydrate metabolism | Citric acid cycle |
| Fumaric acid | Carbohydrate metabolism | Citric acid cycle |
| Malic acid | Carbohydrate metabolism | Citric acid cycle |
| Oxalacetic acid | Carbohydrate metabolism | Citric acid cycle |
| Succinic acid | Carbohydrate metabolism | Citric acid cycle |
| 3-Phosphoglyceric acid | Carbohydrate metabolism | Glycolysis |
| Dihydroxyacetone | Carbohydrate metabolism | Glycolysis |
| Dihydroxyacetone phosphate | Carbohydrate metabolism | Glycolysis |
| Fructose 1,6-bisphosphate | Carbohydrate metabolism | Glycolysis |
| Glucose | Carbohydrate metabolism | Glycolysis |
| Glucose 6-phosphate | Carbohydrate metabolism | Glycolysis |
| Glycerol 3-phosphate | Carbohydrate metabolism | Glycolysis |
| Lactic acid | Carbohydrate metabolism | Glycolysis |
| Phosphoenolpyruvic acid | Carbohydrate metabolism | Glycolysis |
| Pyruvic acid | Carbohydrate metabolism | Glycolysis |
| Glucosamine | Carbohydrate metabolism | Hexosamine biosynthetic pathway |
| Glucosamine 6-phosphate | Carbohydrate metabolism | Hexosamine biosynthetic pathway |
| N-Acetylgalactosamine 1-phosphate | Carbohydrate metabolism | Hexosamine biosynthetic pathway |
| N-Acetylglucosamine | Carbohydrate metabolism | Hexosamine biosynthetic pathway |
| Uridine diphosphate glucose | Carbohydrate metabolism | Hexosamine biosynthetic pathway |
| Uridine diphosphate glucuronic acid | Carbohydrate metabolism | Hexosamine biosynthetic pathway |
| Uridine diphosphate-N-acetylglucosami | Carbohydrate metabolism | Hexosamine biosynthetic pathway |

| | | |
|---|---|---|
| ne | | |
| myo-Inositol | Carbohydrate metabolism | Inositol metabolism |
| myo-Inositol 1-phosphate | Carbohydrate metabolism | Inositol metabolism |
| 6-Phosphogluconic acid | Carbohydrate metabolism | Pentose phosphate pathway |
| Gluconic acid | Carbohydrate metabolism | Pentose phosphate pathway |
| Glucose 6-phosphate | Carbohydrate metabolism | Pentose phosphate pathway |
| Ribose 5-phosphate | Carbohydrate metabolism | Pentose phosphate pathway |
| Ribulose | Carbohydrate metabolism | Pentose phosphate pathway |
| Ribulose 5-phosphate | Carbohydrate metabolism | Pentose phosphate pathway |
| Arachidonic acid | Lipid metabolism | Arachidonic acid metabolism |
| Prostaglandin E1 | Lipid metabolism | Arachidonic acid metabolism |
| Cholesteryl acetate | Lipid metabolism | Cholesterol metabolism |
| Cholesteryl ester 17:0 | Lipid metabolism | Cholesterol metabolism |
| Cholic acid | Lipid metabolism | Cholesterol metabolism |
| Estradiol | Lipid metabolism | Cholesterol metabolism |
| Glucosyl cholesterol | Lipid metabolism | Cholesterol metabolism |
| Taurocholic acid | Lipid metabolism | Cholesterol metabolism |
| 3-Hydroxymethylglutaric acid | Lipid metabolism | Cholesterol synthesis |
| Acetyl-CoA | Lipid metabolism | Cholesterol synthesis |
| Cholesterol | Lipid metabolism | Cholesterol synthesis |
| Mevalonic acid | Lipid metabolism | Cholesterol synthesis |
| Acetylcholine | Lipid metabolism | Choline metabolism |
| Choline | Lipid metabolism | Choline metabolism |
| Glycerophosphocholine | Lipid metabolism | Choline metabolism |
| Acetylcarnitine | Lipid metabolism | Fatty acid oxidation |
| CAR 16:0 | Lipid metabolism | Fatty acid oxidation |
| Carnitine | Lipid metabolism | Fatty acid oxidation |
| Palmitoyl-CoA | Lipid metabolism | Fatty acid oxidation |
| Propionylcarnitine | Lipid metabolism | Fatty acid oxidation |
| Acetyl-CoA | Lipid metabolism | Fatty acid synthesis |

| | | |
|---|---|---|
| Butyric acid | Lipid metabolism | Fatty acid synthesis |
| Citric acid | Lipid metabolism | Fatty acid synthesis |
| Malonic acid | Lipid metabolism | Fatty acid synthesis |
| Palmitic acid | Lipid metabolism | Fatty acid synthesis |
| DG 18:0-22:6 | Lipid metabolism | Glycerolipid metabolism |
| MG 18:1 | Lipid metabolism | Glycerolipid metabolism |
| TG 15:0-18:1-15:0 | Lipid metabolism | Glycerolipid metabolism |
| Cardiolipin 18:1 | Lipid metabolism | Phospholipid metabolism |
| Lyso PA 18:1 | Lipid metabolism | Phospholipid metabolism |
| Lyso PC 15:0 | Lipid metabolism | Phospholipid metabolism |
| Lyso PE 18:0 | Lipid metabolism | Phospholipid metabolism |
| Lyso PG 16:0 | Lipid metabolism | Phospholipid metabolism |
| Lyso PI 17:1 | Lipid metabolism | Phospholipid metabolism |
| Lyso PS 17:1 | Lipid metabolism | Phospholipid metabolism |
| PA 16:0-18:1 | Lipid metabolism | Phospholipid metabolism |
| PC (POPC) 16:0-18:1 | Lipid metabolism | Phospholipid metabolism |
| PC 15:0-18:1 | Lipid metabolism | Phospholipid metabolism |
| PE 18:0-20:4 | Lipid metabolism | Phospholipid metabolism |
| PG 16:0-18:1 | Lipid metabolism | Phospholipid metabolism |
| PI 16:0-18:1 | Lipid metabolism | Phospholipid metabolism |
| PI(5)P 18:1 | Lipid metabolism | Phospholipid metabolism |
| PS (POPS) 16:0-18:1 | Lipid metabolism | Phospholipid metabolism |
| Cer d18:1-22:0 | Lipid metabolism | Sphingolipid metabolism |
| GlcCer d18:1-18:0 | Lipid metabolism | Sphingolipid metabolism |
| GM3 Ganglioside 18:0 | Lipid metabolism | Sphingolipid metabolism |
| SM d18:1-16:0 | Lipid metabolism | Sphingolipid metabolism |
| Sulfo GalCer 18:0(2S-OH) | Lipid metabolism | Sphingolipid metabolism |
| Coenzyme A | Metabolism of vitamins and cofactors | Coenzyme A biosynthesis |
| Pantetheine | Metabolism of vitamins and cofactors | Coenzyme A biosynthesis |

| | | |
|---|---|---|
| Pantothenic acid | Metabolism of vitamins and cofactors | Coenzyme A biosynthesis |
| Folic acid | Metabolism of vitamins and cofactors | Folate cycle |
| Tetrahydrofolic acid | Metabolism of vitamins and cofactors | Folate cycle |
| Niacinamide | Metabolism of vitamins and cofactors | NAD cycle |
| Nicotinamide adenine dinucleotide (NAD) | Metabolism of vitamins and cofactors | NAD cycle |
| Nicotinamide adenine dinucleotide, reduced (NADH) | Metabolism of vitamins and cofactors | NAD cycle |
| Nicotinamide riboside | Metabolism of vitamins and cofactors | NAD cycle |
| Nicotinamide ribotide | Metabolism of vitamins and cofactors | NAD cycle |
| Nicotinic acid | Metabolism of vitamins and cofactors | NAD cycle |
| Flavin adenine dinucleotide | Metabolism of vitamins and cofactors | Redox metabolism |
| Flavin mononucleotide | Metabolism of vitamins and cofactors | Redox metabolism |
| Nicotinamide adenine dinucleotide (NAD) | Metabolism of vitamins and cofactors | Redox metabolism |
| Nicotinamide adenine dinucleotide, reduced (NADH) | Metabolism of vitamins and cofactors | Redox metabolism |
| alpha-Tocopherol | Metabolism of vitamins and cofactors | Vitamin metabolism |
| Ascorbic acid | Metabolism of vitamins and cofactors | Vitamin metabolism |
| Biotin | Metabolism of vitamins and cofactors | Vitamin metabolism |
| Cholecalciferol | Metabolism of vitamins and cofactors | Vitamin metabolism |
| Phylloquinone | Metabolism of vitamins and cofactors | Vitamin metabolism |
| Pyridoxine | Metabolism of vitamins and cofactors | Vitamin metabolism |
| Retinoic acid | Metabolism of vitamins and cofactors | Vitamin metabolism |

| | | |
|---|---|---|
| Riboflavin | Metabolism of vitamins and cofactors | Vitamin metabolism |
| Thiamine | Metabolism of vitamins and cofactors | Vitamin metabolism |
| Adenine | Nucleotide metabolism | Purine metabolism |
| Adenosine diphosphate | Nucleotide metabolism | Purine metabolism |
| Adenosine monophosphate | Nucleotide metabolism | Purine metabolism |
| Adenosine triphosphate | Nucleotide metabolism | Purine metabolism |
| Cyclic adenosine monophosphate | Nucleotide metabolism | Purine metabolism |
| Guanine | Nucleotide metabolism | Purine metabolism |
| Hypoxanthine | Nucleotide metabolism | Purine metabolism |
| Xanthine | Nucleotide metabolism | Purine metabolism |
| 4,5-Dihydroorotic acid | Nucleotide metabolism | Pyrimidine metabolism |
| Cytidine | Nucleotide metabolism | Pyrimidine metabolism |
| Cytidine monophosphate | Nucleotide metabolism | Pyrimidine metabolism |
| Cytosine | Nucleotide metabolism | Pyrimidine metabolism |
| Orotic acid | Nucleotide metabolism | Pyrimidine metabolism |
| Thymine | Nucleotide metabolism | Pyrimidine metabolism |
| Uracil | Nucleotide metabolism | Pyrimidine metabolism |
| Ureidosuccinic acid | Nucleotide metabolism | Pyrimidine metabolism |
| Uric acid | Nucleotide metabolism | Pyrimidine metabolism |
| Uridine | Nucleotide metabolism | Pyrimidine metabolism |
| Uridine diphosphate | Nucleotide metabolism | Pyrimidine metabolism |

#### Supplementary Table 10

Classification of reference metabolites by chemical class and subclass

| **Name** | **Chemical class** | **Chemical subclass** |
|---|---|---|
| Acetylcarnitine | Amines | Quarternary ammonium amines |
| Acetylcholine | Amines | Quarternary ammonium amines |

| | | |
|---|---|---|
| CAR 16:0 | Amines | Quarternary ammonium amines |
| Carnitine | Amines | Quarternary ammonium amines |
| Choline | Amines | Quarternary ammonium amines |
| Glycerophosphocholine | Amines | Quarternary ammonium amines |
| N6,N6,N6-Trimethyllysine | Amines | Quarternary ammonium amines |
| Propionylcarnitine | Amines | Quarternary ammonium amines |
| Agmatine | Amines | Other amines |
| Carbamoyl phosphate | Amines | Other amines |
| Histamine | Amines | Other amines |
| Putrescine | Amines | Other amines |
| Spermidine | Amines | Other amines |
| Spermine | Amines | Other amines |
| 3-Hydroxyanthranilic acid | Amino acids, peptides, and analogues | Acidic amino acids |
| Argininosuccinic acid | Amino acids, peptides, and analogues | Acidic amino acids |
| Aspartic acid | Amino acids, peptides, and analogues | Acidic amino acids |
| Glutamic acid | Amino acids, peptides, and analogues | Acidic amino acids |
| N-Acetylaspartic acid | Amino acids, peptides, and analogues | Acidic amino acids |
| N-Acetylaspartylglutamic acid | Amino acids, peptides, and analogues | Acidic amino acids |
| Phosphoserine | Amino acids, peptides, and analogues | Acidic amino acids |
| Ureidosuccinic acid | Amino acids, peptides, and analogues | Acidic amino acids |
| Agmatine | Amino acids, peptides, and analogues | Arginine derivatives (guanidines) |
| Arginine | Amino acids, peptides, and analogues | Arginine derivatives (guanidines) |

| | | |
|---|---|---|
| Creatine | Amino acids, peptides, and analogues | Arginine derivatives (guanidines) |
| Creatinine | Amino acids, peptides, and analogues | Arginine derivatives (guanidines) |
| Phosphocreatine | Amino acids, peptides, and analogues | Arginine derivatives (guanidines) |
| Dopamine | Amino acids, peptides, and analogues | Aromatic amino acids |
| Epinephrine | Amino acids, peptides, and analogues | Aromatic amino acids |
| Kynurenine | Amino acids, peptides, and analogues | Aromatic amino acids |
| L-Dopa | Amino acids, peptides, and analogues | Aromatic amino acids |
| Phenylalanine | Amino acids, peptides, and analogues | Aromatic amino acids |
| Tyrosine | Amino acids, peptides, and analogues | Aromatic amino acids |
| Carnosine | Amino acids, peptides, and analogues | Histidine derivatives (imidazoles) |
| Histamine | Amino acids, peptides, and analogues | Histidine derivatives (imidazoles) |
| Histidine | Amino acids, peptides, and analogues | Histidine derivatives (imidazoles) |
| Urocanic acid | Amino acids, peptides, and analogues | Histidine derivatives (imidazoles) |
| Glycine | Amino acids, peptides, and analogues | Nonpolar amino acids |
| Proline | Amino acids, peptides, and analogues | Nonpolar amino acids |
| Valine | Amino acids, peptides, and analogues | Nonpolar amino acids |
| 4-Hydroxyproline | Amino acids, peptides, and analogues | Polar amino acids |
| Asparagine | Amino acids, peptides, and analogues | Polar amino acids |
| Carnitine | Amino acids, peptides, and analogues | Polar amino acids |
| Citrulline | Amino acids, peptides, and analogues | Polar amino acids |

| | | |
|---|---|---|
| gamma-Aminobutyric acid | Amino acids, peptides, and analogues | Polar amino acids |
| Glutamine | Amino acids, peptides, and analogues | Polar amino acids |
| N6,N6,N6-Trimethyllysine | Amino acids, peptides, and analogues | Polar amino acids |
| Ornithine | Amino acids, peptides, and analogues | Polar amino acids |
| Pantothenic acid | Amino acids, peptides, and analogues | Polar amino acids |
| Serine | Amino acids, peptides, and analogues | Polar amino acids |
| Cysteine | Amino acids, peptides, and analogues | Sulphur-containing amino acids |
| Cystine | Amino acids, peptides, and analogues | Sulphur-containing amino acids |
| Glutathione | Amino acids, peptides, and analogues | Sulphur-containing amino acids |
| Homocysteine | Amino acids, peptides, and analogues | Sulphur-containing amino acids |
| Methionine | Amino acids, peptides, and analogues | Sulphur-containing amino acids |
| Oxidized glutathione | Amino acids, peptides, and analogues | Sulphur-containing amino acids |
| Pantetheine | Amino acids, peptides, and analogues | Sulphur-containing amino acids |
| S-Adenosylhomocysteine | Amino acids, peptides, and analogues | Sulphur-containing amino acids |
| S-Adenosylmethionine | Amino acids, peptides, and analogues | Sulphur-containing amino acids |
| Taurine | Amino acids, peptides, and analogues | Sulphur-containing amino acids |
| 5-Hydroxyindoleacetic acid | Amino acids, peptides, and analogues | Tryptophan derivatives (indoles) |
| Indole | Amino acids, peptides, and analogues | Tryptophan derivatives (indoles) |
| Melatonin | Amino acids, peptides, and analogues | Tryptophan derivatives (indoles) |
| Serotonin | Amino acids, peptides, and analogues | Tryptophan derivatives (indoles) |

| | | |
|---|---|---|
| Tryptophan | Amino acids, peptides, and analogues | Tryptophan derivatives (indoles) |
| Glucosamine | Carbohydrates | Carbohydrate amines |
| N-Acetylglucosamine | Carbohydrates | Carbohydrate amines |
| Glucosamine 6-phosphate | Carbohydrates | Carbohydrate amino-phosphates |
| N-Acetylgalactosamine 1-phosphate | Carbohydrates | Carbohydrate amino-phosphates |
| 6-Phosphogluconic acid | Carbohydrates | Carbohydrate phosphates |
| Dihydroxyacetone phosphate | Carbohydrates | Carbohydrate phosphates |
| Fructose 1,6-bisphosphate | Carbohydrates | Carbohydrate phosphates |
| Glucose 6-phosphate | Carbohydrates | Carbohydrate phosphates |
| Glycerol 3-phosphate | Carbohydrates | Carbohydrate phosphates |
| myo-Inositol 1-phosphate | Carbohydrates | Carbohydrate phosphates |
| Ribose 5-phosphate | Carbohydrates | Carbohydrate phosphates |
| Ribulose 5-phosphate | Carbohydrates | Carbohydrate phosphates |
| Dihydroxyacetone | Carbohydrates | Carbohydrates |
| Gluconic acid | Carbohydrates | Carbohydrates |
| Glucose | Carbohydrates | Carbohydrates |
| myo-Inositol | Carbohydrates | Carbohydrates |
| Ribulose | Carbohydrates | Carbohydrates |
| Kynurenic acid | Carboxylic acids | Aromatic acids |
| Nicotinic acid | Carboxylic acids | Aromatic acids |
| Quinolinic acid | Carboxylic acids | Aromatic acids |
| 3-Phosphoglyceric acid | Carboxylic acids | Carboxylic acid phosphate |
| 6-Phosphogluconic acid | Carboxylic acids | Carboxylic acid phosphate |
| Phosphoenolpyruvic acid | Carboxylic acids | Carboxylic acid phosphate |
| Butyric acid | Carboxylic acids | Carboxylic acids |
| cis-Aconitic acid | Carboxylic acids | Carboxylic acids |
| Fumaric acid | Carboxylic acids | Carboxylic acids |
| Malonic acid | Carboxylic acids | Carboxylic acids |
| Succinic acid | Carboxylic acids | Carboxylic acids |

| | | |
|---|---|---|
| 3-Hydroxymethylglutaric acid | Carboxylic acids | Hydroxy acids |
| Ascorbic acid | Carboxylic acids | Hydroxy acids |
| Citric acid | Carboxylic acids | Hydroxy acids |
| Gluconic acid | Carboxylic acids | Hydroxy acids |
| Lactic acid | Carboxylic acids | Hydroxy acids |
| Malic acid | Carboxylic acids | Hydroxy acids |
| Mevalonic acid | Carboxylic acids | Hydroxy acids |
| 2-Oxoglutaric acid | Carboxylic acids | Keto acid |
| Acetoacetic acid | Carboxylic acids | Keto acid |
| Oxalacetic acid | Carboxylic acids | Keto acid |
| Pyruvic acid | Carboxylic acids | Keto acid |
| Arachidonic acid | Lipids and lipid-like molecules | Fatty acyl |
| CAR 16:0 | Lipids and lipid-like molecules | Fatty acyl |
| Palmitic acid | Lipids and lipid-like molecules | Fatty acyl |
| Palmitoyl-CoA | Lipids and lipid-like molecules | Fatty acyl |
| Prostaglandin E1 | Lipids and lipid-like molecules | Fatty acyl |
| DG 18:0-22:6 | Lipids and lipid-like molecules | Glycerolipids |
| MG 18:1 | Lipids and lipid-like molecules | Glycerolipids |
| TG 15:0-18:1-15:0 | Lipids and lipid-like molecules | Glycerolipids |
| Cardiolipin 18:1 | Lipids and lipid-like molecules | Glycerophospholipids |
| Lyso PA 18:1 | Lipids and lipid-like molecules | Glycerophospholipids |
| Lyso PC 15:0 | Lipids and lipid-like molecules | Glycerophospholipids |
| Lyso PE 18:0 | Lipids and lipid-like molecules | Glycerophospholipids |
| Lyso PG 16:0 | Lipids and lipid-like molecules | Glycerophospholipids |
| Lyso PI 17:1 | Lipids and lipid-like molecules | Glycerophospholipids |
| Lyso PS 17:1 | Lipids and lipid-like molecules | Glycerophospholipids |
| PA 16:0-18:1 | Lipids and lipid-like molecules | Glycerophospholipids |
| PC (POPC) 16:0-18:1 | Lipids and lipid-like molecules | Glycerophospholipids |
| PC 15:0-18:1 | Lipids and lipid-like molecules | Glycerophospholipids |
| PE 18:0-20:4 | Lipids and lipid-like molecules | Glycerophospholipids |

| | | |
|---|---|---|
| PG 16:0-18:1 | Lipids and lipid-like molecules | Glycerophospholipids |
| PI 16:0-18:1 | Lipids and lipid-like molecules | Glycerophospholipids |
| PI(5)P 18:1 | Lipids and lipid-like molecules | Glycerophospholipids |
| PS (POPS) 16:0-18:1 | Lipids and lipid-like molecules | Glycerophospholipids |
| alpha-Tocopherol | Lipids and lipid-like molecules | Prenol lipids |
| Phylloquinone | Lipids and lipid-like molecules | Prenol lipids |
| Retinoic acid | Lipids and lipid-like molecules | Prenol lipids |
| Cer d18:1-22:0 | Lipids and lipid-like molecules | Sphingolipids |
| GlcCer d18:1-18:0 | Lipids and lipid-like molecules | Sphingolipids |
| GM3 Ganglioside 18:0 | Lipids and lipid-like molecules | Sphingolipids |
| SM d18:1-16:0 | Lipids and lipid-like molecules | Sphingolipids |
| Sulfo GalCer 18:0(2S-OH) | Lipids and lipid-like molecules | Sphingolipids |
| Cholecalciferol | Lipids and lipid-like molecules | Steroids and steroid derivatives |
| Cholesterol | Lipids and lipid-like molecules | Steroids and steroid derivatives |
| Cholesteryl acetate | Lipids and lipid-like molecules | Steroids and steroid derivatives |
| Cholesteryl ester 17:0 | Lipids and lipid-like molecules | Steroids and steroid derivatives |
| Cholic acid | Lipids and lipid-like molecules | Steroids and steroid derivatives |
| Estradiol | Lipids and lipid-like molecules | Steroids and steroid derivatives |
| Glucosyl cholesterol | Lipids and lipid-like molecules | Steroids and steroid derivatives |
| Taurocholic acid | Lipids and lipid-like molecules | Steroids and steroid derivatives |
| Nicotinamide adenine dinucleotide (NAD) | Nucleosides, nucleotides, and analogues | Nicotinamide derivatives |
| Nicotinamide adenine dinucleotide, reduced (NADH) | Nucleosides, nucleotides, and analogues | Nicotinamide derivatives |
| Nicotinamide riboside | Nucleosides, nucleotides, and analogues | Nicotinamide derivatives |
| Nicotinamide ribotide | Nucleosides, nucleotides, and analogues | Nicotinamide derivatives |
| 4,5-Dihydroorotic acid | Nucleosides, nucleotides, and analogues | Nucleobases and analogs |
| Adenine | Nucleosides, nucleotides, and analogues | Nucleobases and analogs |
| Cytosine | Nucleosides, nucleotides, and analogues | Nucleobases and analogs |

| | | |
|---|---|---|
| Guanine | Nucleosides, nucleotides, and analogues | Nucleobases and analogs |
| Hypoxanthine | Nucleosides, nucleotides, and analogues | Nucleobases and analogs |
| Orotic acid | Nucleosides, nucleotides, and analogues | Nucleobases and analogs |
| Thymine | Nucleosides, nucleotides, and analogues | Nucleobases and analogs |
| Uracil | Nucleosides, nucleotides, and analogues | Nucleobases and analogs |
| Uric acid | Nucleosides, nucleotides, and analogues | Nucleobases and analogs |
| Xanthine | Nucleosides, nucleotides, and analogues | Nucleobases and analogs |
| Cytidine | Nucleosides, nucleotides, and analogues | Nucleosides |
| S-Adenosylhomocysteine | Nucleosides, nucleotides, and analogues | Nucleosides |
| S-Adenosylmethionine | Nucleosides, nucleotides, and analogues | Nucleosides |
| Uridine | Nucleosides, nucleotides, and analogues | Nucleosides |
| Acetyl-CoA | Nucleosides, nucleotides, and analogues | Nucleotides |
| Adenosine diphosphate | Nucleosides, nucleotides, and analogues | Nucleotides |
| Adenosine monophosphate | Nucleosides, nucleotides, and analogues | Nucleotides |
| Adenosine triphosphate | Nucleosides, nucleotides, and analogues | Nucleotides |
| Coenzyme A | Nucleosides, nucleotides, and analogues | Nucleotides |
| Cyclic adenosine monophosphate | Nucleosides, nucleotides, and analogues | Nucleotides |
| Cytidine monophosphate | Nucleosides, nucleotides, and analogues | Nucleotides |
| Flavin adenine dinucleotide | Nucleosides, nucleotides, and analogues | Nucleotides |
| Flavin mononucleotide | Nucleosides, nucleotides, and analogues | Nucleotides |

| | | |
|---|---|---|
| Palmitoyl-CoA | Nucleosides, nucleotides, and analogues | Nucleotides |
| Uridine diphosphate | Nucleosides, nucleotides, and analogues | Nucleotides |
| Uridine diphosphate glucose | Nucleosides, nucleotides, and analogues | Nucleotides |
| Uridine diphosphate glucuronic acid | Nucleosides, nucleotides, and analogues | Nucleotides |
| Uridine diphosphate-N-acetylglucosamine | Nucleosides, nucleotides, and analogues | Nucleotides |
| Acetyl-CoA | Vitamins and cofactors | CoA and derivatives |
| Coenzyme A | Vitamins and cofactors | CoA and derivatives |
| Palmitoyl-CoA | Vitamins and cofactors | CoA and derivatives |
| Flavin adenine dinucleotide | Vitamins and cofactors | Flavins |
| Flavin mononucleotide | Vitamins and cofactors | Flavins |
| Riboflavin | Vitamins and cofactors | Flavins |
| Folic acid | Vitamins and cofactors | Folates |
| Tetrahydrofolic acid | Vitamins and cofactors | Folates |
| alpha-Tocopherol | Vitamins and cofactors | Vitamins and cofactors |
| Biotin | Vitamins and cofactors | Vitamins and cofactors |
| Niacinamide | Vitamins and cofactors | Vitamins and cofactors |
| Nicotinic acid | Vitamins and cofactors | Vitamins and cofactors |
| Pantothenic acid | Vitamins and cofactors | Vitamins and cofactors |
| Phylloquinone | Vitamins and cofactors | Vitamins and cofactors |
| Pyridoxine | Vitamins and cofactors | Vitamins and cofactors |
| Retinoic acid | Vitamins and cofactors | Vitamins and cofactors |
| Thiamine | Vitamins and cofactors | Vitamins and cofactors |

#### Supplementary Table 11

Links to example plots generated with interactive web application

| | |
|---|---|
| Which protocols lead to the detection of UDP-GlcNAc with highest intensity on the AP-MALDI-Orbitrap system? | https://metaspace2020.eu/detectability?filter=nL,name&filterValue=None%7CUridine%20diphosphate-N-acetylglucosamine&xAxis= |

| | |
|---|---|
| | name&yAxis=Sample%20name&agg=log10_intensity&metric=2&page=1&pageSize=30&src=EMBL&vis=2 |
| Which ions are detectable when analysing cholesterol using DHB matrix in positive ion mode on AP-MALDI-Orbitrap system? | https://metaspace2020.eu/detectability?filter=name,Polarity,Matrix%20short&filterValue=Cholesterol%7Cpositive%7CDHB&xAxis=nL&yAxis=a&agg=log10_intensity&metric=2&page=1&pageSize=30&src=EMBL&vis=1 |
| Which matrix is a good choice to analyse intermediates of biochemical pathways relevant to macrophage metabolism in negative polarity on AP-MALDI-Orbitrap system? | https://metaspace2020.eu/detectability?filter=nL,fine_path,Polarity&filterValue=None%7CArg%20metabolism%2FUrea%20cycle%23Carnitine%20biosynthesis%23Metabolism%20of%20sulfur-containing%20amino%20acids%23Polyamine%20biosynthesis%23Citric%20acid%20cycle%23Glycolysis%23Hexosamine%20biosynthetic%20pathway%23Cholesterol%20synthesis%23Fatty%20acid%20oxidation%23Fatty%20acid%20synthesis%23NAD%20cycle%23Redox%20metabolism%7Cnegative&xAxis=Matrix%20short&yAxis=fine_path&agg=log10_intensity&metric=2&page=1&pageSize=30&src=EMBL&vis=1 |
| What is the coverage achieved by DHB and DAN in negative ion mode on the AP-MALDI-Orbitrap system? | https://metaspace2020.eu/detectability?filter=nL,Polarity,Matrix%20short&filterValue=None%7Cnegative%7CDHB%23DAN&xAxis=Matrix%20short&yAxis=fine_class&agg=log10_intensity&metric=2&page=1&pageSize=30&src=EMBL&vis=1 |
| What adducts are typically observed for different chemical classes with DESI technologies? | https://metaspace2020.eu/detectability?filter=nL&filterValue=None&xAxis=a&yAxis=main_coarse_class&agg=fraction_detected&metric=2&page=1&pageSize=30&src=INTERLAB&vis=1 |
| Fraction and mean intensity of detected ions per chemical subclass and matrix in positive ionisation mode (as in Figure 3) | https://metaspace2020.eu/detectability?filter=Polarity,nL&filterValue=positive%7CNone&xAxis=Matrix%20short&yAxis=fine_path&agg=log10_intensity&metric=2&page=1&pageSize=15&src=EMBL&vis=1 |
| Fraction of detected metabolites in chemical class in negative ionisation mode for every interlaboratory survey participant (as in Figure 5) | https://metaspace2020.eu/detectability?filter=Polarity,nL&filterValue=negative%7CNone&xAxis=Sample%20name&yAxis=main_coarse_class&agg=fraction_detected&metric=2&page=1&pageSize=30&src=INTERLAB&vis=2 |
| How does the ion image look for UDP-GlcNAc measured with 9AA in | https://metaspace2020.eu/detectability?filter=nL,name,Matrix%20short,Polarity&filterVal |

| | |
|---|---|
| negative ionisation mode on the AP-MALDI-Orbitrap system? (Click on the dot in the plot to be redirected to METASPACE) | ue=None%7CUridine%20diphosphate-N-acetylglucosamine%7C9AA%7Cnegative&xAxis=name&yAxis=Matrix%20short&agg=log10_intensity&page=1&pageSize=30&src=EMBL&vis=1&cmap=-YlGnBu |

### Supplementary Figures

#### Supplementary Figure 1

Technical drawing of the microwell mould. Also provided as .STP file in the digital supplement.

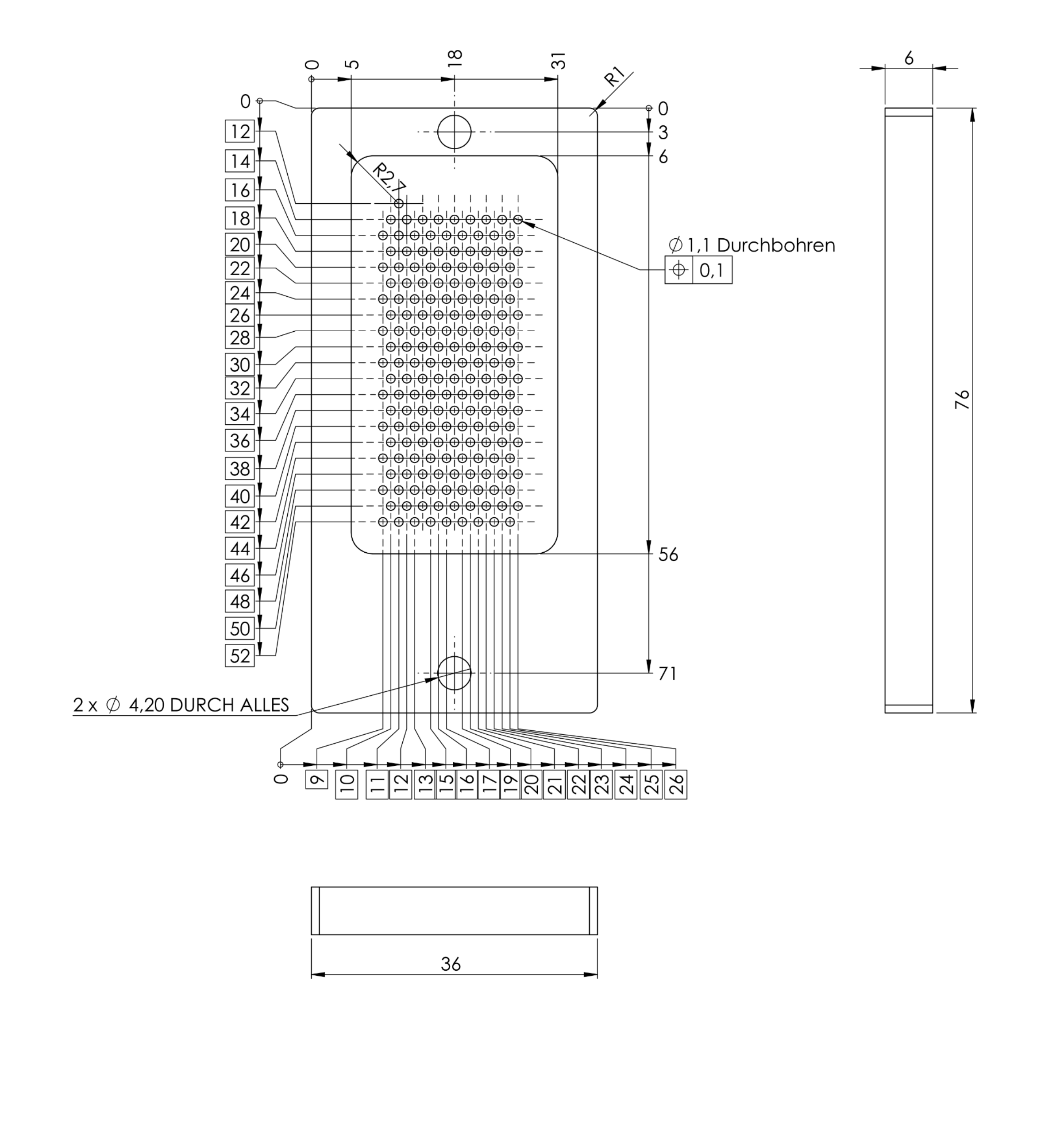

#### Supplementary Figure 2

Plot of four sequential WinCATS programs overlaid in x-y plane, representing the production of a batch of 10 slides. Red: rhodamine fiducial marks, Blue: methanol:chloroform solutions, Yellow: First set of methanol:water solutions, Green: Second set of methanol:water solutions.

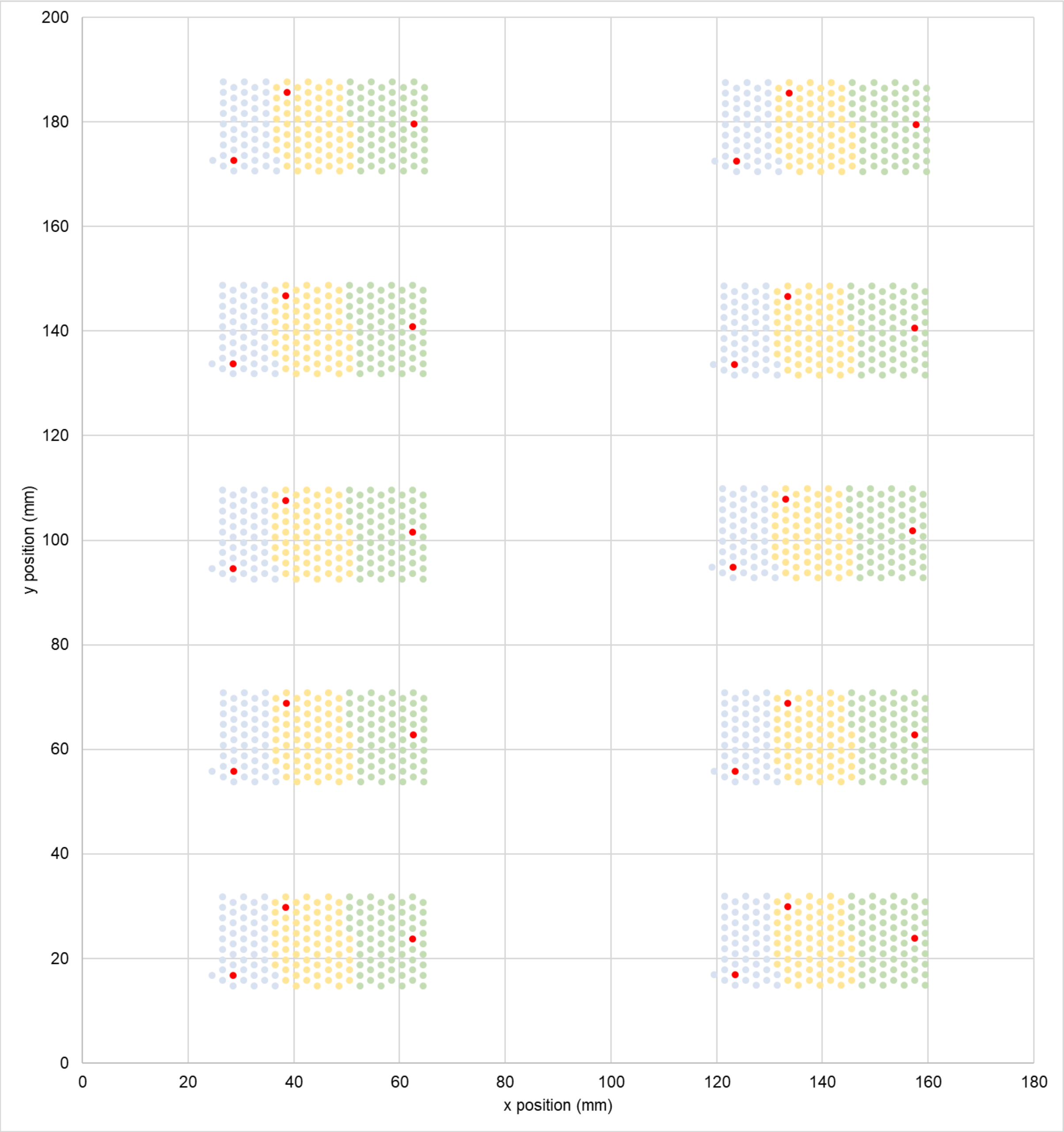

y position (mm)
x position (mm)
0
20
40
60
80
100
120
140
160
180
200

#### Supplementary Figure 3

Standard sample layout with metabolite indices

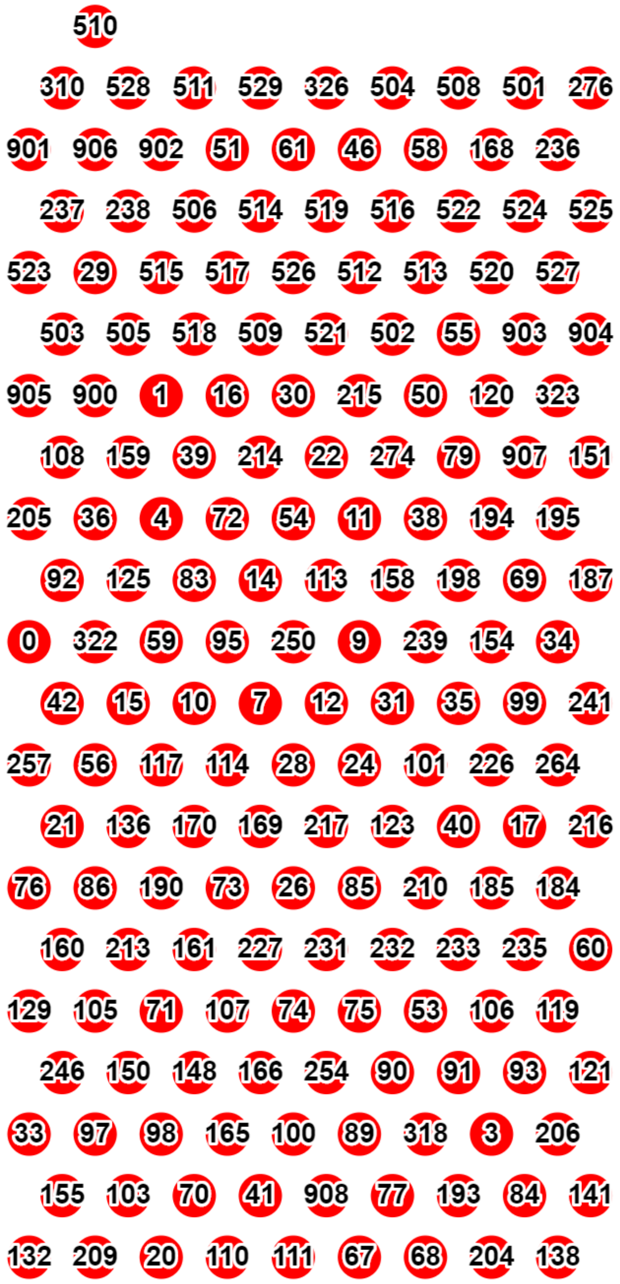

#### Supplementary Figure 4

Map of common biochemical pathways from iPath3 [18] in high resolution and using original colours. The plot shows the distribution of metabolites included in the reference sample (in black) across different pathways, where dots represent molecules and lines represent reactions.

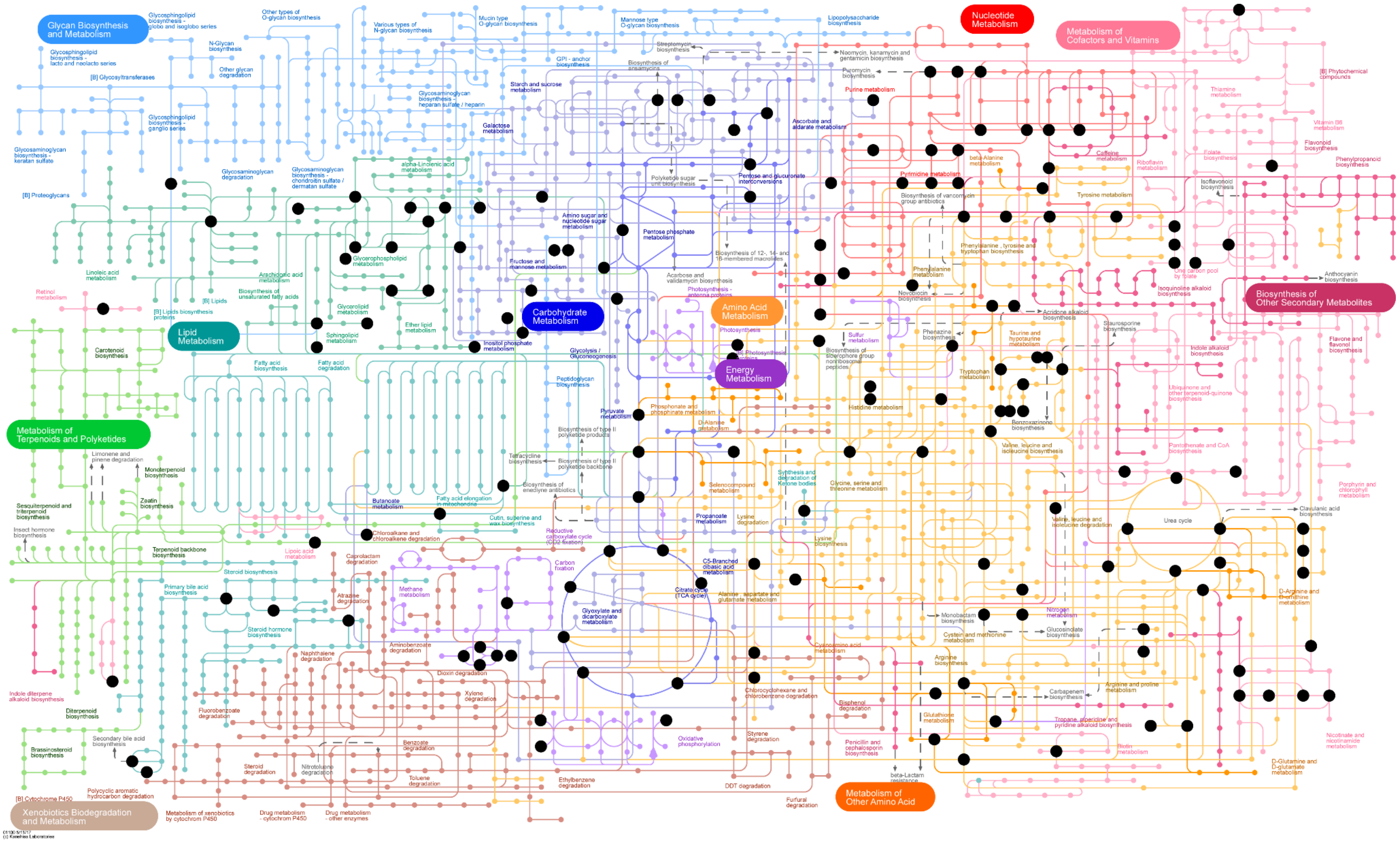

Glycan Biosynthesis and Metabolism
Lipid Metabolism
Metabolism of Terpenoids and Polyketides
Xenobiotics Biodegradation and Metabolism
Carbohydrate Metabolism
Amino Acid Metabolism
Energy Metabolism
Nucleotide Metabolism
Metabolism of Cofactors and Vitamins
Biosynthesis of Other Secondary Metabolites
Metabolism of Other Amino Acid
Glycosphingolipid biosynthesis - globo and isoglobo series
Glycosphingolipid biosynthesis - lacto and neolacto series
[B] Glycosyltransferases
Glycosphingolipid biosynthesis - ganglio series
Glycosaminoglycan biosynthesis - keratan sulfate
[B] Proteoglycans
N-Glycan biosynthesis
Other glycan degradation
Other types of O-glycan biosynthesis
Various types of N-glycan biosynthesis
Mucin type O-glycan biosynthesis
Mannose type O-glycan biosynthesis
Lipopolysaccharide biosynthesis
Glycosaminoglycan degradation
Glycosaminoglycan biosynthesis - chondroitin sulfate / dermatan sulfate
Glycosaminoglycan biosynthesis - heparan sulfate / heparin
Galactose metabolism
Starch and sucrose metabolism
GPI - anchor biosynthesis
Streptomycin biosynthesis
Biosynthesis of ansamycins
Neomycin, kanamycin and gentamicin biosynthesis
Puromycin biosynthesis
Purine metabolism
Ascorbate and aldarate metabolism
Pentose and glucuronate interconversions
Polyketide sugar unit biosynthesis
Pyrimidine metabolism
Biosynthesis of vancomycin group antibiotics
beta-Alanine metabolism
Caffeine metabolism
Riboflavin metabolism
Thiamine metabolism
Folate biosynthesis
Vitamin B6 metabolism
Flavonoid biosynthesis
[B] Phytochemical compounds
Phenylpropanoid biosynthesis
Isoflavonoid biosynthesis
Tyrosine metabolism
Phenylalanine, tyrosine and tryptophan biosynthesis
Phenylalanine metabolism
Novobiocin biosynthesis
One carbon pool by folate
Isoquinoline alkaloid biosynthesis
Anthocyanin biosynthesis
Acridone alkaloid biosynthesis
Staurosporine biosynthesis
Flavone and flavonol biosynthesis
Indole alkaloid biosynthesis
Ubiquinone and other terpenoid-quinone biosynthesis
Pantothenate and CoA biosynthesis
Porphyrin and chlorophyll metabolism
alpha-Linolenic acid metabolism
Linoleic acid metabolism
Arachidonic acid metabolism
Biosynthesis of unsaturated fatty acids
Glycerophospholipid metabolism
Glycerolipid metabolism
Ether lipid metabolism
Sphingolipid metabolism
[B] Lipids
[B] Lipids biosynthesis proteins
Retinol metabolism
Carotenoid biosynthesis
Fatty acid biosynthesis
Fatty acid degradation
Amino sugar and nucleotide sugar metabolism
Fructose and mannose metabolism
Pentose phosphate metabolism
Biosynthesis of 12-, 14- and 16-membered macrolides
Acarbose and validamycin biosynthesis
Photosynthesis - antenna proteins
Photosynthesis
[B] Photosynthesis proteins
Inositol phosphate metabolism
Glycolysis / Gluconeogenesis
Peptidoglycan biosynthesis
Pyruvate metabolism
Sulfur metabolism
Phenazine biosynthesis
Taurine and hypotaurine metabolism
Biosynthesis of siderophore group nonribosomal peptides
Tryptophan metabolism
Histidine metabolism
Benzoxazinoid biosynthesis
Valine, leucine and isoleucine biosynthesis
Phosphonate and phosphinate metabolism
D-Alanine metabolism
Biosynthesis of type II polyketide products
Tetracycline biosynthesis
Biosynthesis of type II polyketide backbone
Biosynthesis of enediyne antibiotics
Selenocompound metabolism
Synthesis and degradation of Ketone bodies
Glycine, serine and threonine metabolism
Monoterpenoid biosynthesis
Limonene and pinene degradation
Sesquiterpenoid and triterpenoid biosynthesis
Zeatin biosynthesis
Insect hormone biosynthesis
Terpenoid backbone biosynthesis
Butanoate metabolism
Fatty acid elongation in mitochondria
Cutin, suberine and wax biosynthesis
Chloroalkane and chloroalkene degradation
Reductive carboxylate cycle (CO2 fixation)
Propanoate metabolism
Lysine degradation
Lysine biosynthesis
Valine, leucine and isoleucine degradation
Urea cycle
Clavulanic acid biosynthesis
Lipoic acid metabolism
Steroid biosynthesis
Caprolactam degradation
Carbon fixation
C5-Branched dibasic acid metabolism
Primary bile acid biosynthesis
Methane metabolism
Atrazine degradation
Citrate cycle (TCA cycle)
Alanine, aspartate and glutamate metabolism
Glyoxylate and dicarboxylate metabolism
D-Arginine and D-ornithine metabolism
Steroid hormone biosynthesis
Aminobenzoate degradation
Naphthalene degradation
Nitrogen metabolism
Monobactam biosynthesis
Glucosinolate biosynthesis
Cysteine and methionine metabolism
Cyanoamino acid metabolism
Arginine biosynthesis
Dioxin degradation
Xylene degradation
Chlorocyclohexane and chlorobenzene degradation
Bisphenol degradation
Carbapenem biosynthesis
Arginine and proline metabolism
Indole diterpene alkaloid biosynthesis
Diterpenoid biosynthesis
Fluorobenzoate degradation
Glutathione metabolism
Tropane, piperidine and pyridine alkaloid biosynthesis
Nicotinate and nicotinamide metabolism
Secondary bile acid biosynthesis
Brassinosteroid biosynthesis
Benzoate degradation
Oxidative phosphorylation
Styrene degradation
Penicillin and cephalosporin biosynthesis
Biotin metabolism
Steroid degradation
Nitrotoluene degradation
Toluene degradation
D-Glutamine and D-glutamate metabolism
[B] Cytochrome P450
Polycyclic aromatic hydrocarbon degradation
Ethylbenzene degradation
DDT degradation
beta-Lactam resistance
Furfural degradation
Metabolism of xenobiotics by cytochrom P450
Drug metabolism - cytochrom P450
Drug metabolism - other enzymes
01100 5/13/17
(c) Kanehisa Laboratories

#### Supplementary Figure 5

Number of metabolites included in reference sample per biochemical pathway. Pathway names were obtained from Roche biochemical pathway map [ref 48] and are tabulated in **Sup.Table 8**. Abbreviations: Trp tryptophan, Arg arginine, Phe phenylalanine, Tyr tyrosine, Glu glutamic acid, Gln glutamine, His histidine, Asp aspartic acid, Asn asparagine, Pro proline, Ser serine, NAD nicotinamide adenine dinucleotide.

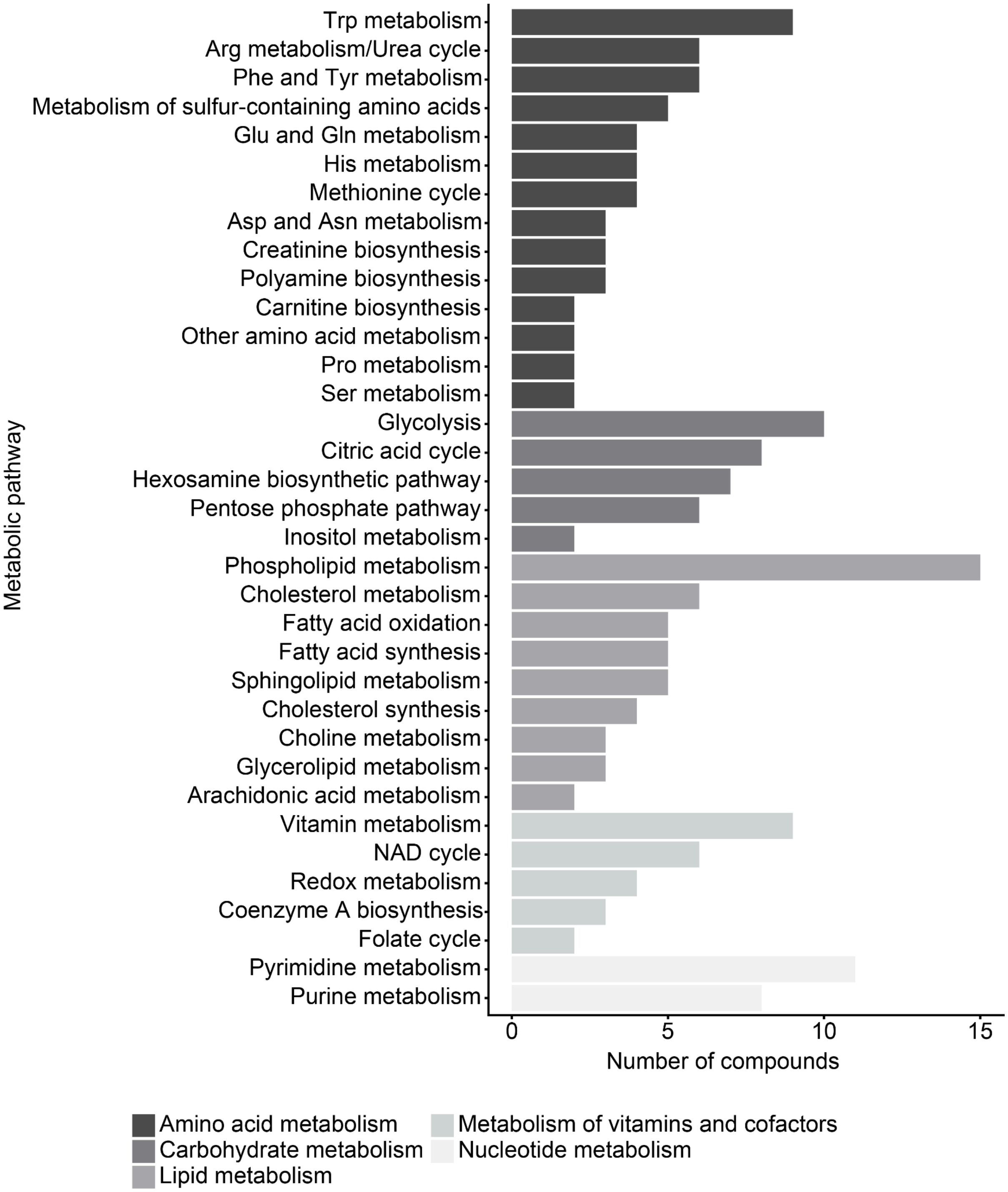

#### Supplementary Figure 6

Number of metabolites included in reference sample per chemical subclass, coloured by chemical class, according to custom classification in **Sup.Table 9.**

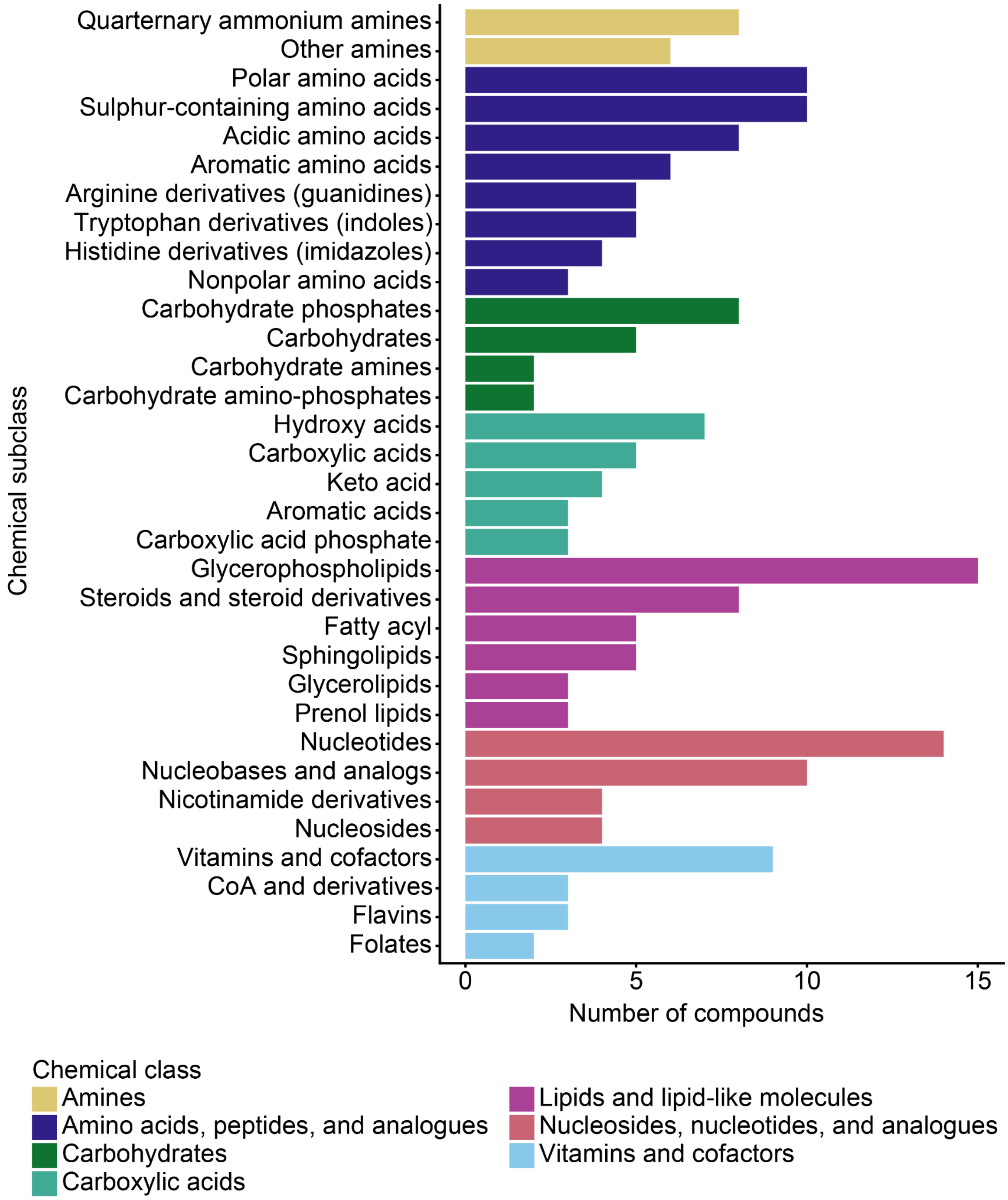

### Supplementary Figure 7

Illustration of the natural abundance of E. coli metabolites by [19] with reference metabolites highlighted, coloured by reference metabolite chemical class.

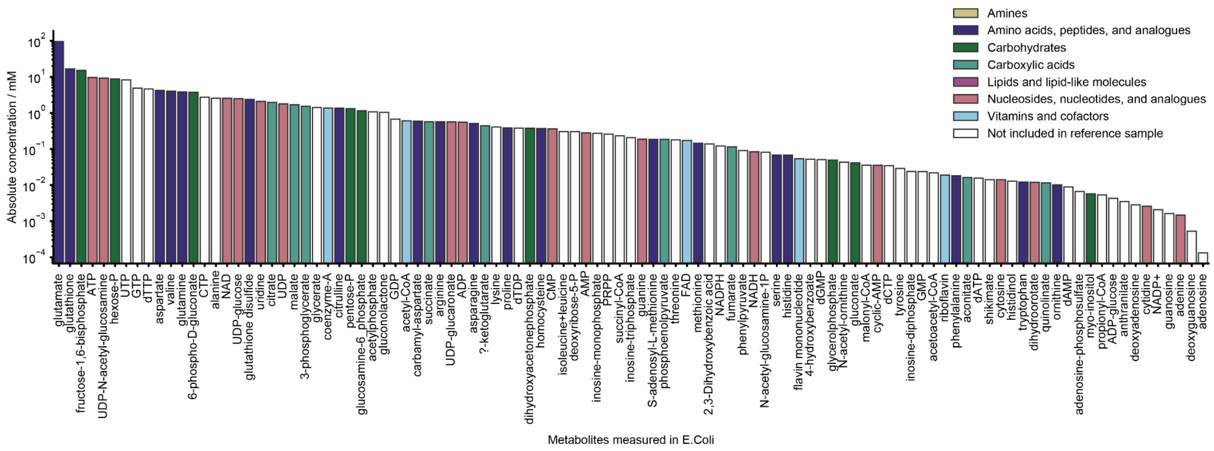

#### Supplementary Figure 8

Number of MALDI-MS protocols (out of 24) detecting each reference metabolite. One bar represents one metabolite.

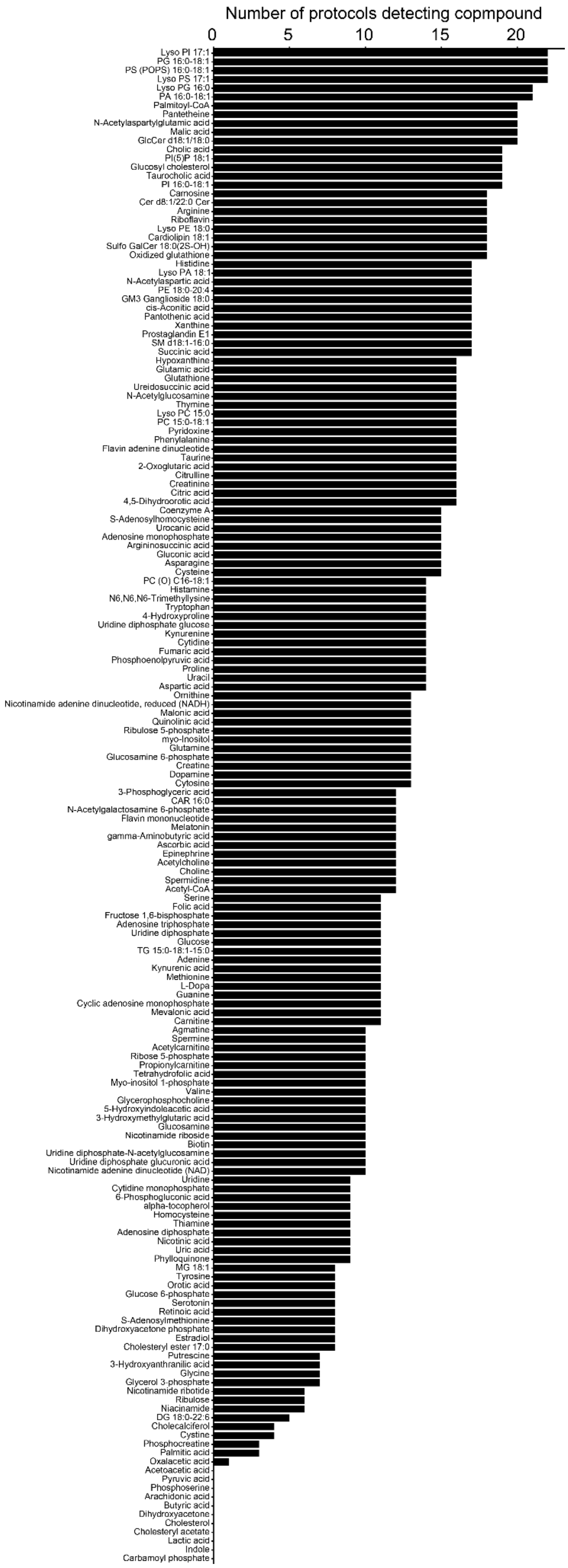
Number of protocols detecting copmpound
0
5
10
15
20
Lyso PI 17:1
PG 16:0-18:1
PS (POPS) 16:0-18:1
Lyso PS 17:1
Lyso PG 16:0
PA 16:0-18:1
Palmitoyl-CoA
Pantetheine
N-Acetylaspartylglutamic acid
Malic acid
GlcCer d18:1/18:0
Cholic acid
PI(5)P 18:1
Glucosyl cholesterol
Taurocholic acid
PI 16:0-18:1
Carnosine
Cer d8:1/22:0 Cer
Arginine
Riboflavin
Lyso PE 18:0
Cardiolipin 18:1
Sulfo GalCer 18:0(2S-OH)
Oxidized glutathione
Histidine
Lyso PA 18:1
N-Acetylaspartic acid
PE 18:0-20:4
GM3 Ganglioside 18:0
cis-Aconitic acid
Pantothenic acid
Xanthine
Prostaglandin E1
SM d18:1-16:0
Succinic acid
Hypoxanthine
Glutamic acid
Glutathione
Ureidosuccinic acid
N-Acetylglucosamine
Thymine
Lyso PC 15:0
PC 15:0-18:1
Pyridoxine
Phenylalanine
Flavin adenine dinucleotide
Taurine
2-Oxoglutaric acid
Citrulline
Creatinine
Citric acid
4,5-Dihydroorotic acid
Coenzyme A
S-Adenosylhomocysteine
Urocanic acid
Adenosine monophosphate
Argininosuccinic acid
Gluconic acid
Asparagine
Cysteine
PC (O) C16-18:1
Histamine
N6,N6,N6-Trimethyllysine
Tryptophan
4-Hydroxyproline
Uridine diphosphate glucose
Kynurenine
Cytidine
Fumaric acid
Phosphoenolpyruvic acid
Proline
Uracil
Aspartic acid
Ornithine
Nicotinamide adenine dinucleotide, reduced (NADH)
Malonic acid
Quinolinic acid
Ribulose 5-phosphate
myo-Inositol
Glutamine
Glucosamine 6-phosphate
Creatine
Dopamine
Cytosine
3-Phosphoglyceric acid
CAR 16:0
N-Acetylgalactosamine 6-phosphate
Flavin mononucleotide
Melatonin
gamma-Aminobutyric acid
Ascorbic acid
Epinephrine
Acetylcholine
Choline
Spermidine
Acetyl-CoA
Serine
Folic acid
Fructose 1,6-bisphosphate
Adenosine triphosphate
Uridine diphosphate
Glucose
TG 15:0-18:1-15:0
Adenine
Kynurenic acid
Methionine
L-Dopa
Guanine
Cyclic adenosine monophosphate
Mevalonic acid
Carnitine
Agmatine
Spermine
Acetylcarnitine
Ribose 5-phosphate
Propionylcarnitine
Tetrahydrofolic acid
Myo-inositol 1-phosphate
Valine
Glycerophosphocholine
5-Hydroxyindoleacetic acid
3-Hydroxymethylglutaric acid
Glucosamine
Nicotinamide riboside
Biotin
Uridine diphosphate-N-acetylglucosamine
Uridine diphosphate glucuronic acid
Nicotinamide adenine dinucleotide (NAD)
Uridine
Cytidine monophosphate
6-Phosphogluconic acid
alpha-tocopherol
Homocysteine
Thiamine
Adenosine diphosphate
Nicotinic acid
Uric acid
Phylloquinone
MG 18:1
Tyrosine
Orotic acid
Glucose 6-phosphate
Serotonin
Retinoic acid
S-Adenosylmethionine
Dihydroxyacetone phosphate
Estradiol
Cholesteryl ester 17:0
Putrescine
3-Hydroxyanthranilic acid
Glycine
Glycerol 3-phosphate
Nicotinamide ribotide
Ribulose
Niacinamide
DG 18:0-22:6
Cholecalciferol
Cystine
Phosphocreatine
Palmitic acid
Oxalacetic acid
Acetoacetic acid
Pyruvic acid
Phosphoserine
Arachidonic acid
Butyric acid
Dihydroxyacetone
Cholesterol
Cholesteryl acetate
Lactic acid
Indole
Carbamoyl phosphate

#### Supplementary Figure 9

Number of reference metabolites detected in each chemical class by at least one protocol, plotted separate for each ionisation mode.

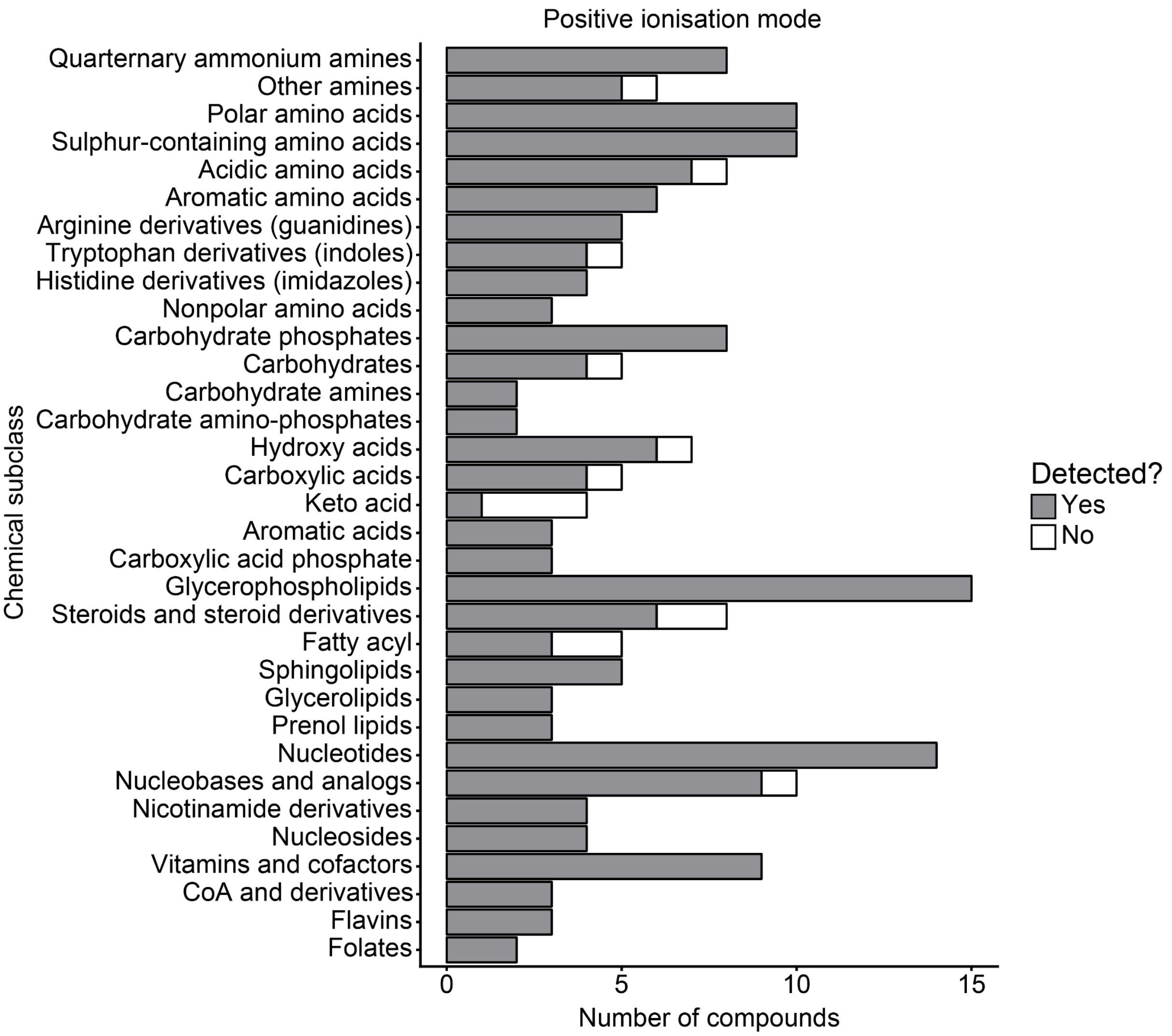

Positive ionisation mode
Chemical subclass
Quarternary ammonium amines
Other amines
Polar amino acids
Sulphur-containing amino acids
Acidic amino acids
Aromatic amino acids
Arginine derivatives (guanidines)
Tryptophan derivatives (indoles)
Histidine derivatives (imidazoles)
Nonpolar amino acids
Carbohydrate phosphates
Carbohydrates
Carbohydrate amines
Carbohydrate amino-phosphates
Hydroxy acids
Carboxylic acids
Keto acid
Aromatic acids
Carboxylic acid phosphate
Glycerophospholipids
Steroids and steroid derivatives
Fatty acyl
Sphingolipids
Glycerolipids
Prenol lipids
Nucleotides
Nucleobases and analogs
Nicotinamide derivatives
Nucleosides
Vitamins and cofactors
CoA and derivatives
Flavins
Folates
0
5
10
15
Number of compounds
Detected?
Yes
No

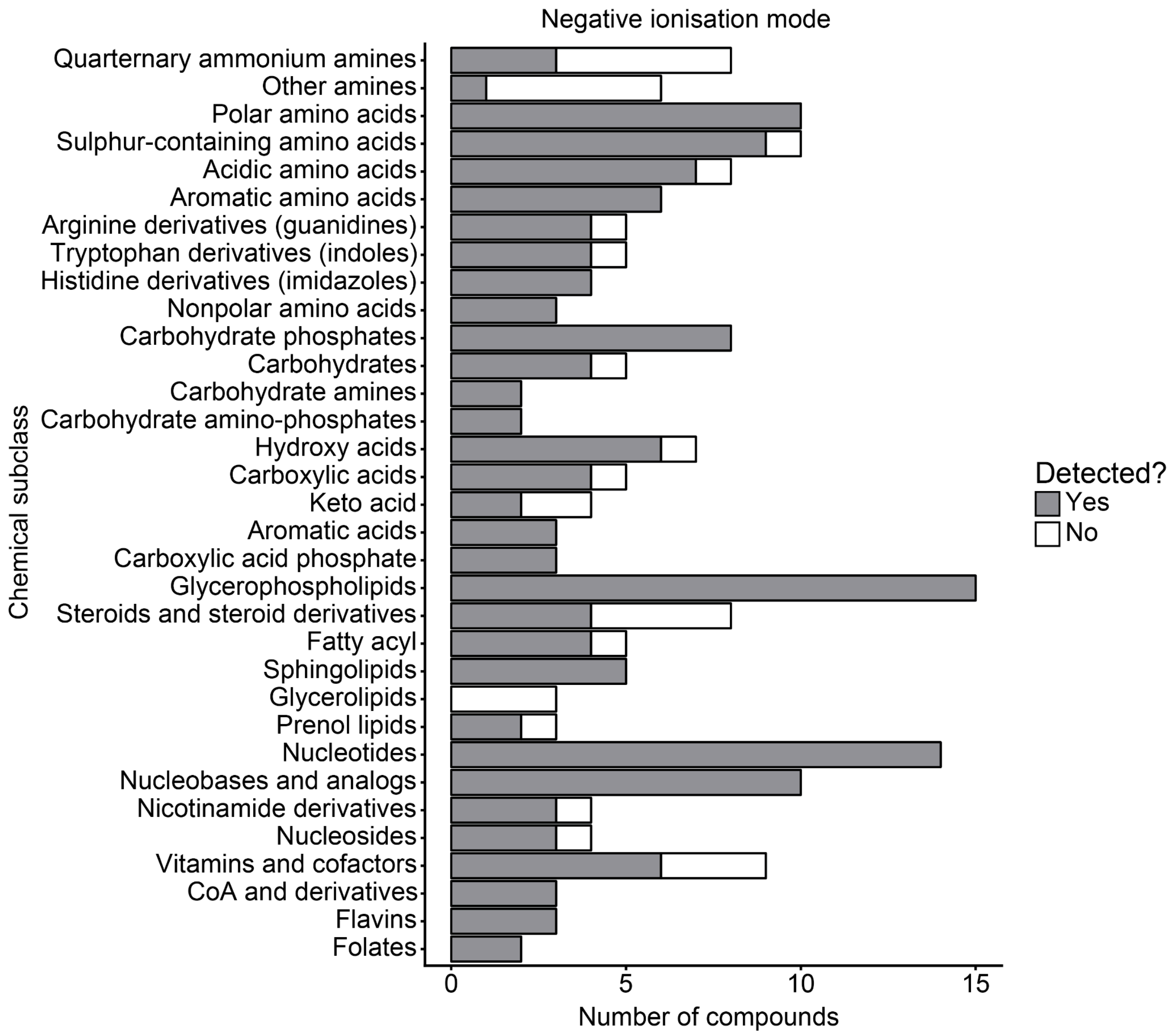
Negative ionisation mode
Chemical subclass
Quarternary ammonium amines
Other amines
Polar amino acids
Sulphur-containing amino acids
Acidic amino acids
Aromatic amino acids
Arginine derivatives (guanidines)
Tryptophan derivatives (indoles)
Histidine derivatives (imidazoles)
Nonpolar amino acids
Carbohydrate phosphates
Carbohydrates
Carbohydrate amines
Carbohydrate amino-phosphates
Hydroxy acids
Carboxylic acids
Keto acid
Aromatic acids
Carboxylic acid phosphate
Glycerophospholipids
Steroids and steroid derivatives
Fatty acyl
Sphingolipids
Glycerolipids
Prenol lipids
Nucleotides
Nucleobases and analogs
Nicotinamide derivatives
Nucleosides
Vitamins and cofactors
CoA and derivatives
Flavins
Folates
0
5
10
15
Number of compounds
Detected?
Yes
No

#### Supplementary Figure 10

Detection of metabolites in chemical classes and subclasses by different MALDI-MS protocols. Dot size shows the fraction of metabolites detected in a given pathways among those considered (note that not all intermediates per pathway were considered); colour shows the average intensity among the detected ions.

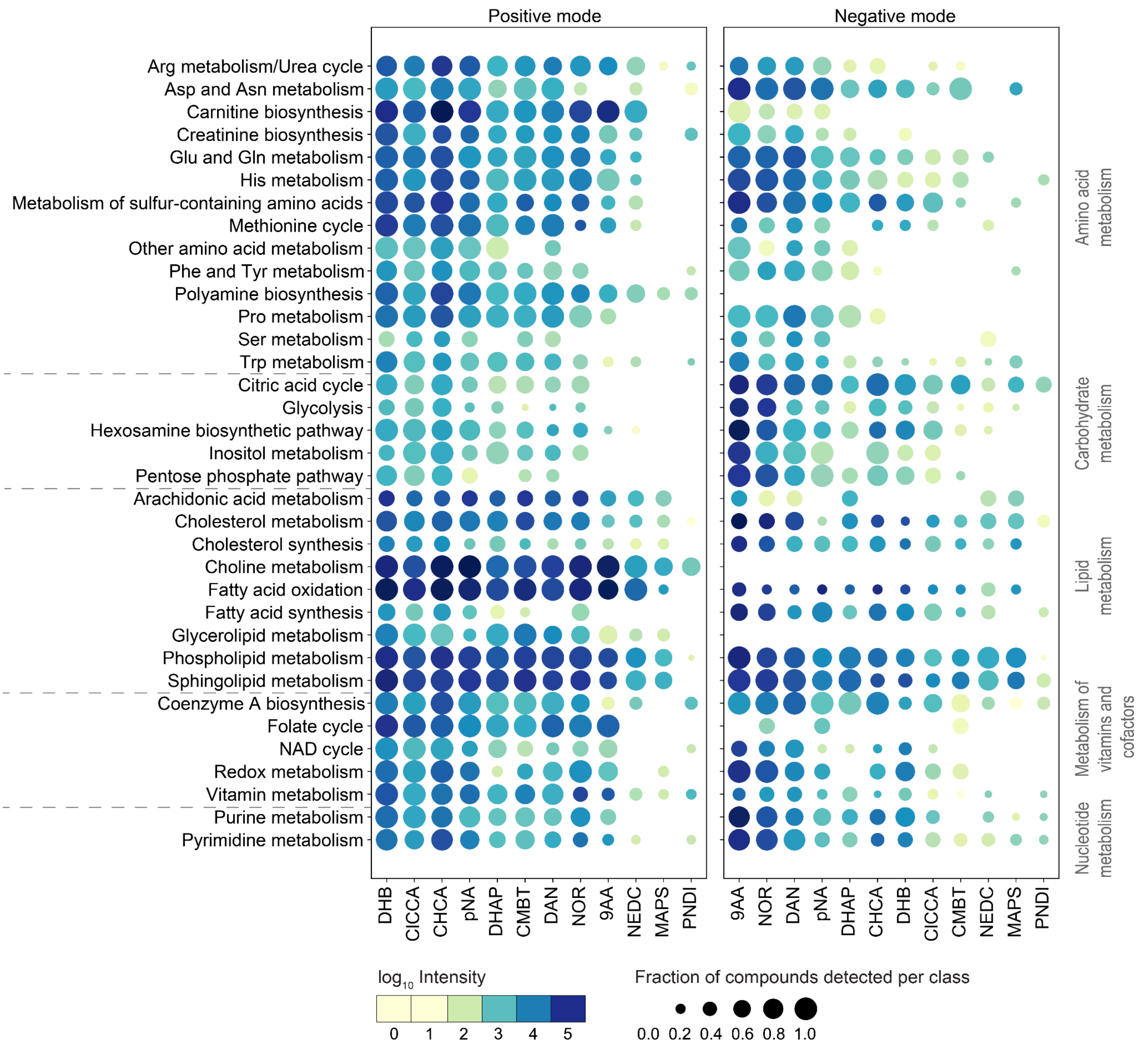

#### Supplementary Figure 11

Fraction of metabolites detected per chemical subclass in the reference sample and in biological tissues, using DHB positive mode, and DAN negative mode protocols. One dot represents a chemical subclass.

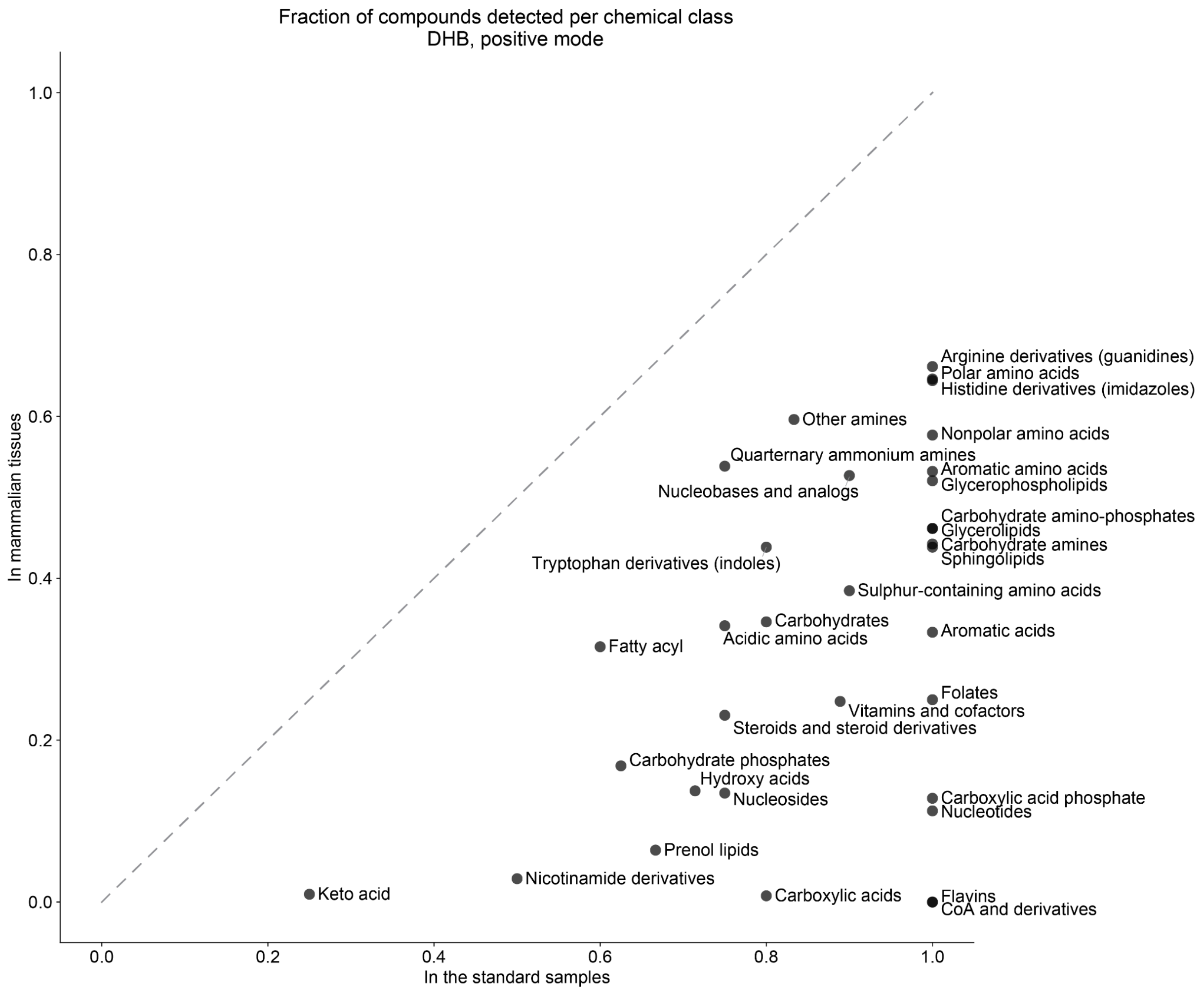

Fraction of compounds detected per chemical class
DHB, positive mode
In mammalian tissues
In the standard samples
0.0
0.2
0.4
0.6
0.8
1.0
Arginine derivatives (guanidines)
Polar amino acids
Histidine derivatives (imidazoles)
Other amines
Nonpolar amino acids
Quarternary ammonium amines
Aromatic amino acids
Glycerophospholipids
Nucleobases and analogs
Carbohydrate amino-phosphates
Glycerolipids
Carbohydrate amines
Sphingolipids
Tryptophan derivatives (indoles)
Sulphur-containing amino acids
Carbohydrates
Acidic amino acids
Aromatic acids
Fatty acyl
Folates
Vitamins and cofactors
Steroids and steroid derivatives
Carbohydrate phosphates
Hydroxy acids
Nucleosides
Carboxylic acid phosphate
Nucleotides
Prenol lipids
Nicotinamide derivatives
Keto acid
Carboxylic acids
Flavins
CoA and derivatives

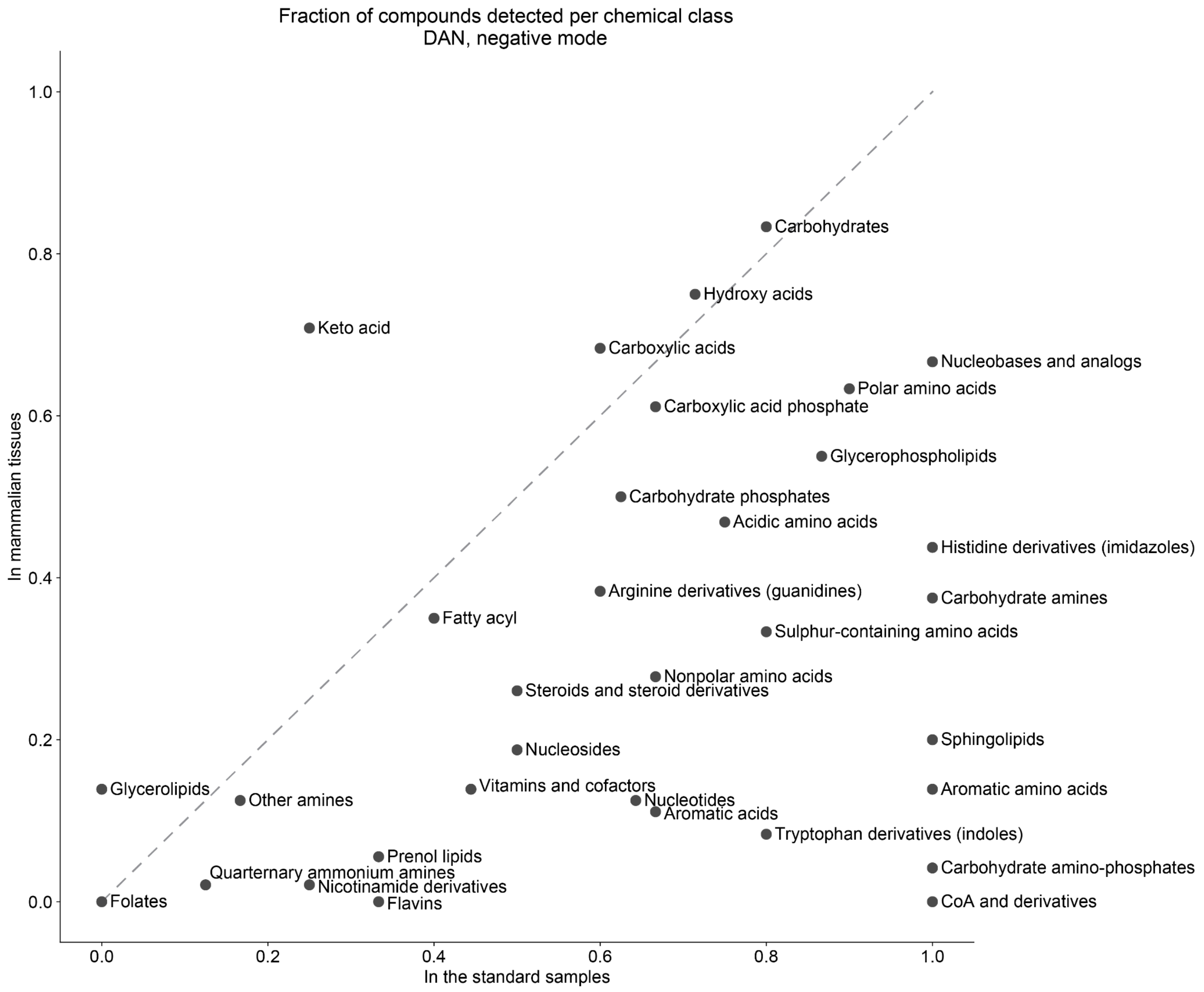
Fraction of compounds detected per chemical class
DAN, negative mode
In mammalian tissues
In the standard samples
0.0
0.2
0.4
0.6
0.8
1.0
Carbohydrates
Hydroxy acids
Keto acid
Carboxylic acids
Nucleobases and analogs
Polar amino acids
Carboxylic acid phosphate
Glycerophospholipids
Carbohydrate phosphates
Acidic amino acids
Histidine derivatives (imidazoles)
Arginine derivatives (guanidines)
Carbohydrate amines
Fatty acyl
Sulphur-containing amino acids
Nonpolar amino acids
Steroids and steroid derivatives
Nucleosides
Sphingolipids
Glycerolipids
Vitamins and cofactors
Aromatic amino acids
Other amines
Nucleotides
Aromatic acids
Tryptophan derivatives (indoles)
Prenol lipids
Carbohydrate amino-phosphates
Quarternary ammonium amines
Nicotinamide derivatives
Folates
Flavins
CoA and derivatives

#### Supplementary Figure 12

Summary of the SHAP feature importance analysis for classification and regression models. One Matrix-polarity combination is one model.

**SHAP scores for classification models, positive polarity**

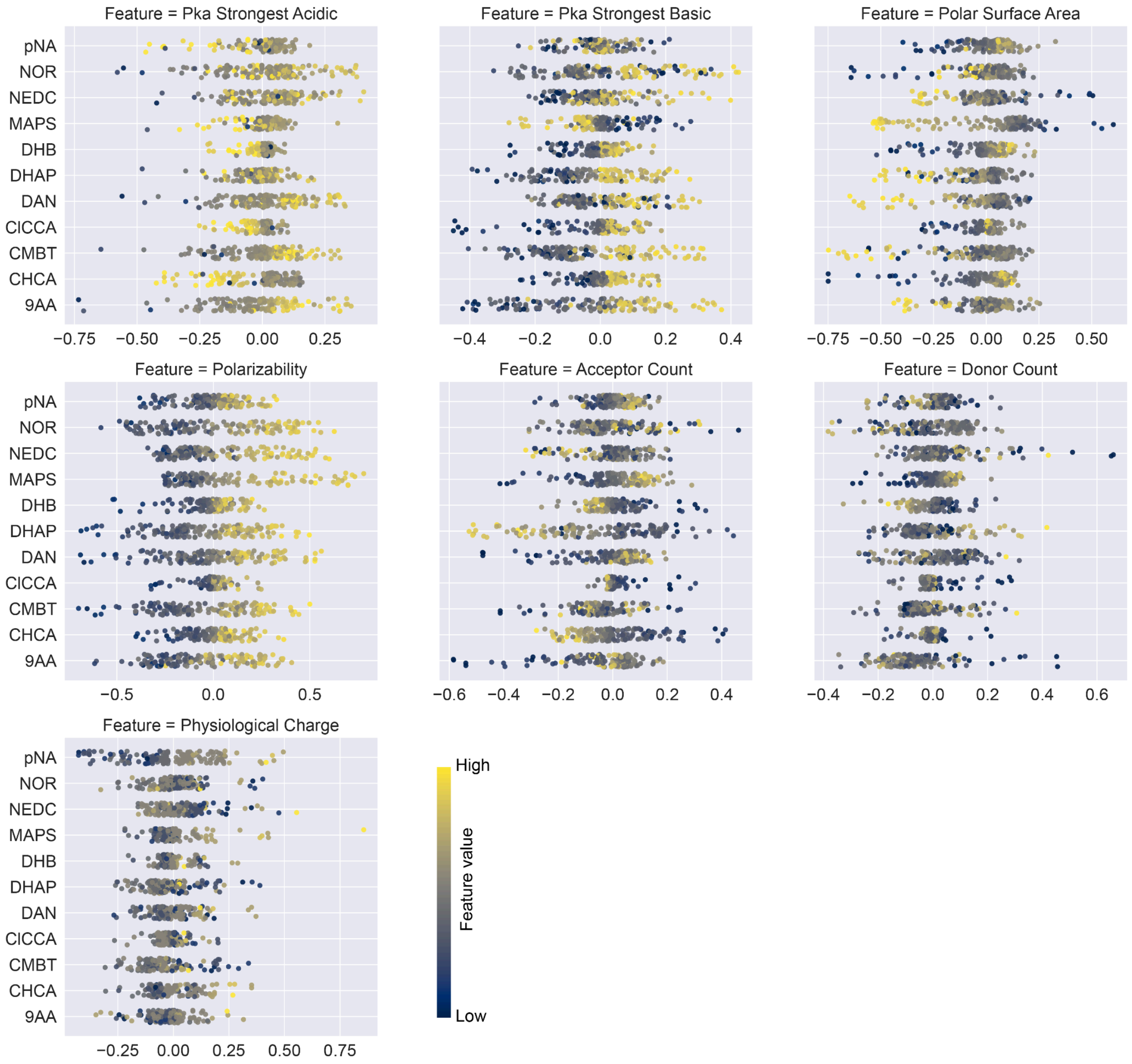

**SHAP scores for classification models, negative polarity**

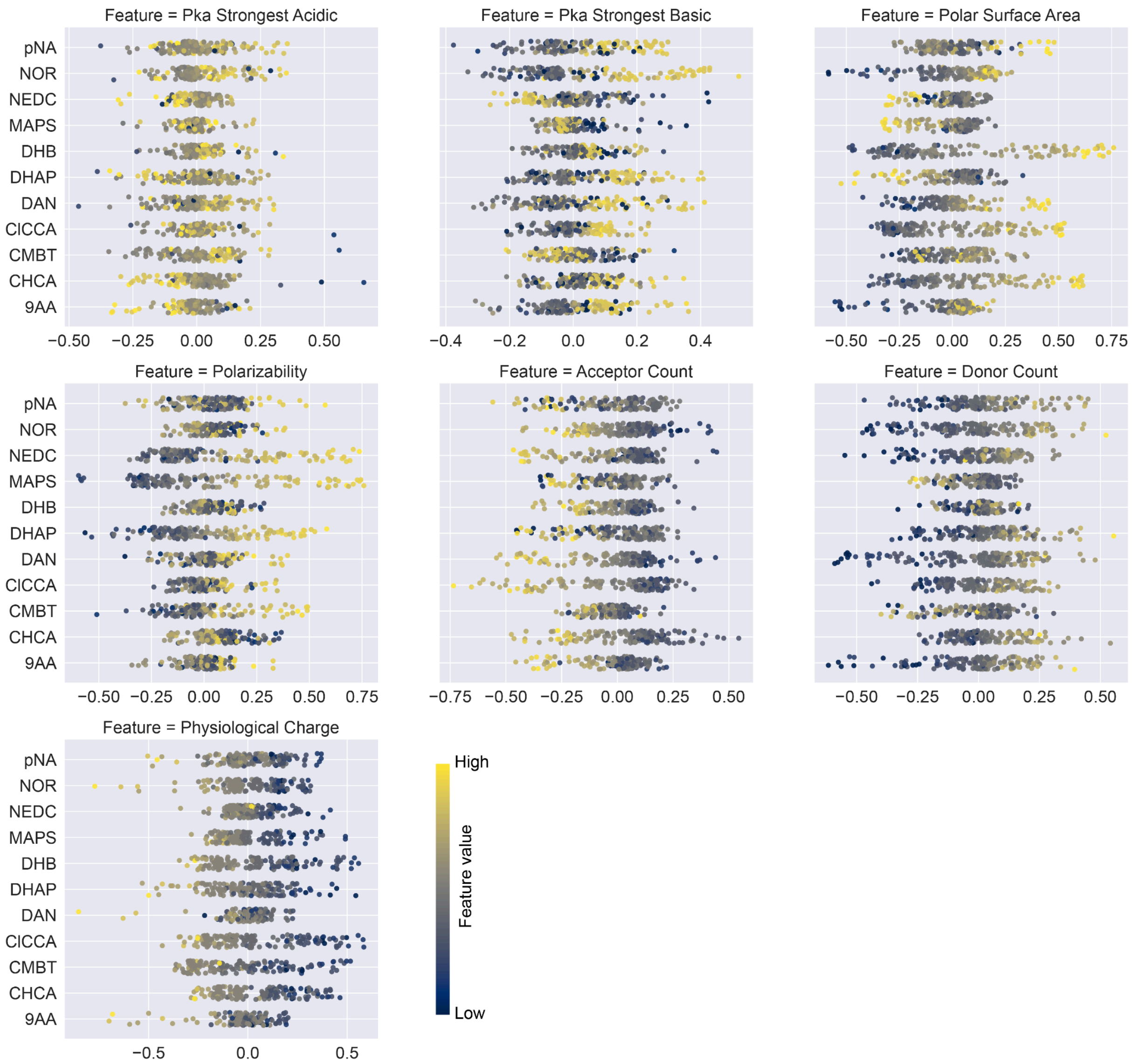

**SHAP scores for regression models, positive polarity**

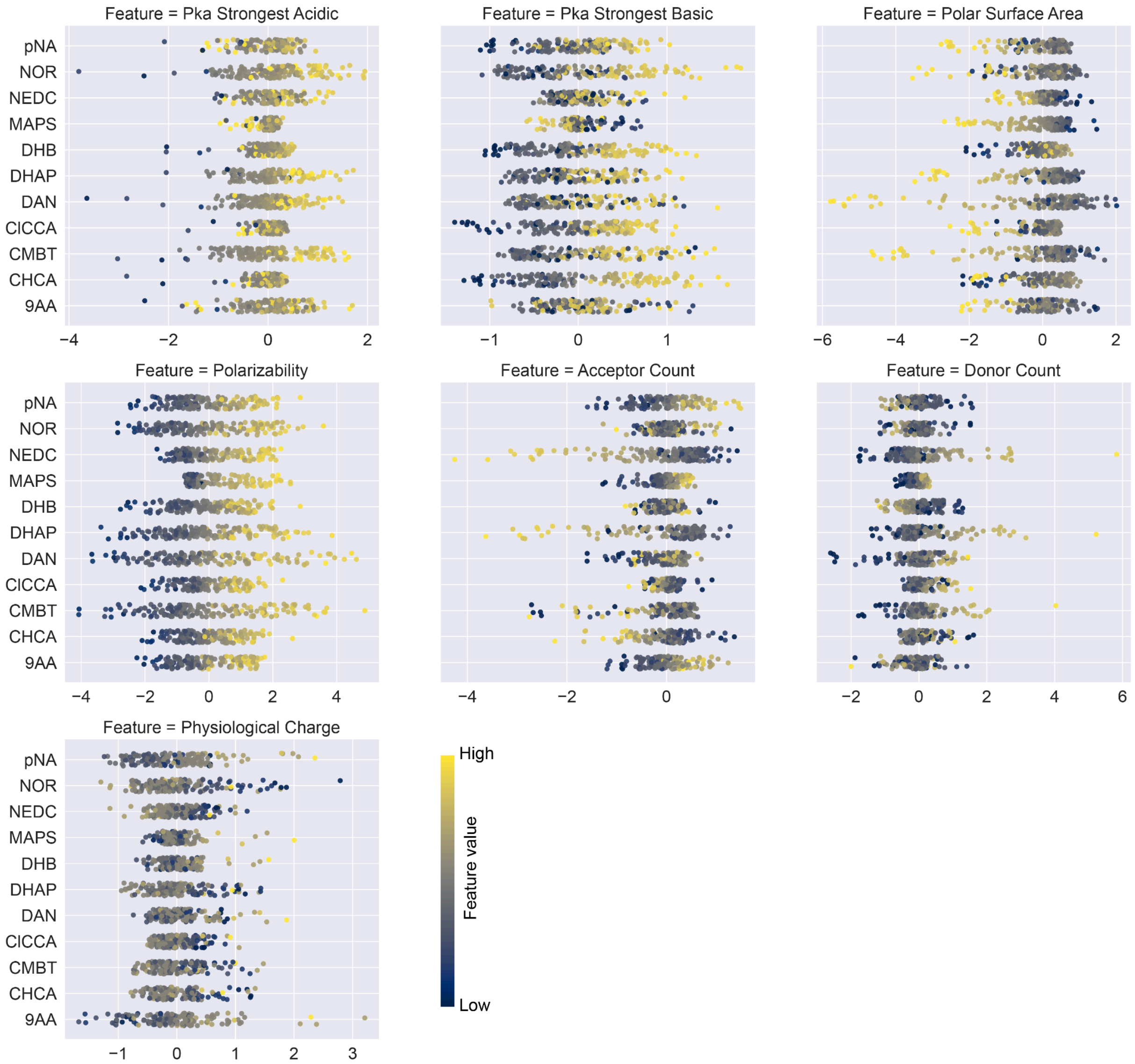

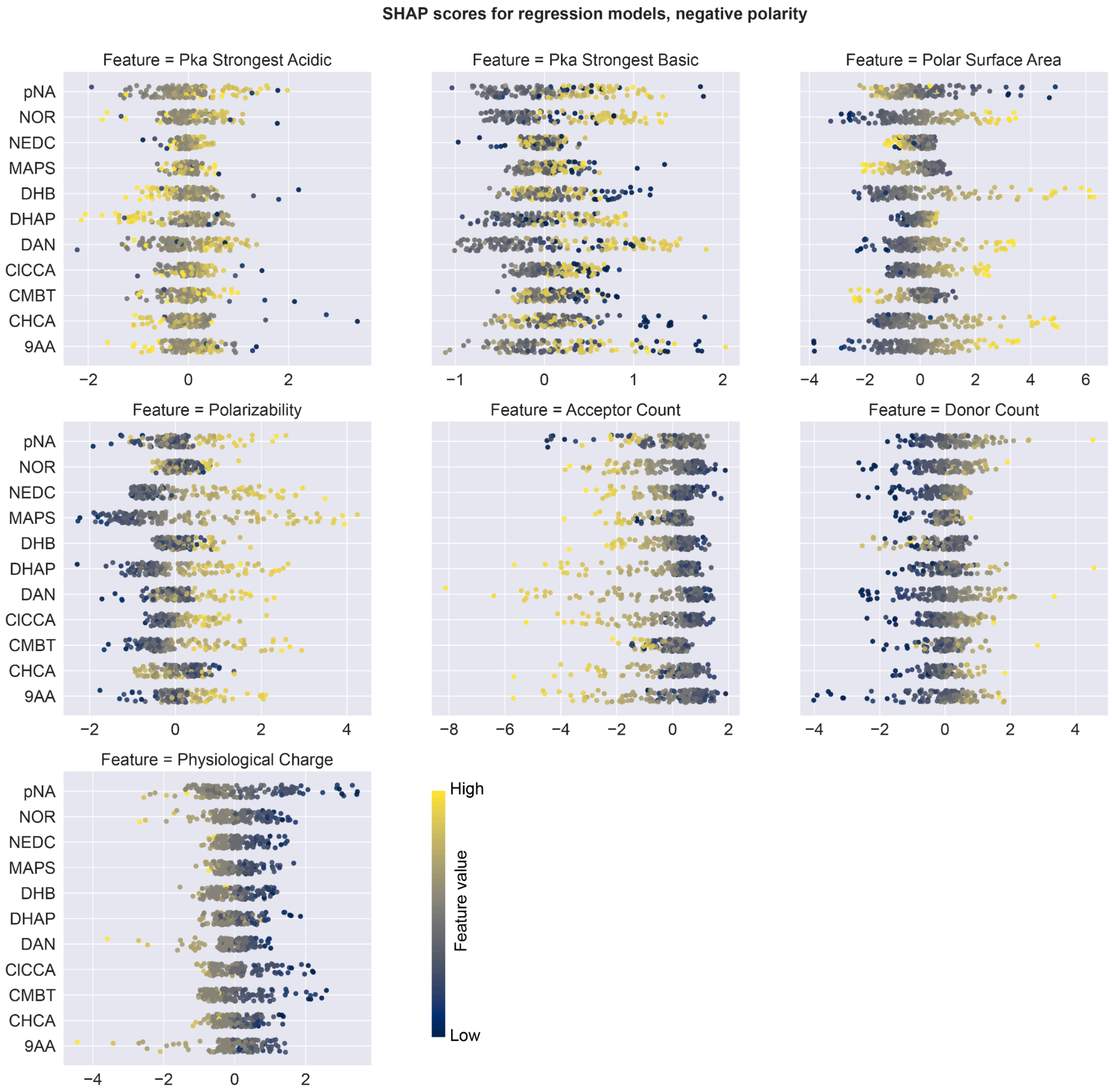

SHAP scores for regression models, negative polarity
Feature = Pka Strongest Acidic
Feature = Pka Strongest Basic
Feature = Polar Surface Area
Feature = Polarizability
Feature = Acceptor Count
Feature = Donor Count
Feature = Physiological Charge
pNA
NOR
NEDC
MAPS
DHB
DHAP
DAN
ClCCA
CMBT
CHCA
9AA
High
Feature value
Low

#### Supplementary Figure 13

Number of public data sets per each ionisation source, MALDI matrix, and mass analyser, available on METASPACE (total number of data sets was 7787, accessed on 03/03/2023).

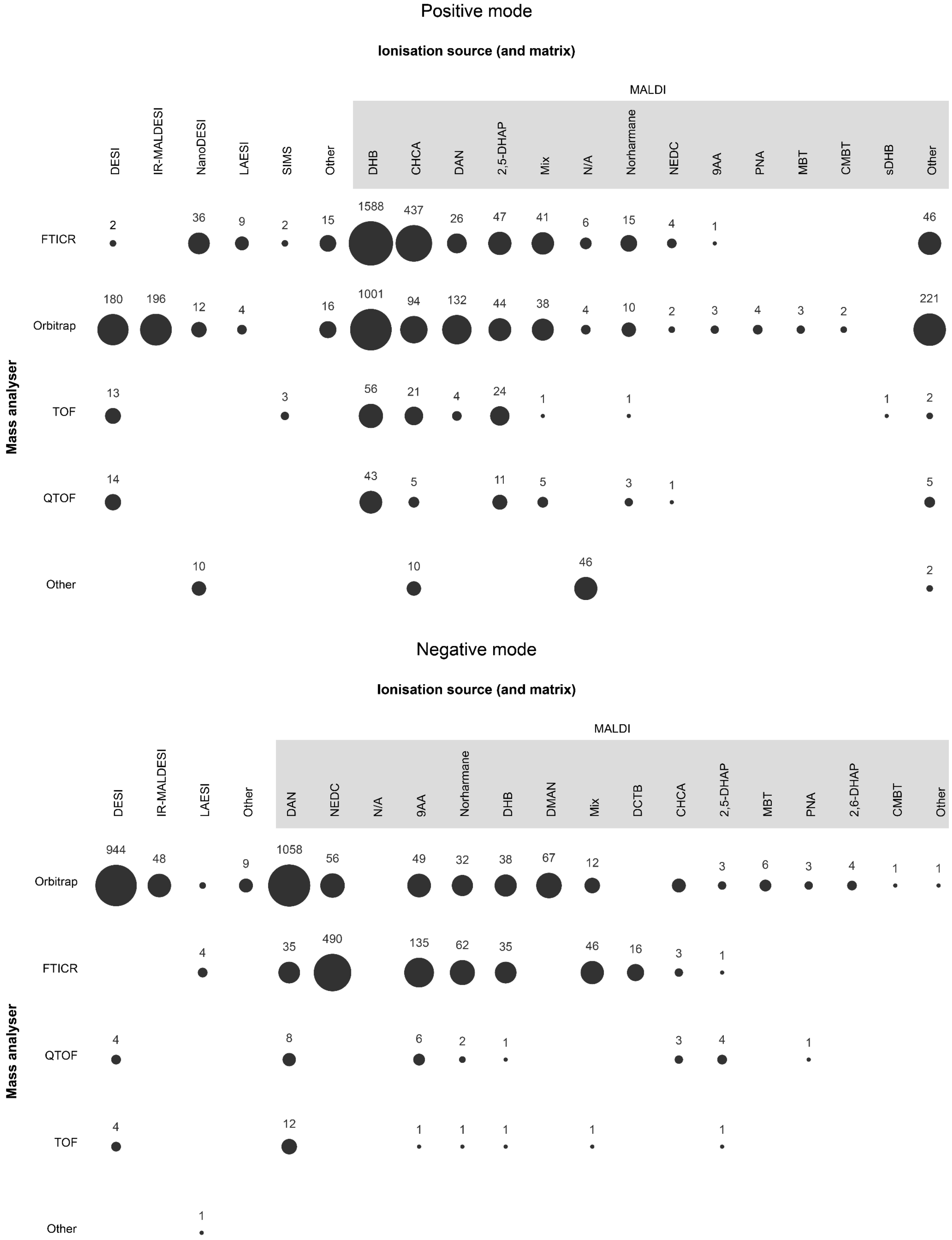

#### Supplementary Figure 14

Comparison of imaging MS instruments based on the detectability of reference metabolites, in positive and negative modes (separate panels).

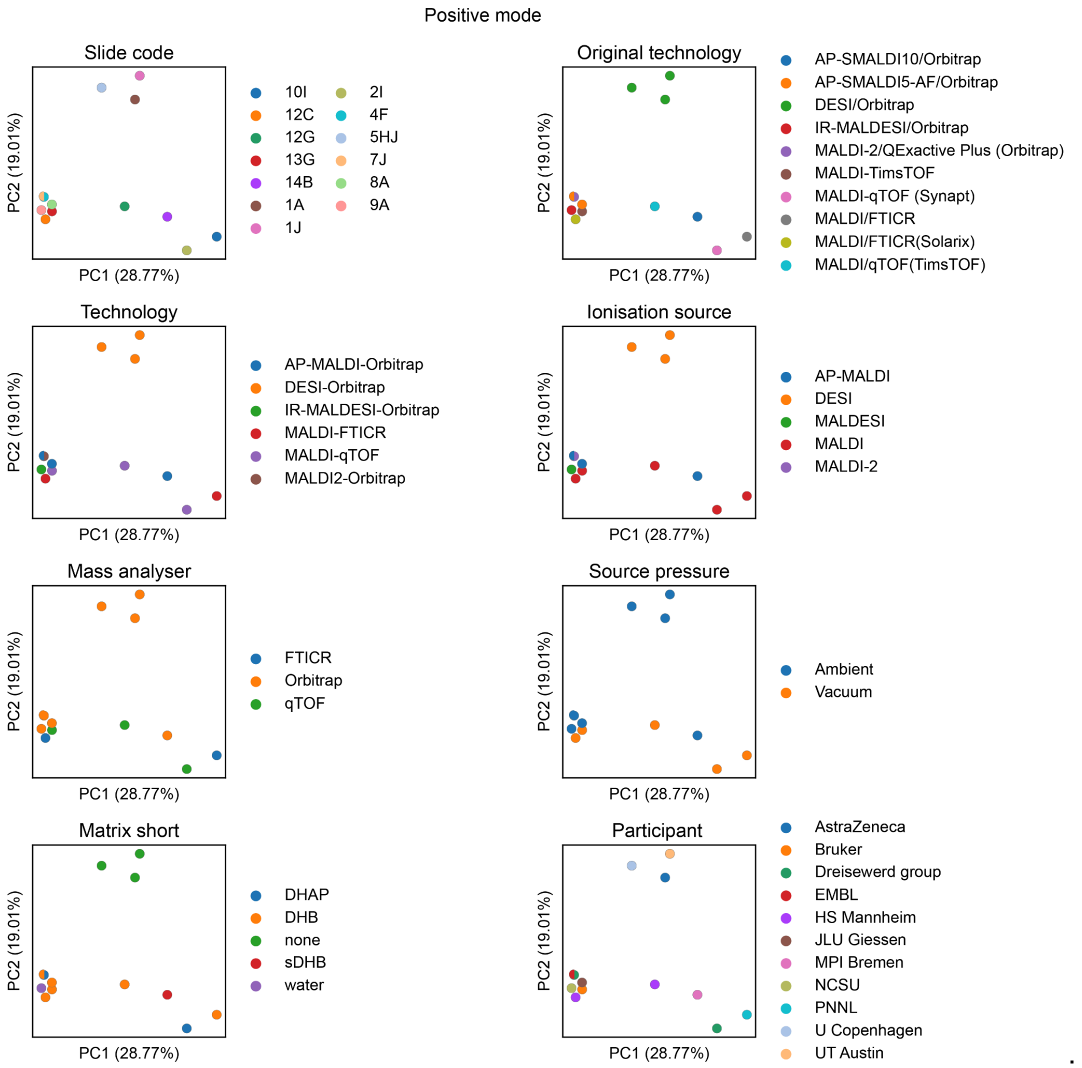
Positive mode
Slide code
PC2 (19.01%)
PC1 (28.77%)
10I
12C
12G
13G
14B
1A
1J
2I
4F
5HJ
7J
8A
9A
Original technology
AP-SMALDI10/Orbitrap
AP-SMALDI5-AF/Orbitrap
DESI/Orbitrap
IR-MALDESI/Orbitrap
MALDI-2/QExactive Plus (Orbitrap)
MALDI-TimsTOF
MALDI-qTOF (Synapt)
MALDI/FTICR
MALDI/FTICR(Solarix)
MALDI/qTOF(TimsTOF)
Technology
AP-MALDI-Orbitrap
DESI-Orbitrap
IR-MALDESI-Orbitrap
MALDI-FTICR
MALDI-qTOF
MALDI2-Orbitrap
Ionisation source
AP-MALDI
DESI
MALDESI
MALDI
MALDI-2
Mass analyser
FTICR
Orbitrap
qTOF
Source pressure
Ambient
Vacuum
Matrix short
DHAP
DHB
none
sDHB
water
Participant
AstraZeneca
Bruker
Dreisewerd group
EMBL
HS Mannheim
JLU Giessen
MPI Bremen
NCSU
PNNL
U Copenhagen
UT Austin

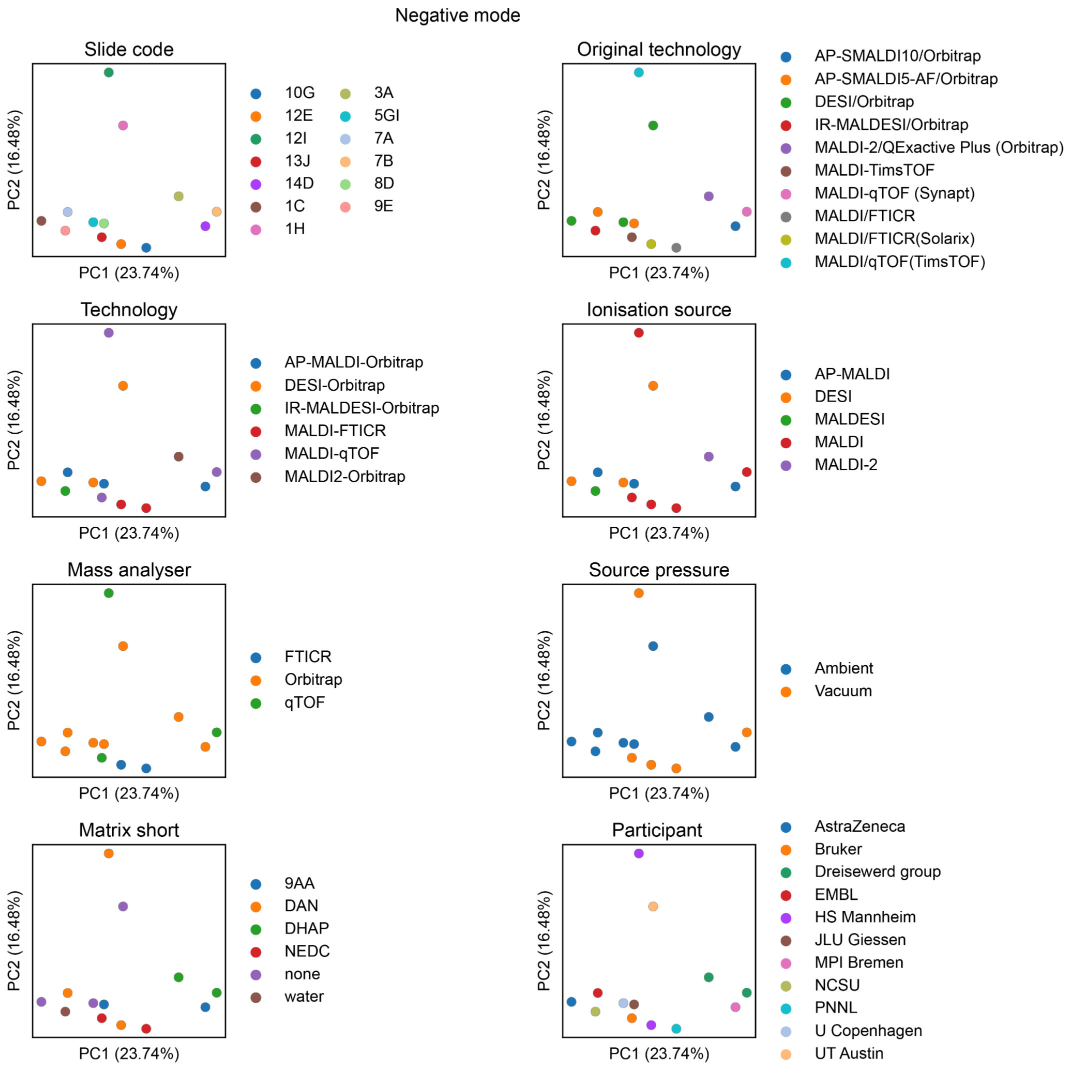
Negative mode
Slide code
PC2 (16.48%)
PC1 (23.74%)
10G
12E
12I
13J
14D
1C
1H
3A
5GI
7A
7B
8D
9E
Original technology
AP-SMALDI10/Orbitrap
AP-SMALDI5-AF/Orbitrap
DESI/Orbitrap
IR-MALDESI/Orbitrap
MALDI-2/QExactive Plus (Orbitrap)
MALDI-TimsTOF
MALDI-qTOF (Synapt)
MALDI/FTICR
MALDI/FTICR(Solarix)
MALDI/qTOF(TimsTOF)
Technology
AP-MALDI-Orbitrap
DESI-Orbitrap
IR-MALDESI-Orbitrap
MALDI-FTICR
MALDI-qTOF
MALDI2-Orbitrap
Ionisation source
AP-MALDI
DESI
MALDESI
MALDI
MALDI-2
Mass analyser
FTICR
Orbitrap
qTOF
Source pressure
Ambient
Vacuum
Matrix short
9AA
DAN
DHAP
NEDC
none
water
Participant
AstraZeneca
Bruker
Dreisewerd group
EMBL
HS Mannheim
JLU Giessen
MPI Bremen
NCSU
PNNL
U Copenhagen
UT Austin

#### Supplementary Figure 15

Proportional contribution of different ion types to the total detected metabolite signal, averaged per chemical class, and averaged per polarity and technology. Contributions below 5% are consolidated and labelled as "Other". Please note that [M]- radical may be confused with the first $^{13}C$ isotope of deprotonated parent ion.

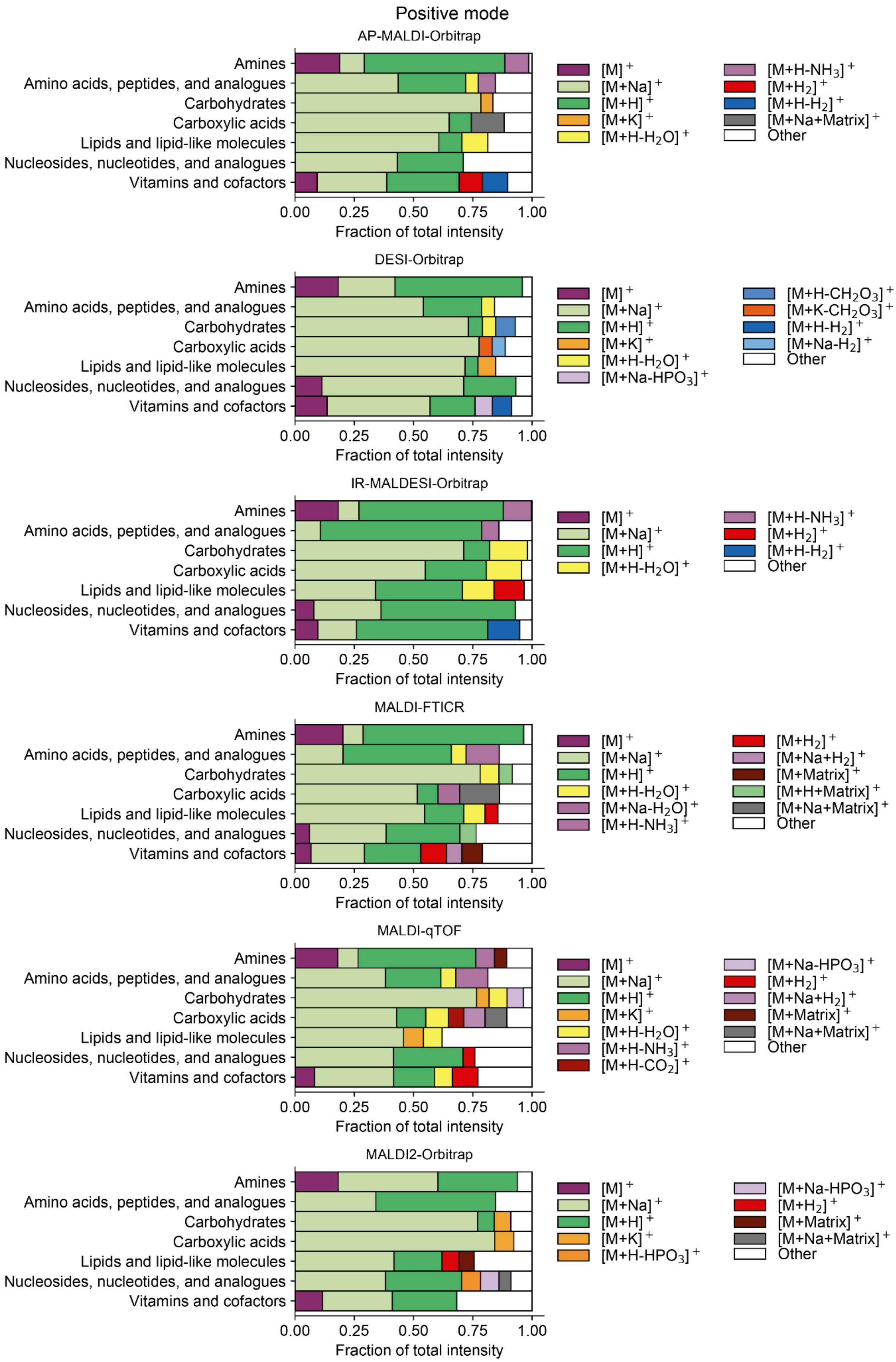
Positive mode
AP-MALDI-Orbitrap
DESI-Orbitrap
IR-MALDESI-Orbitrap
MALDI-FTICR
MALDI-qTOF
MALDI2-Orbitrap
Amines
Amino acids, peptides, and analogues
Carbohydrates
Carboxylic acids
Lipids and lipid-like molecules
Nucleosides, nucleotides, and analogues
Vitamins and cofactors
0.00
0.25
0.50
0.75
1.00
Fraction of total intensity
[M]+
[M+Na]+
[M+H]+
[M+K]+
[M+H-H2O]+
[M+H-NH3]+
[M+H2]+
[M+H-H2]+
[M+Na+Matrix]+
[M+Na-HPO3]+
[M+H-CH2O3]+
[M+K-CH2O3]+
[M+Na-H2]+
[M+Na-H2O]+
[M+Na+H2]+
[M+Matrix]+
[M+H+Matrix]+
[M+H-CO2]+
[M+H-HPO3]+
Other

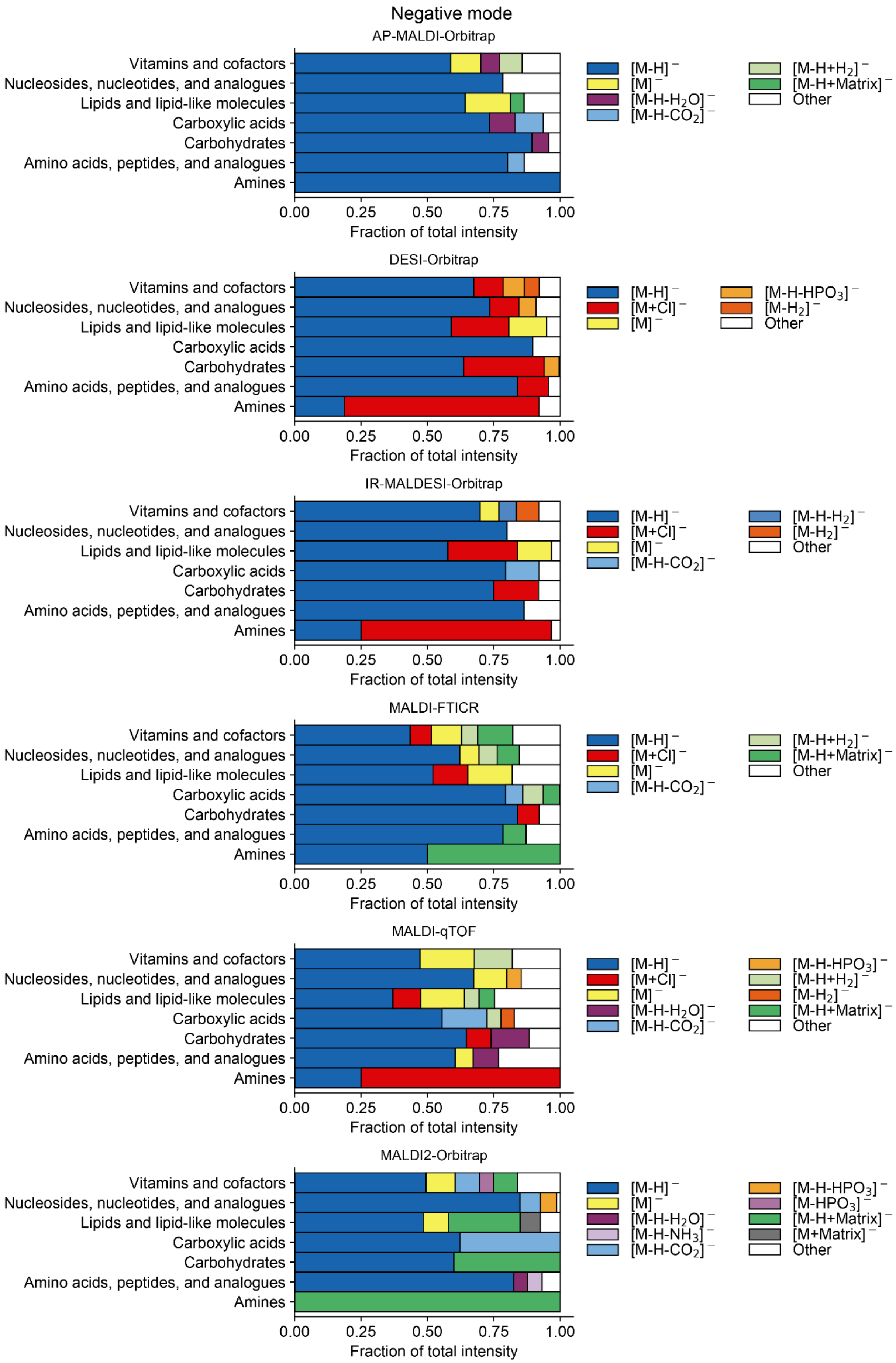

Negative mode
AP-MALDI-Orbitrap
Vitamins and cofactors
Nucleosides, nucleotides, and analogues
Lipids and lipid-like molecules
Carboxylic acids
Carbohydrates
Amino acids, peptides, and analogues
Amines
0.00 0.25 0.50 0.75 1.00
Fraction of total intensity
$[M-H]^-$
$[M]^-$
$[M-H-H_2O]^-$
$[M-H-CO_2]^-$
$[M-H+H_2]^-$
$[M-H+Matrix]^-$
Other
DESI-Orbitrap
Vitamins and cofactors
Nucleosides, nucleotides, and analogues
Lipids and lipid-like molecules
Carboxylic acids
Carbohydrates
Amino acids, peptides, and analogues
Amines
0.00 0.25 0.50 0.75 1.00
Fraction of total intensity
$[M-H]^-$
$[M+Cl]^-$
$[M]^-$
$[M-H-HPO_3]^-$
$[M-H_2]^-$
Other
IR-MALDESI-Orbitrap
Vitamins and cofactors
Nucleosides, nucleotides, and analogues
Lipids and lipid-like molecules
Carboxylic acids
Carbohydrates
Amino acids, peptides, and analogues
Amines
0.00 0.25 0.50 0.75 1.00
Fraction of total intensity
$[M-H]^-$
$[M+Cl]^-$
$[M]^-$
$[M-H-CO_2]^-$
$[M-H-H_2]^-$
$[M-H_2]^-$
Other
MALDI-FTICR
Vitamins and cofactors
Nucleosides, nucleotides, and analogues
Lipids and lipid-like molecules
Carboxylic acids
Carbohydrates
Amino acids, peptides, and analogues
Amines
0.00 0.25 0.50 0.75 1.00
Fraction of total intensity
$[M-H]^-$
$[M+Cl]^-$
$[M]^-$
$[M-H-CO_2]^-$
$[M-H+H_2]^-$
$[M-H+Matrix]^-$
Other
MALDI-qTOF
Vitamins and cofactors
Nucleosides, nucleotides, and analogues
Lipids and lipid-like molecules
Carboxylic acids
Carbohydrates
Amino acids, peptides, and analogues
Amines
0.00 0.25 0.50 0.75 1.00
Fraction of total intensity
$[M-H]^-$
$[M+Cl]^-$
$[M]^-$
$[M-H-H_2O]^-$
$[M-H-CO_2]^-$
$[M-H-HPO_3]^-$
$[M-H+H_2]^-$
$[M-H_2]^-$
$[M-H+Matrix]^-$
Other
MALDI2-Orbitrap
Vitamins and cofactors
Nucleosides, nucleotides, and analogues
Lipids and lipid-like molecules
Carboxylic acids
Carbohydrates
Amino acids, peptides, and analogues
Amines
0.00 0.25 0.50 0.75 1.00
Fraction of total intensity
$[M-H]^-$
$[M]^-$
$[M-H-H_2O]^-$
$[M-H-NH_3]^-$
$[M-H-CO_2]^-$
$[M-H-HPO_3]^-$
$[M-HPO_3]^-$
$[M-H+Matrix]^-$
$[M+Matrix]^-$
Other

#### Supplementary Figure 16

Examples of signal dilution across several ion species for selected metabolites. Bar height shows relative abundance of each ion relative to the sum signal of all dejected ions of that metabolite.

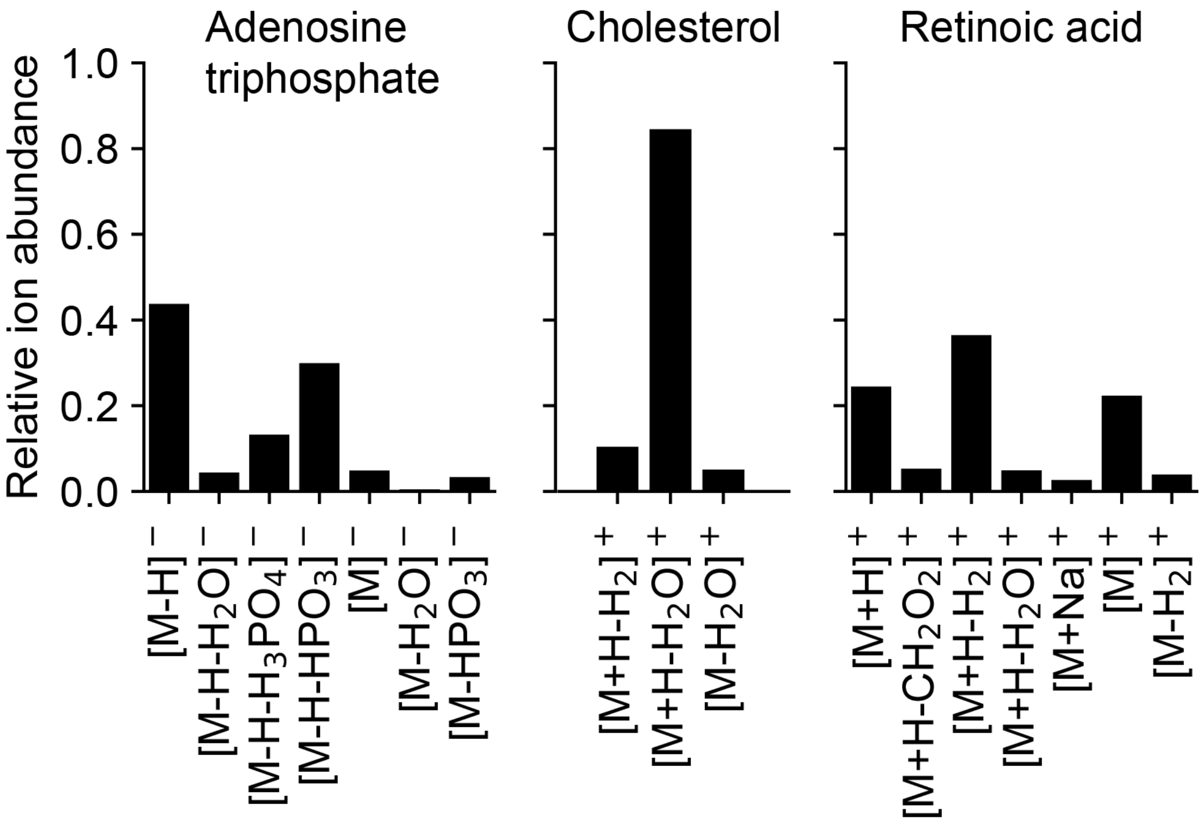
